# McaA and McaB control the dynamic positioning of a bacterial magnetic organelle

**DOI:** 10.1101/2022.04.21.485668

**Authors:** Juan Wan, Caroline L Monteil, Azuma Taoka, Gabriel Ernie, Kieop Park, Matthieu Amor, Elias Taylor-Cornejo, Christopher T Lefevre, Arash Komeili

## Abstract

Magnetotactic bacteria (MTB) are a diverse group of microorganisms that use intracellular chains of ferrimagnetic nanocrystals, produced within their magnetosome organelles, to align and navigate along the geomagnetic field. The cell biological and biochemical properties of magnetosomes make them a powerful model for studying the molecular mechanisms of biomineralization and compartmentalization in bacteria. While several conserved magnetosome formation genes have been described, the evolutionary strategies for their species-specific diversification remain unknown. Here, we demonstrate that the fragmented nature of magnetosome chains in *Magnetospirillum magneticum* AMB-1 is controlled by two genes named *mcaA* and *mcaB*. McaA recognizes the positive curvature of the inner cell membrane while McaB localises to magnetosomes. Along with the MamK actin-like cytoskeleton, they create space for addition of new magnetosomes in between pre-existing magnetosomes. Phylogenetic analyses suggest that McaAB homologs are widespread and may represent an ancient strategy for organelle positioning in MTB.

## Introduction

Cellular compartmentalization results in the formation of different organelles, which need to be positioned correctly to fulfil their specific functions and ensure proper inheritance throughout cell division ^1^. Organelle positioning in eukaryotic cells mainly relies on cytoskeletal and motor proteins ^1^. Many bacteria also produce organelles ^2^, and actively regulate their placement in the cell. For example, the protein-bounded carbon-fixation organelle, the carboxysome, uses the nucleoid as a scaffold with helper proteins that ensure equal distribution in the cell and proper segregation into daughter cells ^3^. Similarly, it has been proposed that the carbon storage polyhydroxybutyrate (PHB) granules associate with nucleoids to mediate segregation during cell division ^4, 5^. A widely studied example of bacterial lipid-bounded organelles is the magnetosome compartment of magnetotactic bacteria (MTB). Magnetosomes mineralize ferrimagnetic nanoparticles composed of magnetite (Fe_3_O_4_) and/or greigite (Fe_3_S_4_) ^2, 6^, which are used as a compass needle for navigation along the geomagnetic field. Magnetic navigation is a common behaviour in diverse organisms, including bacteria, insects, fish, birds, and mammals ^7, 8^. MTB are the simplest and most ancient organism capable of magnetic navigation ^9^ and fossilized magnetosomes chains have been used as robust biosignatures ^10, 11^. Thus, magnetosome production in MTB is an ideal model system for studying mechanisms of organelle positioning, understanding the evolution of magnetic navigation, and connecting the magnetofossil record to the history of life on Earth.

To function as an efficient compass needle, individual magnetosomes need to be arranged into a chain. Various and complex magnetosome chains (single- or multi-stranded, continuous or fragmented) are found in diverse MTB groups ^12,13,14^. The mechanisms leading to distinct chain configurations remain unknown, but may reflect strategies for adaptations to specific biotopes ^10^. The most widely-studied model MTB strains *Magnetospirillum magneticum* AMB-1 (AMB-1) and *Magnetospirillum gryphiswaldense* MSR-1 (MSR-1) are closely related *Alphaproteobacteria* species sharing 96% identity in their 16S rRNA gene sequences ^15^. However, their magnetosome chain organisation strategies are distinct. In AMB-1, magnetosomes containing magnetic crystals and empty magnetosomes are interspersed to form a chain that is fragmented in appearance, extends from pole-to-pole in the cell, and remains stationary during the entire cell cycle ^16^. In contrast, in MSR-1, magnetic crystals are arranged as a continuous chain at the midcell and the divided daughter chains rapidly move from the new poles to the centre of the daughter cells after cell division ^17, 18^. The actin-like protein MamK is conserved in all characterized MTB and forms a cytoskeleton that specifically regulates the stationary or moving behaviours of magnetosome chains in AMB-1 and MSR-1 ^16, 17^. While the overall proteomes of AMB-1 and MSR-1 are on average 66% identical, MamK proteins from the two organisms are 90.8% identical at the amino acid level and *mamK*_AMB-1_ complements the MSR-1 Δ*mamK* mutant ^19^. However, the speed and spatial dynamics of MamK filaments are distinct in each organism ^16, 17^. The acidic protein MamJ is also a key regulator of chain organization. When *mamJ* is deleted in MSR-1, magnetosomes collapse into aggregates in the cell ^20^. However, similar deletions in AMB-1 result in subtle defects with magnetosomes still organised as chains ^21^. Additionally, *mamY* is critical for localizing the chain to the positive curvature of the cell in MSR-1 ^22^ but does not have an impact on chain organization when deleted in AMB-1 ^23^. These observations suggest that unknown genetic elements may be needed for species-specific chain organisation phenotypes.

The genes for magnetosome production and chain assembly in MTB, such as *mamK, mamJ,* and *mamY*, are arranged into magnetosome gene clusters (MGCs) that are often structured as magnetosome gene islands (MAI) ^24^. Unlike MSR-1, AMB-1 contains an extra genomic cluster, termed the magnetotaxis islet (MIS), outside of the MAI region ^25^. The MIS is ∼28kb long and contains 7 magnetosome gene homologs, including a very divergent copy of *mamK* and many genes of unknown function ^25^. The MIS protein, MamK-like, partners with MamK in magnetosome chain formation, but does not contribute to the species-specific chain organisation phenotypes mentioned above ^26^. Whether the other MIS genes are functional and play roles in magnetosome biosynthesis is unclear, especially given the presence of multiple transpose genes and pseudogenes ^25^.

Here, we studied the MIS genomic region and identified two proteins (McaA and McaB) that mediate magnetosome chain assembly in AMB-1. McaA localises to the positively-curved cytoplasmic membrane as a dashed-line even in the absence of magnetosomes whereas McaB associates with magnetosomes. Together, McaA and McaB direct the addition of new magnetosomes to multiple sites between pre-existing magnetosomes to form a fragmented crystal chain. They also influence the dynamics of MamK filaments to control magnetosome positioning during the entire cell cycle. The action of McaA and McaB is sufficient to explain all of the known differences in chain organization between AMB-1 and MSR-1. Broader phylogenetic analysis reveals that McaA and McaB are specific to MTB with distant homologs in the vicinity of MGCs in other MTB. We hypothesize that the MIS is a remnant of an ancient duplication event which paved the way for an alternative chain segregation strategy in AMB-1. This mode of chain segregation may lower the energy requirements for separating magnetic particles at the division septum and eliminating the need for rapidly centring the chain after cell division.

## Results

### MIS genes control the location of and spacing between magnetosomes

To investigate its possible role in magnetosome formation and placement, we deleted the entirety of the MIS from the AMB-1 genome. The deletion’s effect on magnetosome production was assessed by measuring the coefficient of magnetism (Cmag) using a differential spectrophotometric assay that quantifies the ability of MTB to orientate in an external magnetic field ^27^. Unexpectedly, the Cmag values of ΔMIS cultures are much higher than Wild-type (WT) cultures (Fig. 1a), indicating that, as a population, ΔMIS cells better align with the applied external magnetic field. As expected, transmission electron microscopy (TEM) images of WT AMB-1 cultures show that the magnetic crystals are organised into a chain with gaps from cell pole to pole (Fig. 1b). In contrast, the crystals in the ΔMIS strain are organised into a continuous chain at the midcell (Fig. 1b). Analysis of TEM images shows that the number and length of crystals are similar (Supplementary Fig. 1a, b), but the shape factor of crystals (width/length ratio) differs between WT (0.82) and ΔMIS (0.92) strains (Supplementary Fig. 1c, d). These data were collected from strains grown under microaerobic conditions. To ensure that the observed phenotypes were not generated by specific growth conditions, the experiment was repeated under anaerobic conditions and yielded similar results (Supplementary Fig. 2).

**Fig. 1:**
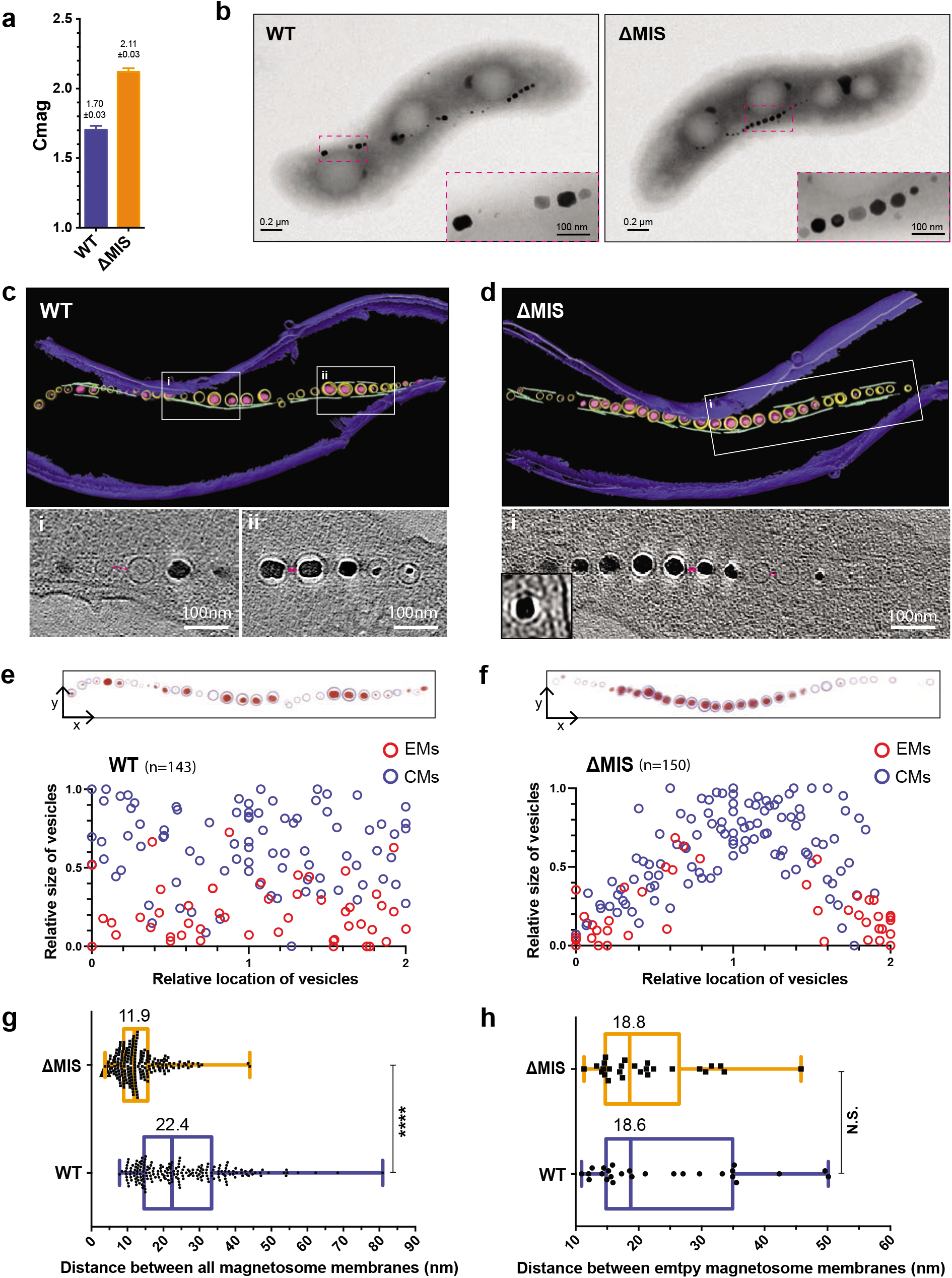
MIS genes contribute to the magnetosome chain assembly. (a) Magnetic response (Cmag) of WT and ΔMIS cultures grown under microaerobic conditions. Each measurement represents the average and standard deviation from three independent growth cultures. (b) TEM micrographs of WT and ΔMIS cells. Insets: magnification of the magnetic crystals in magenta rectangles. (c) and (d) Segmented 3D models (upper panels) and selected area of tomographic slices (lower panels, Box i and ii) showing phenotypes of WT (c) and ΔMIS (d) strains. The outer and inner cell membranes are depicted in dark blue, magnetosome membranes in yellow, magnetic particles in magenta, and magnetosome-associated filaments in green. Full tomograms are shown in Movies S1 and S2. (e) and (f) Relative size and location of magnetosome vesicles in WT and ΔMIS cells, respectively (lower panels). Upper panels are 2D projections of magnetosomes from the 3D models in (c) and (d). The magnetosome membranes are shown in light blue and magnetic particles are shown in red. (g) and (h) Edge-to-Edge distance between all of the magnetosomes (g) and the EMs (h) measured from neighbouring magnetosome membranes. Values represent the median. The box plots show the median (line within box), the first and third quartiles (bottom and top of the box). P-values were calculated by the Mann-Whitney test. No statistically significant difference (P>0.05, N.S.), significant difference (****P < 10^-4^).

Magnetosome biogenesis in WT AMB-1 begins with the invagination of bacterial inner membrane to form empty magnetosomes (EMs), followed by the crystallisation of ferrimagnetic minerals to form crystal-containing magnetosomes (CMs) ^2, 6^. To directly observe the organisation of magnetosome membranes, we imaged WT and ΔMIS cells with whole-cell cryo-electron tomography (cryo-ET) (Fig. 1c, d). Similar to WT ^28^, the magnetosome membranes of ΔMIS mutant are invaginations of the inner membrane (Fig. 1d, lower left corner). We then measured the diameter of magnetosome membranes and the length of crystals. The size distribution of EMs and CMs, as well as the linear relationship between the sizes of crystals and magnetosome membrane diameters are similar between the WT and ΔMIS mutant (Supplementary Fig. 3). Additionally, in both strains, the EMs are significantly smaller than the CMs (Supplementary Fig. 3a). One major phenotypic difference between the two strains is that in WT EMs are present at multiple sites between CMs in the magnetosome chain (Fig. 1c) whereas EMs only localise at both ends of the continuous chain of in the ΔMIS strain (Fig. 1d). Analysing the size and biomineralization status of magnetosome membranes relative to their subcellular position shows that the location of EMs is random along the magnetosome chain in WT but is at both ends of the chain in ΔMIS cells (Fig.1 e, f). These results together suggest that MIS genes control the location of magnetosomes but not the size of magnetosome membranes.

Based on the different sizes and locations of magnetosomes, we hypothesised that the newly-made magnetosomes are added at multiple internal sites of the chain in WT, but only added at the ends of the chain in ΔMIS. To test this hypothesis, we designed a pulse-chase experiment to label and follow a marker protein that incorporates into magnetosomes at the early steps of membrane invagination (Fig. 2a). We examined the magnetosome marker proteins MamI and MmsF ^29,30,31^. Newly-synthesized MmsF proteins incorporate into both the new and old magnetosomes (see more details in supplementary results and supplementary Fig. 4), indicating that it is not suitable for the pulse-chase experiment.

**Fig. 2:**
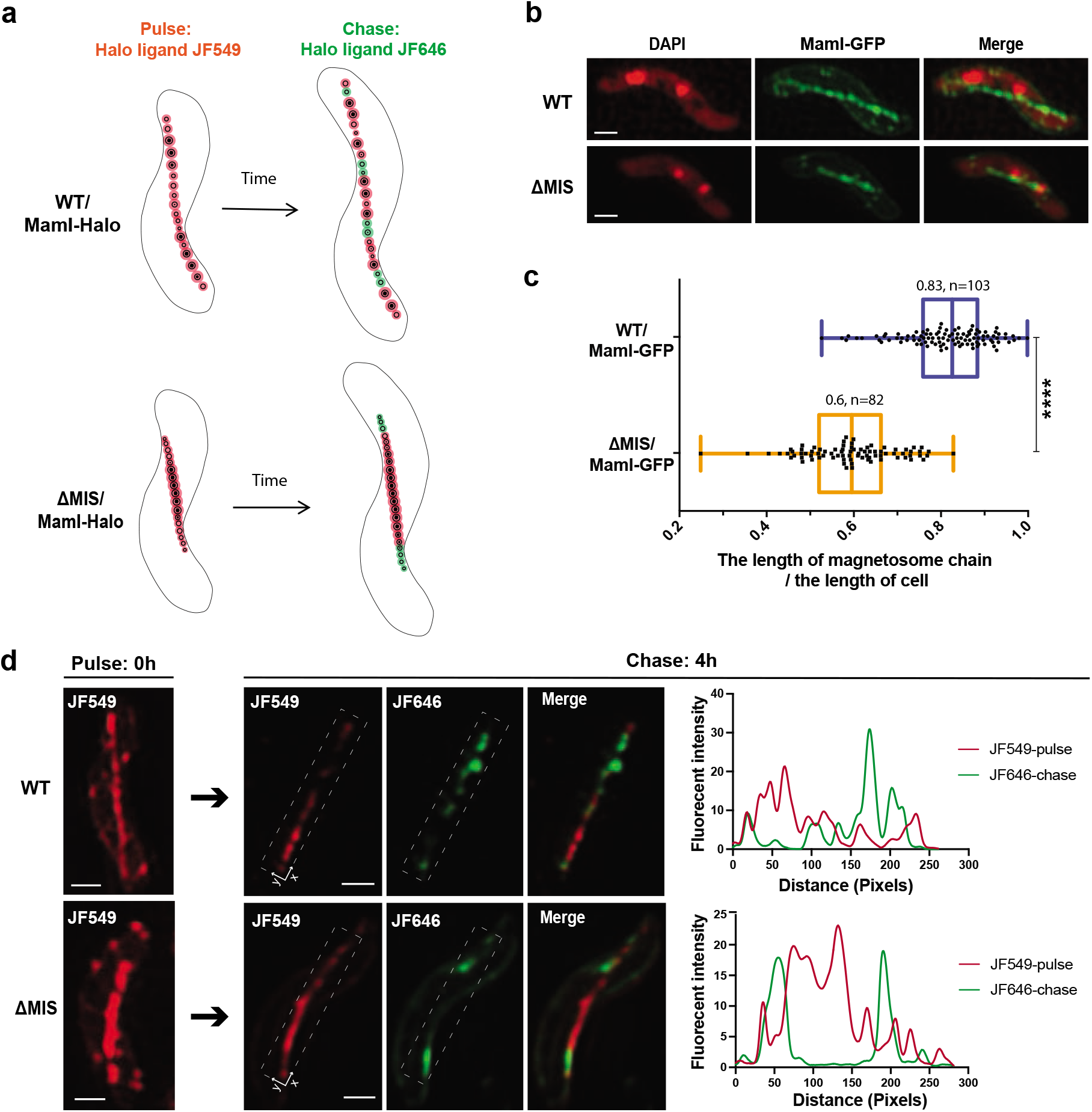
MIS genes determine the location of newly formed magnetosomes in the chain. (a) Model of the pulse-chase experiment shows how the newly-formed EMs are added to the magnetosome chains in WT and ΔMIS. (b) SIM micrographs of WT and ΔMIS cells expressing MamI-GFP under standard growth conditions. The DAPI staining is shown in false-colour red, MamI-GFP is shown in green. (c) Quantification of the length of magnetosome chain versus the length of cell in WT and ΔMIS strains. (d) Left and middle: SIM micrographs from the pulse-chase experiments with MamI-Halo fusion protein for analysing the addition of newly-formed magnetosomes in WT and ΔMIS. The JF549 staining is shown in red, and the JF646 staining is shown in green. Right: fluorescent intensity map of the dashed rectangular area on the SIM micrographs. Scale bars are 0.5 µm in (b) and (d).

The transmembrane protein MamI is needed for EM invagination from the inner membrane of AMB-1 ^26, 31^. We first checked the localization of MamI-GFP in WT and ΔMIS. Structured illumination fluorescent microscopy (SIM) imaging shows MamI-GFP localises as a continuous line from cell pole to pole in WT AMB-1 (Fig. 2b), indicating localization to both the EMs and CMs. In ΔMIS, MamI-GFP only localises in the middle of the cells in a pattern reminiscent of magnetosome organisation as seen in cryo-ET images (Fig. 2b). We then performed pulse-chase experiments using MamI-Halo. The Halo-ligand JF549 was used as the pulse to mark old magnetosomes and the JF646 ligand was chased in to identify the newly-made magnetosomes (Fig. 2a). JF646 signals do not colocalise with JF549 signals in WT AMB-1 (Fig. 2d and Supplementary Fig. 5a), indicating newly-synthesized MamI proteins are only added to the newly-made magnetosomes. Quantitative analysis shows very low colocalization coefficients of the pulse and chase signals in WT and ΔMIS cells (Supplementary Table 1). As expected, JF549-marked old magnetosomes display gaps, which are filled with the JF646-marked newly-made magnetosomes in WT AMB-1 (Fig. 2d and Supplementary Fig. 5a). Conversely, the JF549-marked old magnetosomes still mainly show a continuous chain at the midcell of ΔMIS, and the JF646-marked newly-made magnetosomes localise at both ends of the chain (Fig. 2d and Supplementary Fig. 5a). Together, these results confirm our hypothesis that the varying chain phenotypes between WT and ΔMIS strains are in part due to changes in the location where new magnetosomes are added.

Additionally, we found that the length ratio between the MamI-GFP marked magnetosome chain and the cell body is significantly larger in WT than in ΔMIS (Fig. 2c). As mentioned above, the number of crystals in WT and ΔMIS cells is similar, indicating the distance between the magnetosomes might be different in these two strains. We therefore measured the edge-to-edge magnetosomes distance and found that the distance between all magnetosomes in WT is about twice as long as in the ΔMIS strain (Fig. 1g), while the distance between EMs in these two strains is similar (Fig. 1h), indicating the difference is mainly due to the distance between CMs.

To summarise, MIS genes control the shape of crystals, the distance between CMs, and the location for the addition of newly-made magnetosomes leading to the characteristic pattern of chain organisation in AMB-1.

### Comprehensive dissection of the MIS new chain organization factors

To identify the key genes that control magnetosome positioning, we conducted conventional recombination mutagenesis to create unmarked deletions of selected segments in the MIS (Fig. 3a). We first deleted large domains (LD1 and LD2) to narrow down the region of interest in LD1 (Fig. 3b, c and Supplementary results). We then generated small islet region deletions (ΔiR1, ΔiR2, ΔiR3, and ΔiR4) of LD1 to pinpoint specific genes involved in chain organisation. Genes in the iR2 region control magnetosome positioning, while those in iR3 contribute to crystal shape control (Fig. 3b, d and Supplementary results). iR2 contains a small putative operon with two hypothetical genes (*amb_RS23835* and *amb_RS24750*), which we have named **<u>m</u>**agnetosome **<u>c</u>**hain **<u>a</u>**ssembly genes A and B (*mcaA* and *mcaB*) (Fig. 3a). The reference genome in NCBI shows the iR2 region includes a third transposase gene (*amb_RS23840*) (Fig. 3a), which does not exist in our lab strain. We then deleted these two genes individually. Both Δ*mcaA* and Δ*mcaB* strains have dramatically higher Cmag compared to WT and are similar to ΔMIS (Fig. 3b). Similar to the ΔMIS mutant, Δ*mcaA* and Δ*mcaB* strains contain continuous crystal chains in the midcell region when viewed by TEM (Fig. 3d), indicating that both play essential roles in magnetosome positioning.

**Fig. 3:**
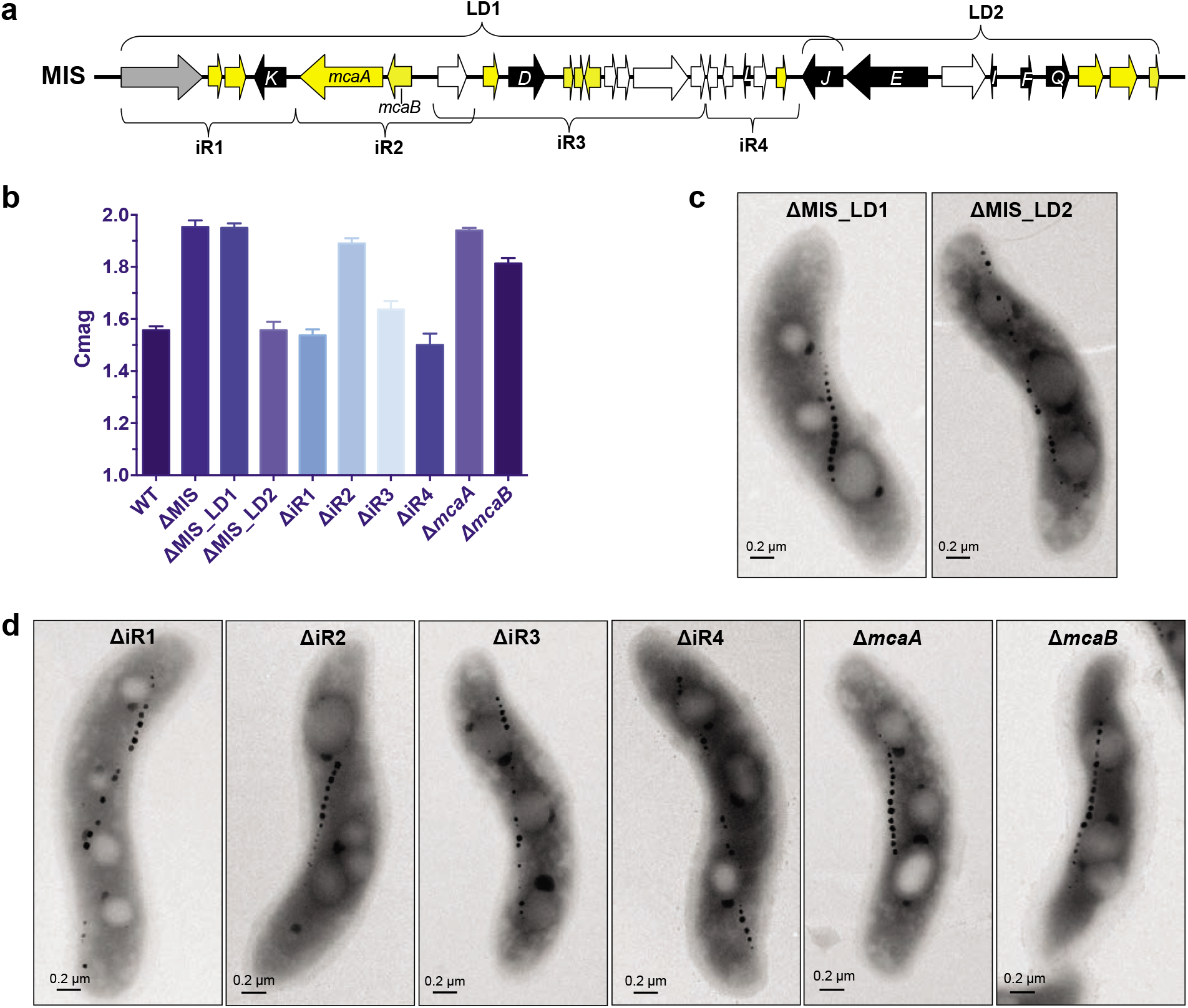
Comprehensive genetic dissection of the MIS identified *mcaA* and *mcaB* as chain assembly factors. (a) Schematic depicting the MIS region of AMB-1, including predicted magnetosome gene homologs (black), a phage-associated protein (grey), transposases (white), and hypothetical genes (yellow). (b) Cmag of WT and different mutants in the MIS region. Each measurement represents the average and standard deviation from three independent growth cultures. (c) TEM micrographs of ΔMIS_LD1 and ΔMIS_LD2 cells. (d) TEM micrographs of ΔiR1-ΔiR4, Δ*mcaA*, and Δ*mcaB* cells.

### McaA localises to the positively-curved cytoplasmic membrane as a dashed line

We interrogated the localization of McaA and McaB in order to understand their role in controlling magnetosome positioning. McaA is predicted to contain a signal peptide, followed by a periplasmic von Willebrand factor type A (VWA) domain, a transmembrane (TM) domain, and a cytoplasmic C-terminus (Fig. 4a and Supplementary results). However, in some bioinformatic predictions, the signal peptide region is predicted to be a TM domain with the N-terminus facing the cytoplasm (Supplementary Table 2). Using GFP fusions to either end of the protein, we predict that the C-terminus of McaA faces the cytoplasm (See more details in Supplementary results). SIM images show that, when expressed in WT AMB-1, McaA-GFP localises in a dashed-line pattern distributed along the positive inner curvature of the cell, forming a line covering the shortest distance from cell pole to pole (Fig. 4b). Since magnetosomes are also located to the positive inner curvature of AMB-1 cell^22^, we wondered if McaA is associated with magnetosomes. To test this, we expressed McaA-GFP in different genetic backgrounds and growth conditions, including ΔMIS and ΔiR2 mutants which contain continuous crystal chains, WT under iron starvation where only EMs are present, and the ΔMAIΔMIS mutant that is incapable of magnetosome production. McaA-GFP localises as a dashed line in all of the above strains and conditions (Fig. 4b), indicating that the association of McaA with the positive curvature of cytoplasmic membrane and its dashed-line localization are independent of magnetosome membrane formation, magnetite production, and chain organisation.

**Fig. 4:**
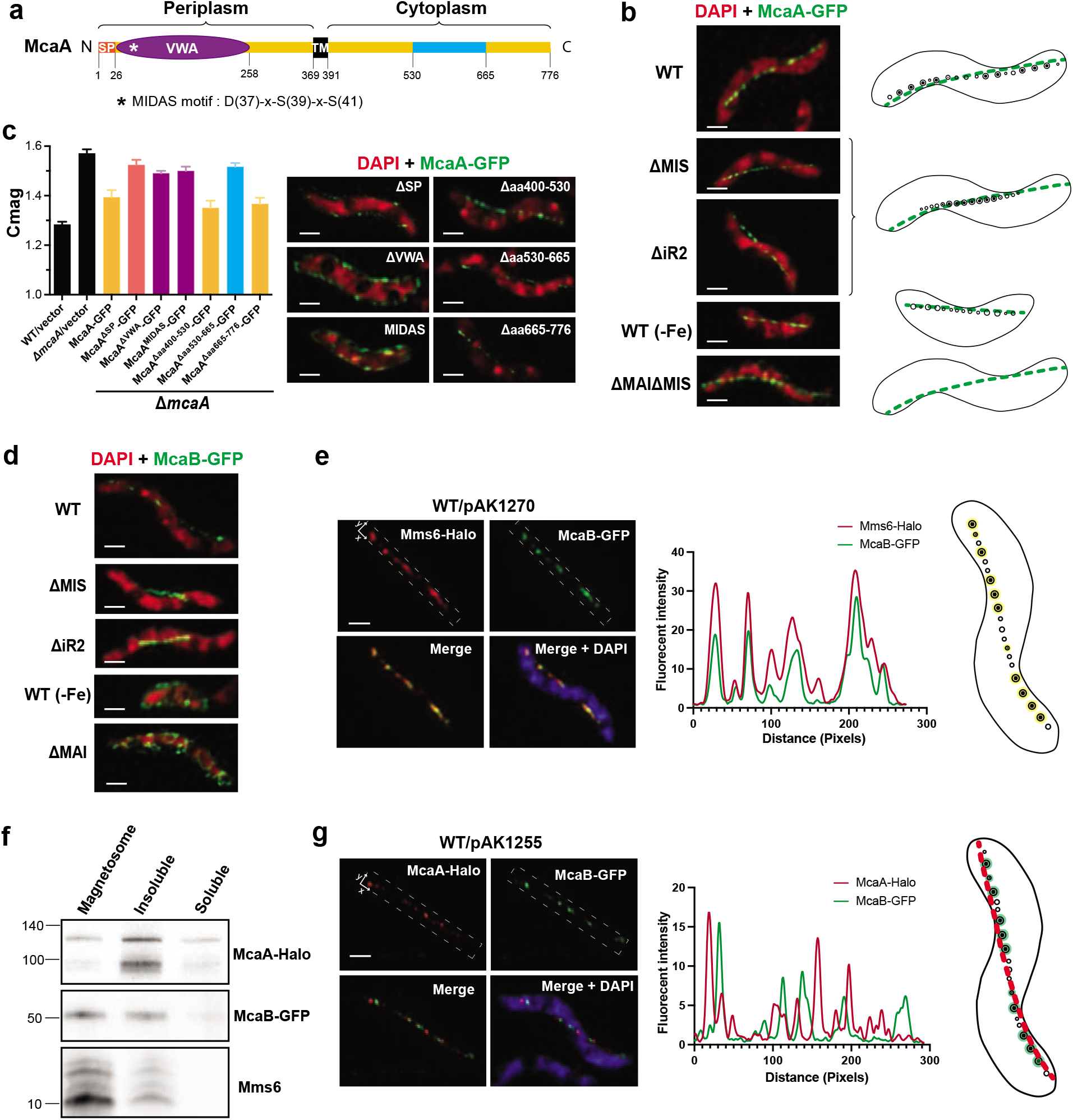
Localization of McaA and McaB. (a)Predicted secondary structure and topology of McaA. SP, signal peptide. VWA, von Willebrand factor type A domain. TM, transmembrane domain. (b) SIM micrographs of cells expressing McaA-GFP (left) and models of magnetosome production (right) in WT and different genetic backgrounds or growth conditions. (c) Cmag (left) and SIM micrographs (right) of mutated McaA-GFP expressed in Δ*mcaA* cells. (d) SIM micrographs show the localization of McaB-GFP in WT and different genetic backgrounds. From (b) to (d), DAPI staining is shown in false-colour red and the GFP fusion proteins are shown in green. (e) Left: SIM micrographs show the localization of Mms6-Halo and McaB-GFP in WT. Middle: fluorescent intensity map of the dashed rectangular area on the SIM micrographs. Right: model of the colocalization (yellow) of Mms6 and McaB at CMs. (f) Western blotting shows McaA and McaB are enriched in different cellular fractionations. (g) Left: SIM micrographs show the localization of McaA-Halo and McaB-GFP in WT. Middle: fluorescent intensity map of the dashed rectangular area on the SIM micrographs. Right: model of the association of McaA (red) and McaB (green) with magnetosomes. In (e) and (g), the DAPI staining is shown in blue, the JF549-stained Halo proteins are shown in red, GFP fusion proteins are shown in green. Scale bars: 0.5 µm.

Using a series of truncations, we found that the N-terminus of McaA (including the predicted signal peptide and the VWA domain) is essential for its localization and magnetosome positioning. In contrast, the C-terminus conserved region (aa 530-665) is not essential for McaA localization but important for magnetosome positioning (Fig. 4c and Supplementary Fig. 8). The VWA domain is commonly involved in protein-protein interactions, largely *via* a noncontiguous metal ion-binding motif (DXSXS) called metal ion-dependent adhesion site (MIDAS) ^32, 33^. McaA VWA domain contains an intact MIDAS motif. We investigated whether this MIDAS motif plays an important role using site-directed mutagenesis. McaA^MIDAS^ mutant cannot complement Δ*mcaA* and is evenly distribute within the cytoplasmic membrane (Fig. 4c and Supplementary Fig. 8), highlighting the important role of the MIDAS motif and divalent cations in the localization and function of McaA.

### McaB associates with crystal-containing magnetosomes

McaB is predicted to contain one TM domain that is close to the N-terminus, which is mostly facing the periplasm (Supplementary Fig. 9a). We confirmed the cytoplasmic location of C-terminus McaB through the fluorescent signal of McaB-GFP (Fig. 4d and Supplementary results). SIM images show that McaB-GFP forms a dotted line from cell pole to pole along the positive inner curvature of WT AMB-1 cells, whereas it exhibits a continuous line in the middle of the ΔMIS and ΔiR2 cells (Fig. 4d and Supplementary Fig. 9b), indicating McaB might be associated with magnetosomes. Interestingly, McaB-GFP is not present at the magnetosome chain when WT cells are grown under low iron conditions that prevent magnetite production (Fig. 4d and Supplementary Fig. 9b), indicating that it does not localise to the EMs. These results together show that McaB might be specifically associated with CMs, resembling the localization pattern of the magnetosome protein Mms6 ^34^. Thus, we co-expressed McaB-GFP and Mms6-Halo in the same AMB-1 cell, and the SIM images, as well as quantitative determination of colocalization coefficients (Supplementary Table 1), show that the two proteins colocalise (Fig. 4e), further confirming the association of McaB with CMs.

We also examined the localization of McaA and McaB using cellular fractionation and immunoblotting analysis. McaA is mainly detected in the insoluble portion which includes the cytoplasmic membranes, while similar to Mms6, McaB is mainly detected in the magnetosome fraction (Fig. 4f). Thus, biochemical fractionation experiments confirm the association of McaA with cytoplasmic membrane and McaB with magnetosome membranes.

### McaA and McaB coordinate magnetosome positioning

To explore the relationship between McaA and McaB, we co-expressed McaA-Halo and MacB-GFP in WT AMB-1 cells (see more details in supplementary results). Interestingly, SIM images show that McaB localises within the gaps of dashed McaA, indicating McaB-marked CMs are located in the gaps of dashed McaA (Fig. 4g and Supplementary Fig. 10d). Accordingly, quantitative analysis shows low colocalization coefficients for McaA and McaB signals in WT AMB-1 cells (Supplementary Table 1). Bacterial adenylate cyclase two-hybrid (BACTH) assays did not show any positive interactions between McaA and McaB (Table 1 and Supplementary Fig. 11a), indicating that either the fusion proteins do not interact strongly, are nonfuncional in the context of BACTH, or unknown intermediate proteins are needed to facilitate their interactions.

**Table 1.** The interaction results of BACTH.

|  | Zip-T25 | T25-McaA | McaA-T25 | T25-McaB | McaB-T25 | T25-MamK | MamK-T25 | T25-MamJ | T25-MamY | MamY-T25 |
| --- | --- | --- | --- | --- | --- | --- | --- | --- | --- | --- |
| **Zip-T18** | + | NA | NA | NA | NA | NA | NA | NA | NA | NA |
| **T18-McaA** | NA | - | - | - | - | - | - | - | - | - |
| **McaA-T18** | NA | - | - | - | - | - | - | - | - | - |
| **T18-McaB** | NA | - | - | - | - | - | - | - | - | - |
| **McaB-T18** | NA | - | - | - | - | - | - | - | - | - |
| **T18-MamK** | NA | - | - | - | - | + | + | NA | NA | NA |
| **MamK-T18** | NA | - | - | - | - | + | + | NA | NA | NA |
| **MamJ-T18** | NA | - | - | - | - | NA | NA | - | NA | NA |
| **T18-MamY** | NA | - | - | - | - | NA | NA | NA | - | - |
| **MamY-T18** | NA | - | - | - | - | NA | NA | NA | - | + |

NA, No analysis was performed. -, Negative Interaction (White Colonies), +, Positive Interaction (Blue Colonies).

We next investigated whether McaA and McaB directly affect the positioning of EMs when CMs are not produced. We grew WT, ΔMIS, and ΔiR2 strains expressing MamI-GFP under iron starvation conditions. SIM images show that MamI-marked EMs are located continuously in the midcell of all three strains (Fig. 5a), indicating that McaAB do not participate in the chain organisation under low iron conditions. We then examined the dynamics of magnetosome chain organisation as WT and cells missing *mcaAB* transitioned from low to high iron conditions to trigger magnetite production in EMs (Fig. 5b). We performed pulse-chase experiments using MamI-Halo with cells growing from iron starvation (pulse with JF549) to standard iron growth conditions (chase with JF646). As expected, the pulse experiments show a continuous chain of EMs in the middle of all WT and *mcaAB* deficient cells under iron starvation conditions (Fig. 5c and Supplementary Fig. 5b). After iron addition to the growth medium, we observed the formation of gaps between JF549-marked old magnetosomes in WT, but not in ΔMIS or ΔiR2 cells (Fig. 5c and Supplementary Fig. 5b). In addition, the JF646-marked newly-made EMs filled the gaps between older magnetosomes in WT but were only added at both ends of the chain in ΔMIS and ΔiR2 (Fig. 5c and Supplementary Fig. 5b). Accordingly, quantitative analysis shows a low colocalization coefficient of the pulse and chase signals in WT, ΔMIS, and ΔiR2 cells (Supplementary Table 1). Together, these results support the hypothesis that McaA serves as a landmark on the positively-curved inner membrane and coordinates with McaB to control the location and spacing between CMs, allowing the addition of newly-made EMs to multiple sites between pre-existing magnetosomes in the chain of WT AMB-1, which forms the fragmented crystal chain.

**Fig. 5:**
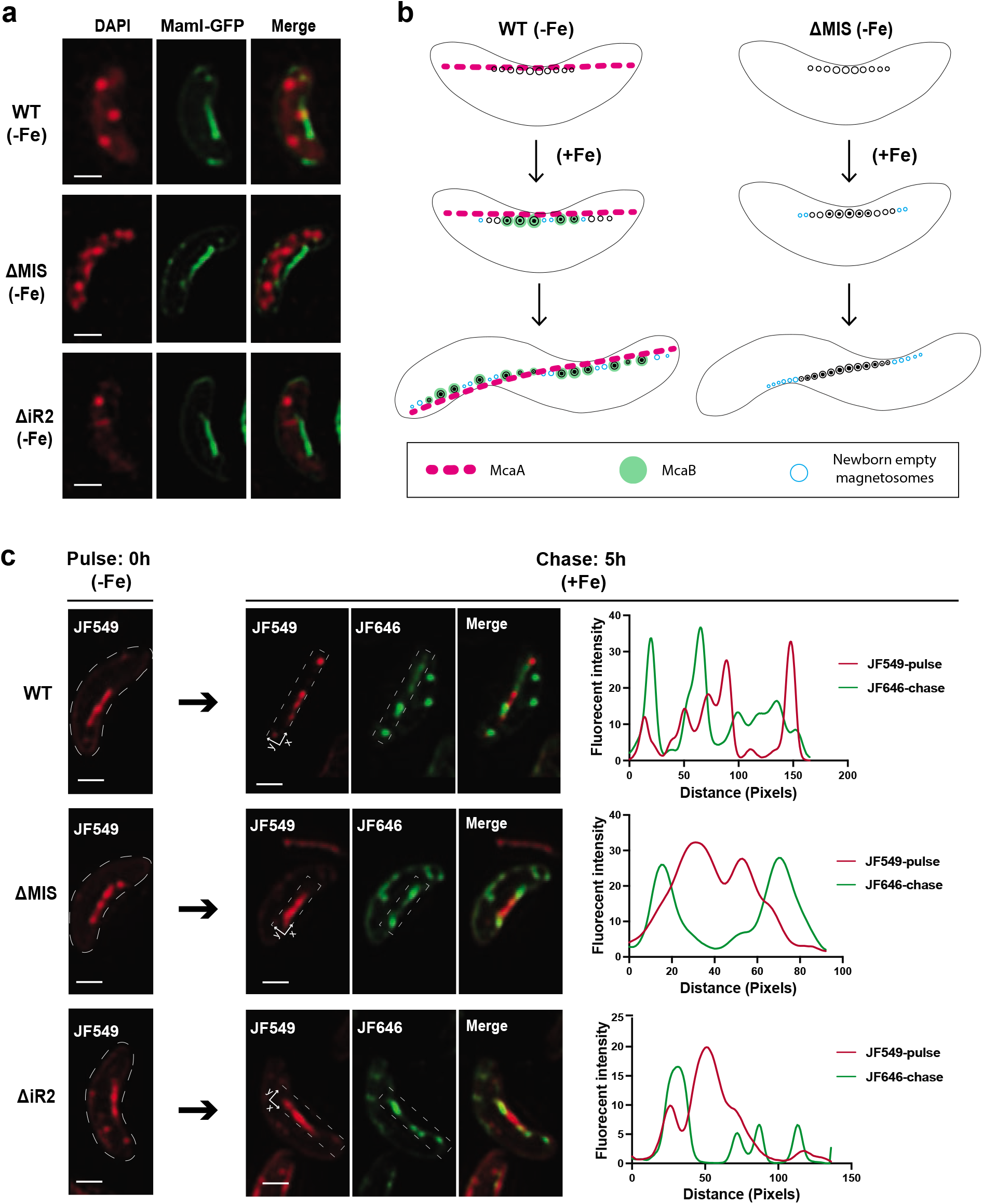
Addition of newly formed magnetosomes to the chain from iron starvation to normal iron growth conditions. (a) Representative SIM micrographs of WT, ΔMIS, and ΔiR2 cells expressing MamI-GFP under iron starvation growth conditions. The DAPI staining is shown in false-colour red, MamI-GFP is shown in green. (b) Model of how McaA and McaB coordinate to control the addition of newly formed magnetosomes in WT and ΔMIS cells. A continuous chain of EMs is generated in the middle of both WT and McaAB deficient cells under iron starvation conditions. After iron is added to the medium, magnetic crystals are mineralized in the existing EMs ^20, 22^. Once EMs become CMs, the McaB is recruited to CMs and help locating CMs to the gaps of dashed McaA, thereby increasing the distance between CMs, which allows for the addition of newly formed EMs inbetween CMs in WT AMB-1. In the absence of McaAB, CMs are located close to each other, and newly formed EMs are added at both ends of the chain. (c) Left and middle: SIM micrographs from pulse-chase experiments with MamI-Halo fusion protein for analysing the addition of newly formed magnetosomes in different AMB-1 genetic backgrounds from iron starvation to normal iron growth conditions. Right: fluorescent intensity map of the dashed rectangular area on the SIM micrographs. JF549 staining is shown in red, and JF646 staining is shown in green. Scale bars are 0.5 µm in (a) and (c).

### McaA contributes to the differences of *mamJ* and *mamY* deletions between AMB-1 and MSR-1

As mentioned above, the phetotypes of *mamJ* and *mamY* deletion mutants in AMB-1 and MSR-1 are distinct. MamJ is proposed as a linker to attach MamK filaments to magnetosomes and its deletion in MSR-1 causes the magnetosome chain to collapse and form an aggregate ^20^. In contrast, the deletion of *mamJ* and its homolog *limJ* in AMB-1 still shows a magnetosome chain with some minor structural defects ^21^. However, deletion of the entire MIS in a 11*mamJ*11*limJ* strain causes a dramatic chain collapse phentype resembling those of MSR-111*mamJ* mutant ^10, 20^. MIS contains a second *mamJ* homolog called *mamJ-like* and our deletion analysis shows that it does not contribute to chain maintenance (Fig. 3a, Fig. 6a, c and more details in Supplementary results). To figure out the specific genes, we generated large domain and small region deletion mutants of the MIS in a Δ*mamJ*Δ*limJ* background. Cmag and TEM images show that *mcaA* is the specific gene that prevents magnetosome aggregation in the Δ*mamJ*Δ*limJ* strain (Fig. 6a, c and Supplementary Fig. 14).

**Fig. 6:**
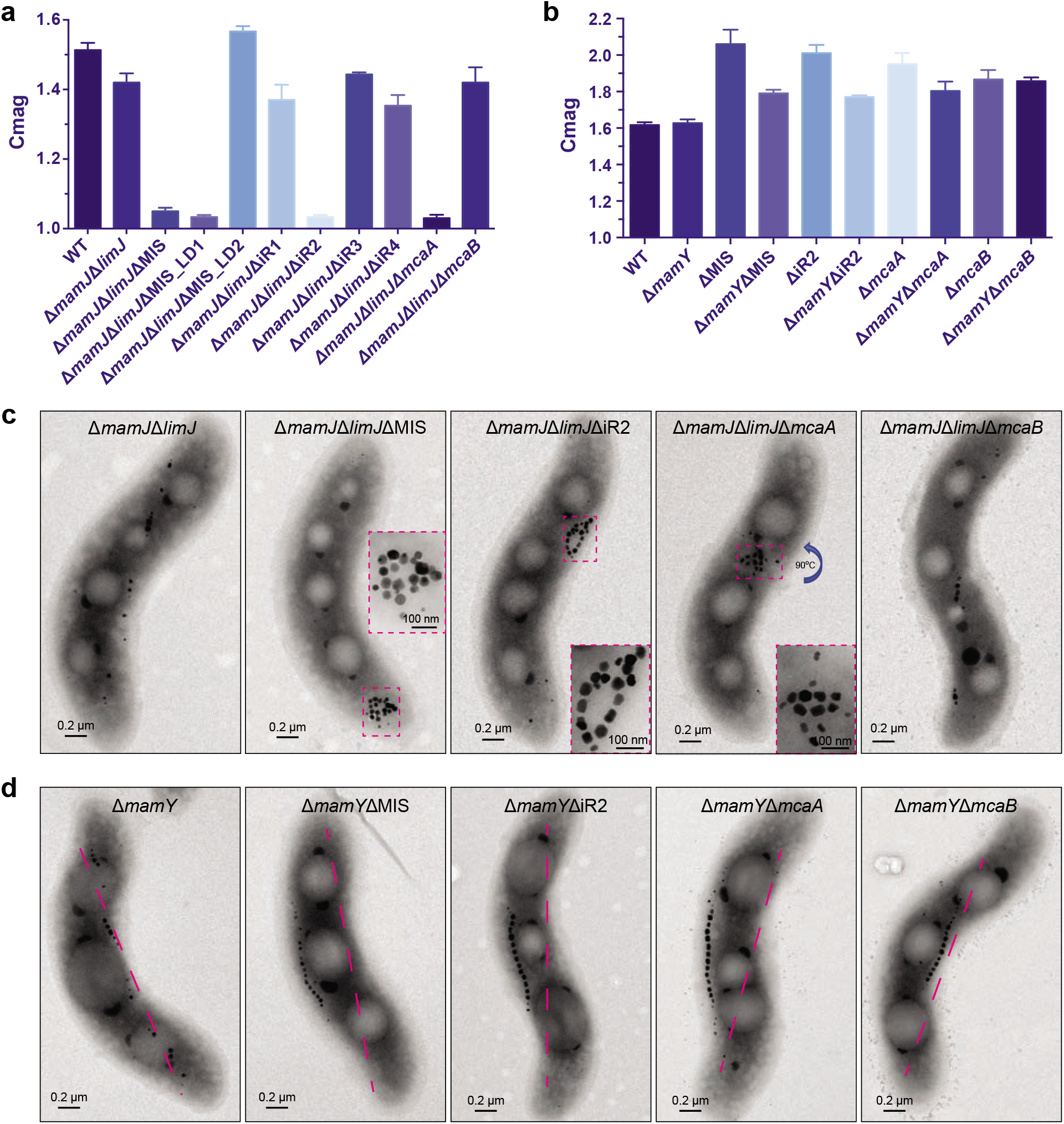
McaA prevents magnetosome chain aggregation and directs the chain to the positively curved regions of the cytoplasmic membrane. (a) and (b) Cmag of WT and different mutants in the Δ*mamJ*Δ*limJ* (a) and Δ*mamY* (b) backgrounds. Each measurement represents the average and standard deviation from three independent growth cultures. (c) TEM micrographs of Δ*mamJ*Δ*limJ*, Δ*mamJ*Δ*limJ*ΔMIS, Δ*mamJ*Δ*limJ*ΔiR2, Δ*mamJ*Δ*limJ*Δ*mcaA*, and Δ*mamJ*Δ*limJ*Δ*mcaB* cells. (d) TEM micrographs of Δ*mamY*, Δ*mamY*ΔMIS, Δ*mamY*ΔiR2, Δ*mamY*Δ*mcaA*, and Δ*mamY*Δ*mcaB* cells.

MamY is a membrane protein that directs magnetosomes to the positively-curved inner membrane in MSR-1, thus aligning the magnetosome chain to the motility axis within a helical cell ^22^. When *mamY* is deleted in MSR-1, the magnetosome chain is no longer restricted to the positively-curved regions of the membrane and can also be found at the negatively-curved membrane leading to a much lower Cmag compared to WT ^22^. Surprisingly, when *mamY* is deleted in AMB-1, the Cmag is similar to WT (Fig. 6b), and the magnetosome chain still localises to the positively-curved membrane (Fig. 6d), indicating there might be other proteins that are functionally redundant to MamY in AMB-1. A Δ*mamY*ΔMIS mutant of AMB-1 has a much lower Cmag than ΔMIS and produces magnetosome chains that localise to both positively- and negatively-curved cell membranes (Fig. 6b, d). Further deletion mutagenesis shows that McaA helps magnetosomes localise to the positive-curved membrane when MamY is lost in AMB-1 (Fig. 6b, d).

Despite their genetic interactions, BACTH analysis does not show any direct interactions between MamJ and McaA, -B or between MamY and McaA, -B (Table 1 and Supplementary Fig. 11b-e). Nevertheless, our results indicate that the activity of McaA accounts for the distinct phenotypes of *mamJ* or *mamY* deletion mutants between AMB-1 and MSR-1.

### McaA and McaB control the dynamic positioning of magnetosomes by influencing the MamK filaments

In addition to the appearance of the chain, the dynamic movements and positioning of the chain differs between AMB-1 and MSR-1 ^16, 17^. We reasoned that the McaAB system may contribute to the different dynamic chain positioning in these two strains. Using highly inclined and laminated optical sheet (HILO) microscopy, we performed live-cell imaging analysis to follow the dynamics of Mms6-GFP labelled magnetosomes during cell division in WT and McaAB deficient cells.

In WT AMB-1, magnetosomes are in static, spotty positions during cell division as indicated by the parallel lines in the kymographs of GFP fluorescence (Fig. 7a and Supplementary Movie 1), whereas every ΔMIS and Δ*mcaA* cells shows dynamic magnetosome chain segregation after cell division (Fig. 7b, c and Supplementary Movie 2, 3). In ΔMIS and Δ*mcaA*, magnetosomes are positioned at the midcell until the cell divides. After cytokinesis, magnetosomes are moved synchronously toward the centres of both daughter cells. Magnetosome migration to the middle of the daughter cells was completed within about 1 hour after cell division. Magnetosome chain displacement velocity in Δ*mcaA* cells was about 20 nm/min, in line with what has been reported previously for MSR-1 ^17^. Supplementary Movies 5 and 6 are long time-lapse videos of the ΔMIS and Δ*mcaA* cells, in which the magnetosomes are stably positioned at the middle of the daughter cells during the entire cell cycle after the migration of the magnetosomes. In contrast, in Δ*mcaB* cells, magnetosome displacements are incomplete (Fig. 7d and Supplementary Movie 4, 7 (short and long time-lapse)). Magnetosomes do not move synchronously toward the middle of the cell and are randomly positioned in the cell. In other words, these results show that McaB does not impact daughter chain positioning in Δ*mcaA* strain, while McaA impedes daughter chain positioning in Δ*mcaB* strain, indicating McaA might have extra functions in magnetosome chain positioning during cell division.

**Fig. 7:**
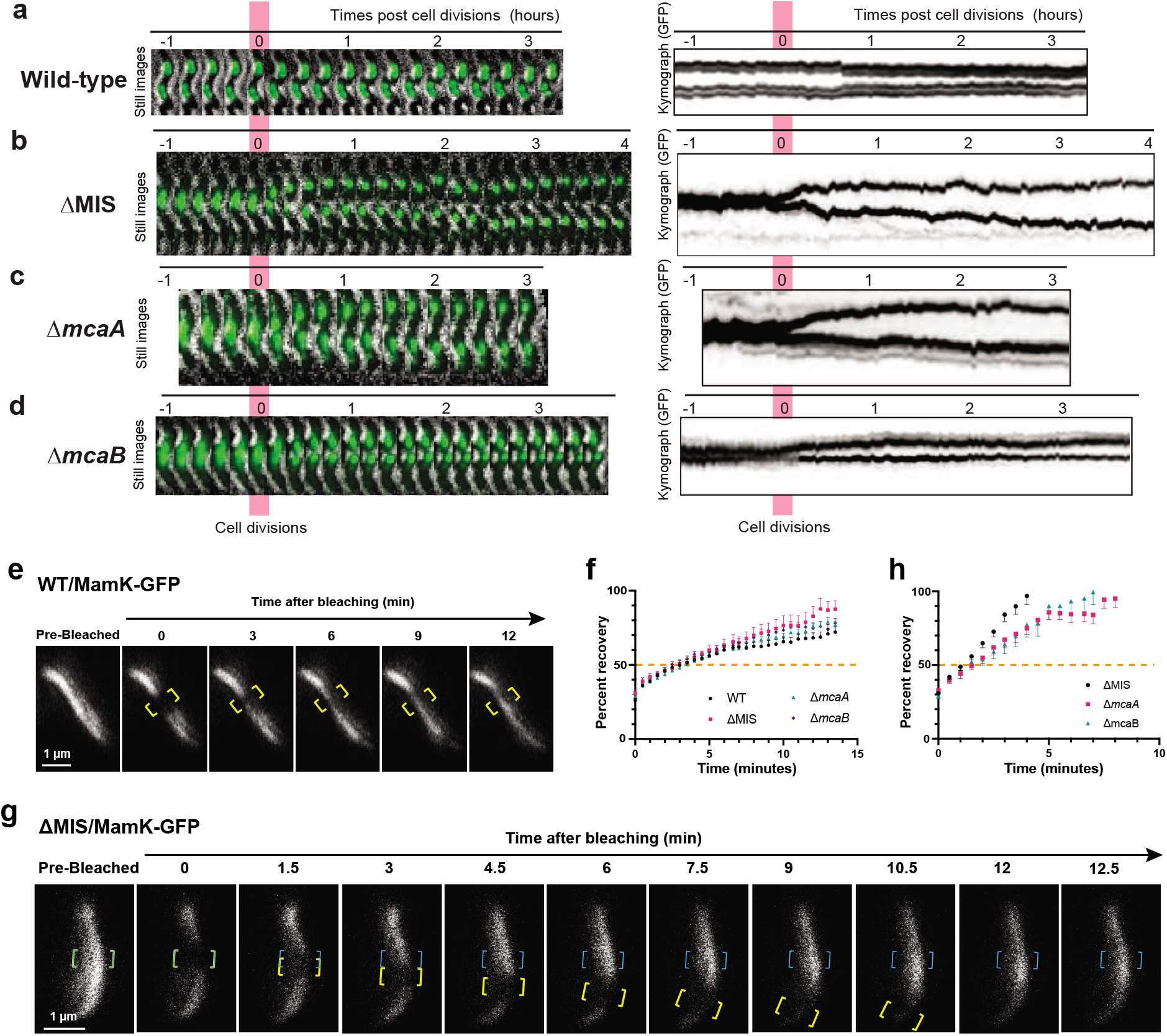
McaAB control the segregation of magnetosomes and dynamics of MamK filaments. (a) to (d) Effects of *mcaA* and *mcaB* deletions on magnetosomes segregation. Live-cell time-lapse imaging of magnetosome segregation in wild-type (a), ΔMIS (b), Δ*mcaA* (c), and Δ*mcaB* (d) cells during cell division. Magnetosomes are marked with Mms6-GFP. Left: GFP fluorescence and bright field merged time-lapse still images. Right: kymographs of Mms6-GFP signals in maximum projection. (e) A FRAP experiment time course with WT AMB-1 expressing MamK-GFP. Yellow brackets indicate the portion of the MamK-GFP filament designated for photobleaching. (g) A FRAP experiment time course with ΔMIS expressing MamK-GFP where the bleached area moved from its original position toward the cell pole. Yellow and blue brackets indicate the portion of the MamK-GFP filament designated for photobleaching. Blue brackets indicate the original bleaching area, and yellow brackets track the movement of the bleached area. The MamK-GFP is shown in false-colour white in (e) and (g). (f) and (h) Normalized (average mean and standard error of mean [SEM]) percent recovery of each strain’s recovering cells with non-moving (f) and moving (h) bleached area. The 50% mark is noted with a dashed orange line.

It has been shown that the dynamics of MamK filaments is essential for magnetosome chain positioning in AMB-1 and MSR-1 ^16, 17^. To test whether the McaAB system influences the dynamics of MamK filaments, we performed fluorescence recovery after photobleaching (FRAP) assays on MamK-GFP filaments in WT and McaAB deficient cells. GFP-tagged MamK filaments localise as even thin lines from cell pole to pole in both WT and McaAB deficient AMB-1 strains. During FRAP experiments, sections of GFP-tagged MamK filaments are irreversibly photobleached and the recovery of fluorescence in the bleached area is tracked over time (Fig. 7e). The half-life (t_1/2_) of recovery represents the time point at which 50% of the fluorescence intensity returns to the bleached region relative to the whole filament at that same time point. The bleached area does not move in WT (Fig. 7e and Supplementary Fig. 15a), but it moves in a fraction of ΔMIS, Δ*mcaA*, and Δ*mcaB* cells (Fig. 7g, Supplementary Fig. 15b, and Supplementary Table 3). The t_1/2_ fluorescence recovery is similar in WT and McaAB deficient cells containing an immotile bleached spot (Fig. 7f and Supplementary Fig. 15a). For the McaAB deficient cells with a moving bleached spot, the t_1/2_ fluorescence recovery of the original bleached area is similar but much faster than in the cells containing an immotile bleached spot (Fig. 7h), indicating a similar moving speed of the bleached area. BACTH analysis did show MamK self-interactions but did not show any interactions between McaA, -B and MamK (Table 1 and Supplementary Fig. 11f, g). Together, these results indicate that McaA, -B influence the dynamics of MamK filaments which, in turn, leads to the AMB-1-specific pattern of magnetosome chain organization.

### *mcaA* and *mcaB* genes are specific to MTB

To understand the evolutionary origins of the *mcaAB* system, we searched for homologs of these two genes in diverse species of MTB. Distant homologs were found in 38 MTB species. All of them belong to MTB strains either with characterized magnetosome chain phenotypes (Fig. 8a, b) or metagenomes obtained from a magnetic enrichment (supplementary Fig. 16 a, b), with the majority affiliated to the *Rhodospirillaceae* family in the *Alphaproteobacteria* class. Based on published reports, most studied MTB contain continuous crystal chains (supplementary dataset 1). However, some species show fragmented crystal chains, including the two *Alphaproteobacteria* species (*Ca*. Terasakiella magnetica PR-1 and *Terasakiella sp*. SH-1) that contain distant *mcaAB* homologs and two *Deltaproteobacteria* species that do not contain *mcaAB* homologs (Fig. 8a, b and supplementary dataset 1), which suggests that different mechanisms may exist for crystal chain fragmentation.

**Fig. 8:**
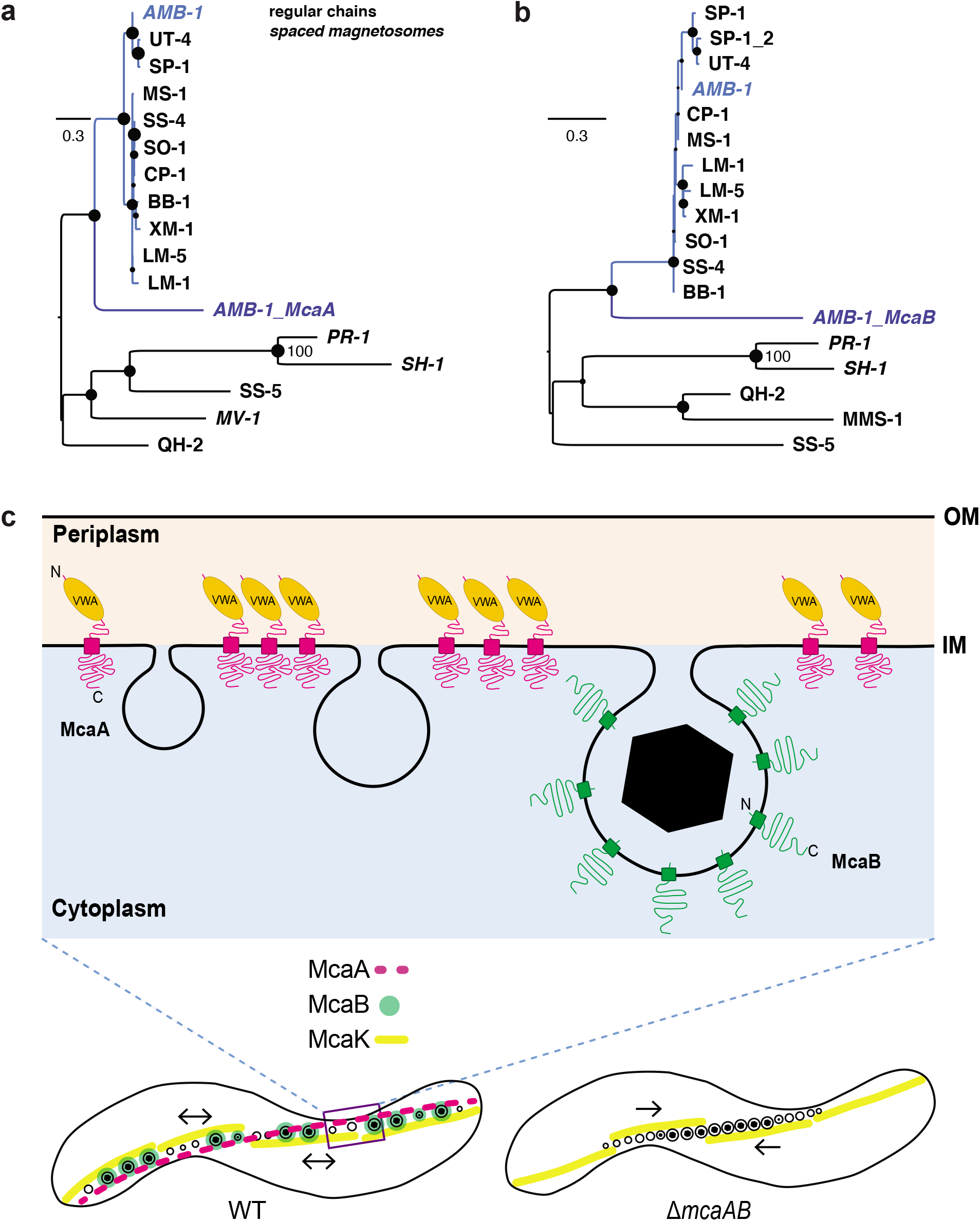
Phylogenetic analysis and model of McaAB-mediated magnetosome chain assembly. (a) and (b) Maximum likelihood trees showing the ancestry of McaA (a) and McaB (b) proteins in relation to their homologs in freshwater magnetotactic *Rhodospirillaceae* (blue clade) and the external groups of other *Proteobacteria*. All strains’ accession numbers are given in supplementary dataset 1. Trees were drawn to scale and branch length refers to the numbers of substitution per site. Robustness of the internal branches is symbolized by a circle whose size is proportional to the bootstrap value estimated from 500 non-parametric replicates. The magnetosome chains in these strains were previously characterized. If magnetosomes are spaced from each other similarly to strain AMB-1, names are in italics. (c) McaA serves as a landmark on the positively curved inner membrane and coordinates with McaB to control the location of CMs to the gap region of dashed-McaA. As a consequence, the neighbouring CMs are separated from each other, which allows the addition of newly formed EMs to multiple sites of the magnetosome chain in WT AMB-1. Alternatively, the CMs are located closely together without McaAB, leaving no space for the addition of newly formed EMs between CMs but at both ends of the magnetosome chain. McaAB also influence the dynamics of MamK filaments to control the dynamic positioning of magnetosomes during the whole cell cycle. OM, outer membrane. IM, inner membrane.

Besides the *mcaAB* genes of the MIS, AMB-1 contains two additional homologs of *mcaA* and *mcaB* with a similar domain architecture (named *mcaA-like* and *mcaB-like* here) that are present close to the MAI. McaA-like (encoded by *amb0908*/*amb_RS04660*) has ∼45% amino acid sequence identity over ∼66% McaA sequence length, while McaB-like (encoded by *amb0907*/*amb_RS24855*) has ∼44% amino acid sequence identity over ∼66% McaB sequence length. We deleted *mcaAB*-like genes in both WT and ΔMIS AMB-1 strains but did not observe any obvious defects or changes in magnetosome formation or chain organisation (supplementary Fig. 16 c, d), indicating *mcaAB*-like genes are not functionally redundant with *mcaAB* genes in AMB-1. Based on comparative genomic analyses, *mcaA* and *mcaB* of AMB-1 form a distinct cluster, while *mcaA-like* and *mcaB-like* of AMB-1 cluster with a second group of homologs in the strains that do not show a fragmented crystal chain phenotype (Fig. 8a, b, clade in light blue), indicating those homologs might have functions more similar to *mcaAB-like* genes in AMB-1.

Molecular phylogenetics indicates that the last common ancestor to Mca proteins of AMB-1 and Mca-like proteins detected in *Magnetospirillum* species and *Ca*. Magneticavibrio boulderlitore LM-1 emerged before the first freshwater magnetotactic *Rhodospirillaceae* (Fig. 8a, b). High protein sequence identity percentages of Mca-like proteins between *Magnetospirillum* strains (85% up to 100%) compared to the average amino acid ID % at the genome scale ^35^ indicates that the *mca-like* genes are under purifying selection. Together these results may indicate that McaAB-mediated crystal chain fragmentation could be a trace of an ancient chain organisation strategy that is lost in most of the modern MTB species. However, the long external branches of McaA and McaB (Fig. 8a, b and supplementary Fig. 16 a, b) suggest a recent acceleration of the evolution of Mca proteins which could be linked with their neofunctionalization in AMB-1.

## Discussion

In this study, we uncovered the mechanisms of a novel magnetosome chain organisation strategy that explains phenotypic differences between closely related MTB. We demonstrated that the fragmented crystal chain organisation in WT AMB-1 strain is not growth condition dependent but genetically controlled by *mcaA* and *mcaB*. In their absence, magnetite crystals form a continuous chain similar to the chain phenotype of MSR-1 ^22^. The McaAB system also contributes to the differences in magnetosome dynamic positioning between AMB-1 and MSR-1 during the entire cell cycle.

Based on our study, we propose a model for McaAB-mediated dynamic positioning of magnetosomes (Fig. 8c). McaA localises to the positive inner curvature of the cell as a dashed line via its N-terminal periplasmic VWA domain. McaA serves as a landmark to regulate the placement and distance of McaB-marked CMs through its cytoplasmic C-terminal domain. As a consequence, neighbouring CMs are separated from each other allowing newly made EMs to be added at multiple locations of the chain. Without McaAB, the CMs are located closely together such that newly made EMs can only be added at the ends of the magnetosome chain. Furthermore, cryo-ET shows that MamK cytoskeleton is composed of short filaments and located along the magnetosome chain in both WT and ΔMIS cells (Fig. 1c, d). Based on FRAP experiments, MamK filaments in WT display local recovery that can be caused by monomer turnover (depolymerization/polymerization), filament sliding, or formation of new filaments. In many McaAB deficient cells, MamK filaments recover and at the same time move across the cell, which might help position the magnetosomes in the midcell (Fig. 8c). We propose that the McaAB system localises the turnover of MamK filaments to allow for new magnetosome addition in between pre-existing magnetosomes in WT AMB-1 (Fig. 8c).

Beyond elucidating an unknown aspect of magnetosome chain formation, our findings raise new questions regarding the cell biological mechanisms of organelle formation and maintenance in bacteria. For instance, how McaA detects the positive curvature of cytoplasmic membrane and localises as a dashed pattern remains a mystery. We confirmed that the localization of McaA is dependent on its VWA domain. Eukaryotic VWA-containing proteins are involved in a wide range of cellular functions, but they share the common feature of being involved in protein-protein interactions, many of which depend on divalent cations coordinated by the MIDAS motif ^32^. Consistently, the MIDAS motif is essential for the location and function of McaA. VWA-domain proteins have been identified in some bacteria and archaea with different functions, but they are not well characterised ^36,37,38^. These data suggest that there might be unknown proteins that partner with McaA to determine its specific localization. In addition to coordinating with McaB to control the fragmented crystal chain assembly, McaA also helps to prevent magnetosome chain aggregation when *mamJ* and its homologs are deleted. It also assists in keeping magnetosomes to the positively curved membranes when *mamY* is deleted, indicating that McaA contributes to multiple aspects of magnetosome chain organisation. Recently, a curvature-inducing protein CcfM has been identified and characterized in MSR-1 ^39^. CcfM localizes in a filamentous pattern along the positively curved inner membrane by its coiled-coil motifs, and it also functions as an intermediate protein that link the interaction between MamY and MamK ^39^. Whether CcfM plays any role in McaA localization and its interactions with MamK still need to be investigated.

Although many MTB species from different taxa contain the McaA and McaB homologs, only AMB-1 and two other species of *Proteobacteria* form a fragmented magnetosome chain. Given our data, the evolutionary history of the fragmented chain formation strategy in AMB-1 may be explained by several scenarios. In the first one, the MIS could have been acquired after AMB-1 emergence from an unknown magnetotactic *Rhodospirillaceae* donor through a bacteriophage-mediated lateral gene transfer. Indeed, the MIS is flanked with putative phage-related proteins that appears to be very well conserved in *Rhodospirillaceae* spp. Even if this scenario is parsimonious because it minimizes the number of evolutionary events, the presence of homologs of several magnetosome genes in the MIS with partial synteny conservation and many transposases could also be evidence for an old duplication event. In this case, the MIS would be a partial remnant of one of the duplicated versions. The duplication event at the origin would be even more ancestral to the one that led to the emergence of the *lim* cluster ^35^. However, given the known *Magnetospirillum* evolutionary history ^35^, this scenario would imply that many independent losses occurred over *Magnetospirillum* diversification. Assuming the latter scenario, fragmented chain formation would either be the trace of an ancestral strategy progressively replaced in the majority of lineages, or a recent one that emerged in AMB-1.

Our discoveries also highlight that previously undiscovered genes (*mca* and *mca-like* homologs) outside of MAI and conserved in diverse MTB species can play essential roles in magnetosome biosynthesis. The function of Mca-like proteins conserved in MTB remains to be elucidated; their proximity to the MAI, the conservation of their synteny and the presence of the VWA domain in McaA-like proteins indicate that they probably play important role in magnetosome positioning along the magnetoskeleton.

It is notable that the action of two proteins is sufficient to fundamentally alter the assembly and organisation of magnetosome chains in AMB-1 as compared to MSR-1, one of its closest relatives. We propose that the alternative mode of chain organisation in AMB-1 may provide advantages that have led to its selective maintenance. Since magnetic particles are arranged as sub-chains along the length of AMB-1, daughter cells are ensured to inherit equal numbers of magnetic particles that are centrally positioned. Additionally, the distribution and spacing of CMs and EMs may reduce the forces needed to separate magnetic particles. In contrast, MTB such as MSR-1, need to break the closely located continuous crystal chain in the middle and dynamically reposition the entire chain after cell division, which could be more energy-demanding than the stationary ones in WT AMB-1. However, as seen in our Cmag data, AMB-1 cells, as a population, align better in magnetic fields in the absence of *mcaAB*. Given the specific biological interventions required for their assembly, preserved magnetite or greigite chains are also considered an important criterion for magnetofossil recognition and characterization ^10, 11^. Thus, understanding the selective pressures that dictate the species-specific mechanisms of chain organisation in modern day organisms can provide much needed insights into the conditional functions of magnetosomes across evolutionary time.

## Methods

### Bacterial growth

The 1.5-mL stock cultures of AMB-1 strains were prepared as described previously ^40^. The stock cultures were then used for larger volume growth with a dilution of 1:100. For Cmag measurements, TEM, cryo-ET, and fluorescent microscopy, 100 μL of stock cultures were added into 10 mL of MG medium in the 24-mL green-capped tubes and kept in a microaerobic glovebox (10 % oxygen) at 30 °C for 1 to 2 days. For anaerobic growth conditions, 100 μL of stock cultures were added into 10 mL of MG medium in sealed Balch tubes. The MG medium was flushed with N2 gas for 10 min and then was autoclaved. For both microaerobic and anaerobic growth conditions, 1/100 vol of Wolfe’s vitamin solution and 30 μM ferric malate were added to the MG medium just before inoculation with bacteria. For low iron growth condition, 100 μL of stock cultures were added in the 10-mL MG medium supplied with only 1/100 vol of Wolfe’s vitamin solution, but no ferric malate, in the green-capped tubes that treated with 0.375 % oxalic acid (to remove the trace iron on the wall of glass tubes).

*E. coli* strains DH5a, XL1 Blue, DHM1, and WM3064 were grown in Lysogeny broth (LB) medium with appropriate antibiotics. For *E. coli* strain WM3064, 300-µM diaminopimelic acid was added to the LB medium before inoculation with bacteria.

### Genetic manipulation

The genome sequence of *Magnetospirillum magneticum* AMB-1 (GenBank accession number NC_007626.1) was used for oligonucleotide design. Oligonucleotides were purchased from Elim Biopharm or Integrated DNA technologies. All constructs were confirmed by sequencing in UC Berkeley DNA Sequencing Facility. Plasmids were constructed by PCR amplifying DNA fragments of interest with the Phusion High Fidelity DNA Polymerase (New England Biolabs) or CloneAmp HiFi PCR Premix (Takara). All plasmids were introduced into AMB-1 by conjugation. The details about generation of plasmids and strains are described in the supplementary methods. The strains, plasmids, and primers used in this study are described in Supplementary Tables 5 to 12.

### Deletion mutagenesis

A two-step homologous recombination method was used to generate deletion mutants in AMB-1 strains as previously described ^41^. Briefly, an approximately 800 to 1000 bp region upstream and downstream of the deleted gene or genomic region were PCR amplified from the AMB-1 genomic DNA using primer pairs (A, B) and (C, D), respectively (Supplementary Table 8). The two PCR fragments were cloned into the SpeI restriction site of the pAK31 suicide plasmid using Gibson assembly to generate the deletion plasmids (Supplementary Table 6). The deletion plasmid was conjugated into AMB-1 strain using *E.coli* WM3064 donor strain. Colonies that had successfully integrated the plasmid were selected on MG agar plates containing 15 μg/mL kanamycin. To select for colonies that had undergone a second recombination event to lose the integrated plasmid, a counter-selectable marker sacB, which is toxic in the presence of sucrose, was used. Colonies were then passed in 10 mL of growth media without kanamycin and plated on MG agar plates containing 2% sucrose. The resulting sucrose-resistant colonies were checked for the successful deletions at their native locus by colony PCR with primers listed in Supplementary Table 9.

### Cellular magnetic response

The optical density at 400 nm (OD_400_) of AMB-1 cultures in the green-caped tubes was measured at 24 hours and 48 hours using a spectrophotometer. A large magnet bar was placed parallel or perpendicular to the sample holder outside the spectrophotometer, the maximum and minimum OD_400_ were recorded. The ratio of the maximum to the minimum was designated as AMB-1 cells’ Cmag.

### Transmission electron microscopy (TEM)

For imaging the whole AMB-1 cells by TEM, 1-mL AMB-1 cells were taken from the 10-mL cultures that grew under different conditions. The 1-mL cells were pelleted and resuspended into 5-10 µL of MG medium. The resuspended cells were applied on a 400-mesh copper grid coated with Formvar and carbon films (Electron Microscopy Sciences). The grids were glow-discharged just before use. Then the air-dried cells were imaged on an FEI Tecnai 12 transmission electron microscope equipped with a 2k x 2k charge-coupled device (CCD) camera (The Model 994 UltraScan®1000XP) at an accelerating voltage of 120 kV. Crystal size quantification and statistical tests were performed as described previously ^42^.

### Cryo-electron tomography (cryo-ET)

Cryo-ET sample preparation and data collection were performed as described previously ^42^. The two-dimensional (2D) images of WT and ΔMIS cells were recorded using JEOL JEM–3100 FFC FEG TEM (JEOL Ltd.) equipped with a field emission gun electron source operating at 300 kV, an Omega energy filter (JEOL), and a K2 Summit counting electron detector camera (Gatan). Single-axis tilt series were collected using SerialEM software ^43^ from −60° to +60° with 1.5° increments, at a final magnification of 6,000x corresponding to a pixel size of 0.56 nm at the specimen, and a defocus set to −15 μm under low dose conditions (a cumulative electron dose of ∼120 e/A^2^). Tomogram reconstructions were visualized using the IMOD software package ^44^. Amira was used for the 3D model segmentation (Thermo Fisher Scientific).

### Size and location analysis of magnetosome membranes

The reconstructed tomograms were visualized using a 3dmod software package ^44^. To evaluate the relative size and location of magnetosome membranes, the diameter of each magnetosome membrane was measured as described previously ^40^. The size of magnetosome membranes in each cell was sorted from largest to smallest and then was nominated from 1 (largest) to 0 (smallest) accordingly. The first magnetosome membrane on the left of the chain was numbered as 0, the middle one was numbered as 1, and the last magnetosome membrane on the right of the chain was numbered as 2. The location of other magnetosome membranes was nominated accordingly.

For measuring the distance between individual magnetosome membranes, the shortest distances between neighbouring magnetosome membranes were found by working through the tomogram slices and manually measured by 3dmod.

### Structured illumination fluorescent microscopy (SIM)

To stain the genomic DNA, AMB-1 cells growing in 10-mL MG medium of the green-capped tubes were collected by 16,800 x g for 3 min. The cell pellets were resuspended in a 1-mL fresh MG medium and stained with 1.4 µM 4′,6-Diamidino-2-Phenylindole (DAPI) in dark at room temperature for 15 min, and then washed 3 times with fresh MG medium. After washing, the pellet cells were resuspended with 30-50 µL of MG medium and were immediately imaged by Carl Zeiss Elyra PS.1 structured illumination microscopy with objective lens Plan-APOCHROMAT 100 ×/1.46. DAPI, GFP, JF549, and JF646 were excited by 405 nm, 488 nm, 561 nm, and 642 nm lasers, respectively, and fluorescence from each fluorophore was acquired through 420-480 nm, 495-550 nm, 570-620 nm, and LP655 nm bandpass filters, respectively. Raw images were acquired and processed using ZEN software (Zeiss). The processed images were then visualized using Imaris (Bitplane).

MamI-GFP localization patterns in WT and ΔMIS cells were manually measured using the Fiji software package ^45^. The magnetosome chain showed a linear line across the cellular axis, so the end-to-end distance of the GFP fluorescence line was considered as the length of the magnetosome chain. A line that parallels the magnetosome chain was drawn from cell pole to pole, and this line was considered as the length of the whole AMB-1 cell.

### Pulse-Chase analysis

To study the addition of newly-made empty magnetosomes into magnetosome chains over time, we applied pulse-chase analysis using Halo-tagged magnetosome proteins as magnetosome markers. Halo ligands can irreversibly bind to Halo proteins. Under standard growth conditions, WT or mutated AMB-1 cells expressing Halo-tagged magnetosome proteins were grown to early exponential phase (OD_400_ is ∼0.05). Cells were pelleted and resuspended with 60-μL MG medium and mixed with 120-µL 5 µM pulse Halo ligand JF549, kept at 30 °C for 2.5 hours in dark to make sure the Halo proteins were stained saturated with JF549. Then the extra JF549 ligand was washed away with MG medium (3 times, 10 min for each time). keep a few microliters of JF549-stained cells for SIM microscopy imaging. Put the rest cells back in a green-capped tube containing 10-mL MG medium, Kept in the 10% oxygen glove box at 30 °C for 4 hours in dark to allow new protein and new magnetosome production. Cells were then pelleted and resuspended with 60-μL MG medium, and mixed with 120-µL 5 µM chase Halo ligand JF646, kept at 30 °C for 1 hour in dark. Then the extra JF646 ligand was washed away with MG medium (3 times, 10 min for each time) and imaged immediately with a SIM microscope. For the control group, after one quick wash with MG medium, pulse-stained cells were fixed with 4% paraformaldehyde for 1 hour at room temperature to prevent the production of new Halo-tagged proteins, then washed with MG medium three times and kept for the chase staining. For transition experiments from iron starvation to standard iron growth conditions, AMB-1 cells were first grown with an MG medium without added iron and stained with JF549, then were inoculated to the MG medium with iron and incubated in the 10% oxygen glove box for 4-5 hours before JF646 staining.

### Quantitative colocalization analysis of fluorescent-labelled magnetosome proteins

After image collection from the SIM microscope, the subcellular distributions of GFP-and Halo-labelled proteins or different-coloured ligands during pulse-chase experiments were quantitatively analysed by Pearson’s Correlation Coefficient (PCC) and Manders’ Colocalization Coefficients (MCC) ^46^ using the ImageJ JaCoP plugin (Supplementary Table 1). To avoid background and noise signals, only the region of interest (magnetosome chain) was cropped from the original image and measured for colocalization analysis.

### Fluorescence recovery after photobleaching (FRAP)

10 mL of AMB-1 cells in the early exponential phase (OD_400_ is ∼0.05) were pelleted and resuspended in ∼20 µL of MG medium. 3 µL of concentrated AMB-1 cells were applied in a glass-bottom dish (MatTek Corporation) and then covered with 2% solidified agarose for cell immobilization. The 2% agarose covers were formed by spotting 400-µL melted 2% agarose prepared in MG medium to the hole of the glass-bottom dish and solidifying for more than 30 min.

FRAP experiments were carried out on an inverted Carl Zeiss LSM880 FCS laser scanning confocal microscope with an objective lens Plan-Apochromat 100x/1.40 Oil DIC. MamK-GFP filaments were imaged using 488 nm excitation at 0.4 % laser power and fluorescence was acquired through a 490-600 nm bandpass filter. A small area of the filaments was bleached using 488 nm laser light at 100 % laser power for 5 iterations with intervals of 30 s and 30 cycles. For each strain, images were captured through the LSM880 Zen software (Zeiss) and analysed using Fiji ^45^.

### Time-lapse imaging using HILO microscopy

For sample preparation, round coverslips (Matsunami, 25-mm diameter, 0.12–0.17 mm thick) were used as the imaging support. The coverslip was coated with poly-L-lysine and 500 µL of culture was added to an Attofluor cell chamber (Thermo Fisher Scientific). Then, a 5-mm thick gellan gum pad (containing 0.55% gellan gum and 0.08 mM MgCl_2_ in MG liquid medium) on the top of the coverslip to sandwich the cells against the bottom coverslip during time-lapse imaging. After removing excess culture by a pipette, the chamber was filled with fresh MG liquid medium, and the top of the chamber was covered with another coverslip to allow adequate microaerobic conditions to support the growth of AMB-1 cells. The sample was set up under about 10% oxygen atmosphere. Bacteria imaging and processing were then performed as previously described ^16^.

### Protein secondary structure prediction Methods

Membrane topology prediction method CCTOP (ref or http://cctop.enzim.ttk.mta.hu) was used with the TM Filter and Signal Prediction. The CCTOP prediction is a consensus of 10 different methods enhanced with available structural and experimental information of any homologous proteins in the TOPDB database. Signalp 4.1 (http://www.cbs.dtu.dk/services/SignalP-4.1/) and Phobius (https://phobius.sbc.su.se) were used for signal peptide prediction. SMART (http://smart.embl-heidelberg.de) and InterProScan (https://www.ebi.ac.uk/interpro/search/sequence/) were used for protein domain prediction.

### Cellular fractionation

WT AMB-1 cells expressing pAK1255 were first grown in 50-mL MG medium in conical tubes with a 1:100 dilution from stock cultures at 30°C for two days and then grown in 2-L MG medium in a microaerobic glove box (10% oxygen) for two days. These 2-L cells were pelleted by centrifugation at 8,000 × g for 15 min and kept at −80 °C freezer for future use. Cell pellets were thawed on ice and resuspended in 5-mL ice-cold 25 mM Tris buffer (pH 7.0). Pepstatin A and leupeptin were added to a final concentration of 1 µg/mL, and PMSF was added to a final concentration of 1 mM. The resuspension was passed through a French press two times at 1000 psi. From this step, all samples were kept on ice or at 4 °C. 20 µg/mL DNase I and 2 mM MgCl_2_ were added to the homogenate and incubated at 4 °C for 30 min. To separate the magnetosome fraction, the cell lysates were passed through a magnetized MACS LS column (Miltenyi Biotec Inc.) that was surrounded by magnets. After washing the column 3 times with 25 mM Tris buffer (pH 7.0), the magnets were removed and the magnetosome fraction was eluted in 5-mL of 25 mM Tris buffer (pH 7.0). To separate soluble and insoluble non-magnetic fractions, the column flow-through was centrifuged at 160,000 x g for 2 hours. The sedimented membrane fraction was resuspended with 100 mM Tris buffer (pH 7.0) and both fractions were centrifuged a second time at 160,000 x g for 2 hours. The resulting supernatant contained the non-magnetic soluble fraction and the resuspended pellet contained the non-magnetic insoluble fraction.

Cellular fractions were analysed by SDS-PAGE as described previously ^42^. In brief, different fractions were mixed with 2x Laemmli Sample Buffer (Bio-Rad) and heated for 15 min at 95°C. Proteins were resolved by Bio-rad stain-free any KDs gels before transfer to nitrocellulose membrane by electroblotting. Immunological detection was performed with primary antibodies, including anti-GFP polyclonal antibodies (Abcam), anti-Mms6 polyclonal antibodies (Produced by ProSci Inc), or anti-HaloTag monoclonal antibody (Promega), and HRP-Conjugated secondary antibodies (Bio-Rad).

### Bacterial adenylate cyclase two-hybrid assay (BACTH)

The assay was performed as described in the Euromedex BACTH system kit manual. N- and C-terminal T18 and T25 fusions of McdA, McdB, or MamY proteins were constructed using plasmid pKT25, pKNT25, pUT18C, and pUT18 in *E. coli* K12 recA strain XL1-Blue. T18 and T25 fusions with MamK or MamJ were generated previously ^47^. Sequence-verified constructs expressing T18/T25 magnetosome protein fusions were co-transformed into competent *E. coli* DHM1 cells (lacking endogenous adenylate cyclase activity) in all pairwise combinations ^48^, then plated on LB agar plates containing 100 µg/mL carbenicillin and 50 µg/mL kanamycin, and incubated at 30°C overnight. Several colonies of T18/T25 cotransformants were isolated and grown in LB liquid medium with 100 µg/mL carbenicillin and 50 µg/mL kanamycin overnight at 30°C with 220 rpm shaking. Overnight cultures were spotted on indicator LB agar plates supplemented with 40 µg/mL X-gal, 100 µg/mL carbenicillin, 25 µg/mL kanamycin, and 0.5 mM IPTG. Plates were incubated 24 - 48 hours at 30°C before imaging. Bacteria expressing interacting hybrid proteins will show blue, while bacteria expressing non-interacting proteins will remain white.

### Comparative genomics and molecular phylogenetics

McaA and McaB homologs were searched in other bacterial genomes available in public databases. Protein sequences were aligned against reference proteins and non-redundant protein sequences of the *refseq_protein* and *nr* NCBI databases respectively in October 2021 using the BLASTP algorithm, a word size of 6 and default scoring parameters. A similar task was performed using public genomic assemblies of MTB annotated with the Microscope platform ^49^. BLAST hits with an expectation value below 5 × 10^-2^ were further analysed. First, pairwise sequence comparisons were performed using BLASTP (BLAST+ version 2.10.0). Sequence clustering was then performed with the Mmseqs2 ^50^ clustering algorithm version 13.45111 to define groups of distant homologs using the default parameters, a sequence identity threshold of 30% and an alignment coverage of 80% for the longer sequence and for the shorter sequence.

A first phylogenetic tree was built to determine the monophyly of the different clusters and evaluate their taxonomic and phenotypic composition. For this task, the total 38 McaA and 31 McaB homologous sequences retrieved were aligned using MAFFT version 7.487 ^51^. Relaxed trimming on the alignments was then performed using BMGE ^52^, selecting the BLOSUM30 substitution matrix, a minimum block size of 2 and removing characters with more than 50% gaps. Maximum likelihood trees were built using IQ-TREE ^53^ version 2.1.3, the substitution model for each protein was selected by ModelFinder ^54^ with the Bayesian Information Criterion. The statistical support of the branches was estimated by a standard nonparametric bootstrapping approach implemented in IQ-TREE applying 500 replicates. All sequences retrieved belong to magnetotactic bacteria from *Proteobacteria* classes and *Nitrospirae*. Non-Alphaproteobacteria members were used to form external groups and infer the ancestry of McaA and McaB genes compared to other clusters.

Guided by these first phylogenies, a second set of trees was built following the same approach using genome sequences of strains for which transmission electron microscopy images of the magnetosome chains are available and for which organisation could be compared with that of *Magnetospirillum magneticum* AMB-1. Relationships between sequences and chain features were then inferred after collecting all metadata including transmission electron microscopy images published previously (Table S1). The synteny analyses were further explored using the tools implemented in the Microscope ^49^ platform.

## Supporting information

Phylogenetic Dataset

## Acknowledgments

We thank members of the Komeili lab for helpful discussions and suggestions. We thank Pranami Goswami for helping generate the plasmid pAK1037, Julia Borden for helping generate the plasmids pAK1101 and pAK1102, Pedro Leão for helping generate the plasmid pAK1121. We thank Elizabeth Montabana and Kenneth H. Downing for assistance with cryo-ET data collection using the cryo-EM facility at Lawrence Berkeley National Laboratory. We also thank Danielle Jorgens, Reena Zalpuri, and Guangwei Min from the UC Berkeley Electron Microscope Laboratory for assistance in electron microscopy data collection. We thank Steven Ruzin and Denise Schichnes from the CNR Biological Imaging Facility at UC Berkeley for their technical support with fluorescent microscopes. The Zeiss Elyra PS.1 Super-Resolution microscope for SIM was supported in part by the National Institutes of Health S10 program under award number 1S10OD018136-01. We thank the INRA MIGALE bioinformatics platform (http://migale.jouy.inra.fr) for providing computational resources. We thank Michelle C. Chang from Department of Chemistry at UC Berkeley for the kind gift of anti-Mms6 antibodies. Finally, We thank Luke D. Lavis from Howard Hughes Medical Institute Janelia Research Campus for providing free JF549 and JF646 Halo ligands before they are commercialized.

## Author Contributions

J.W. conceived, performed, and analysed most of the experiments. A.T conducted the live-cell imaging analysis with HILO microscopy. C.L.M and C.T.L. performed the phylogenetic analysis of McaA and McaB. G.E. designed and generated the small region (iR1-iR4) deletion plasmids, generated the deletion mutants of ΔiR1-ΔiR4 and Δ*mamJ*Δ*limJ*ΔiR1-Δ*mamJ*Δ*limJ*ΔiR4, and collected the TEM images of these mutants. K.P. generated the deletion mutants of ΔMIS and Δ*mamJ*Δ*limJ* ΔMIS, collected the TEM images of these two mutants. M.A. discovered the crystal shape difference between WT and ΔMIS. E.T-C. helped to establish the Halo-based pulse-chase experiment system. A.K. supervised the experimental design, data analysis, and data presentation. J.W., A.T., C.L.M., C.T.L., and A.K. wrote the manuscript, and all co-authors edited the manuscript.

