## Supplemental methods and figures for "McaA and McaB control the dynamic positioning of a bacterial magnetic organelle"

### Contribute equally

#### Supplementary Results

##### Testing magnetosome marker MmsF for pulse-chase experiments

MmsF is a magnetosome membrane protein that has been used as a magnetosome marker [1, 2]. We first checked the localization of GFP-MmsF in WT and ΔMIS strains. Structured illumination fluorescent microscopy (SIM) imaging shows that GFP-MmsF is located as a continuous chain from cell pole to pole in WT AMB-1, indicating MmsF located to both EMs and CMs (Supplementary Fig. 4a). However, GFP-MmsF is only located in the middle of ΔMIS cells, which is consistent with the magnetosome organisation seen in cryo-ET images (Supplementary Fig. 4a). Similar to MamI-GFP (Fig. 2c), the length ratio of GFP-MmsF marked magnetosome chain versus the whole cell in WT is significantly larger than in ΔMIS (Supplementary Fig. 4b). Halo-MmsF fusion proteins were then used to test if MmsF is a suitable marker for the pulse-chase experiments. As a control, we fixed the pulse ligand JF549-stained cells with 4% paraformaldehyde and then stained them with the chase ligand JF646. The result shows that almost no chase signal is detected in the cells (Supplementary Fig. 4c), indicating Halo-tag staining with the pulse ligand JF549 is saturated. Additionally, when we directly fixed AMB-1 cells and then stained them with the Halo ligand JF549, we could detect the signal that was similar to unfixed cells (Supplementary Fig. 4d), indicating the fixed cells do not prevent the distribution and function of Halo ligands. For the pulse-chase experiments under standard growth conditions, the chase signals of Halo-MmsF colocalise with the pulse signals in both WT and ΔMIS cells (Supplementary Fig. 4e, f), indicating the newly-synthesized MmsF proteins were added to the old magnetosomes. Moreover, quantitative analysis shows very high colocalization coefficients of pulse and chase signals in WT and ΔMIS cells (Supplementary Table 1). Hence MmsF is not a good marker for testing the addition of newly-made magnetosomes.

##### Comprehensive genetic dissection of the MIS genes

To identify the key genes that control magnetosome positioning and crystal shape control in the MIS region, we firstly generated large domain (LD) deletions (ΔMIS_LD1 and ΔMIS_LD2, both including *mamJ-like*) (Fig. 3a). The Cmag, crystal chain organisation, and shape factor of ΔMIS_LD1 is similar to ΔMIS, while ΔMIS_LD2 is similar to WT (Fig. 3b, c and Supplementary Fig. 6c, d), indicating that our genes of interest are in LD1, not LD2. We then generated small islet region deletions (ΔiR1, ΔiR2, ΔiR3, and ΔiR4) of LD1 (Fig. 3a). The Cmag, crystal chain organisation, and shape factors of ΔiR1 and ΔiR4 are similar to WT (Fig. 3b, d and Supplementary Fig. 6e, h). Only the Cmag and crystal chain organisation of ΔiR2 are similar to ΔMIS (Fig. 3b, d). Consistently, the iR2 region can complement the Cmag and crystal chain organisation of ΔMIS and ΔiR2 strains (Supplementary Fig. 7a, b), confirming that the different chain organisation phenotypes in ΔMIS are due to loss of the iR2 region. We then quantified the number and size of crystals in ΔiR2, and the results show that the number and length distributions of crystals in ΔiR2 are similar to ΔMIS (Supplementary Fig. 6a, b), but the shape of crystals in ΔiR2 is similar to WT, not ΔMIS (Supplementary Fig. 1c, d and Supplementary Fig. 6f). Interestingly, the shape factor of crystals in ΔiR3 is similar to ΔMIS (Supplementary Fig. 1d and Supplementary Fig. 6g), which might be the reason for a slightly higher Cmag of ΔiR3 compared to WT (Fig. 3b). Thus, we conclude that there are genes in the iR2 region control magnetosome positioning, while one or more genes in iR3 contribute to crystal shape control.

**Secondary structure prediction and topology confirmation of McaA and McaB**

We predicted the secondary structure of McaA and McaB using several prediction programs (Supplementary Table 2). McaA consists of 776 amino acids (aa), and all membrane prediction programs predicted a transmembrane (TM) domain at around aa 370-390 (Fig. 4a and Supplementary table 2). Some programs predicted the N-terminus aa 1-26 as a signal peptide, and many also predicted the aa 7-22 to be a TM domain with the N-terminus facing the cytoplasm (Supplementary Table 2), followed by a conserved von Willebrand factor type A (VWA) domain (aa 29-258) that is predicted facing the periplasm (Fig. 4a). The C-terminus of McaA is predicted to face the cytoplasm (Fig. 4a).

GFP can be used to discriminate between cytoplasmic and periplasmic domains of bacterial inner-membrane proteins [3, 4]. GFP can fluoresce in the cytoplasm, but cannot fold properly and does not fluoresce when translocated to the periplasm via the *sec* pathway [4]. To validate the membrane topology of McaA, we fused GFP to both the N- and C-terminus of McaA (GFP-McaA and McaA-GFP). We confirmed the ability of both fusion proteins to complement the McaA deletion mutant (Supplementary Fig. 7c, d), and then monitored their fluorescence and localization in WT AMB-1 cells using SIM. If the aa 7-22 is a TM domain, both the N-terminus and C-terminus of McaA would be in the cytoplasm, and both GFP fusions would be fluorescent. However, GFP-McaA was not fluorescent when expressed in WT AMB-1, indicating the GFP might be present in the periplasmic space, not enabling its proper folding, which rules out the TM domain prediction of aa 7-22. Conversely, McaA-GFP was fluorecent with a specific pattern when expressed in WT AMB-1 (Fig. 4b), confirming the C-terminus of McaA is facing the cytoplasm.

McaB consists of 219 aa, and it is predicted to contain an N-terminal TM domain around aa 5-27 (Supplementary Fig. 9a). According to predictions, the N-terminus is mostly facing the periplasm while the C-terminus is located in the cytoplasm (Supplementary Fig. 9a and Supplementary table 2).

To validate the membrane topology of McaB, we fused GFP to both the N- and C-terminus of McaB (GFP-McaB and McaB-GFP). GFP-McaB could not be used to test the topology of McaB due to invalidity of the construct. When GFP-McaB was expressed in WT and Δ*mcaB*, no fluorescence was detected in the cells. The $OD_{400}$ of stationary phase WT/vector and Δ*mcaB*/vector cultures are about 0.25-0.3, but the $OD_{400}$ of stationary phase WT/GFP-McaB and Δ*mcaB*/GFP-McaB cultures are about 0.1, indicating GFP-McaB fusion protein inhibits the growth of both WT and Δ*mcaB* cells. Even the Cmag of Δ*mcaB*/GFP-McaB is lower than Δ*mcaB*/vector, TEM image observation showed that GFP-McaB could not complement Δ*mcaB* (Supplementary Fig. 7e, f). Hence GFP-McaB might not be functional, and cannot be used to test the topology of McaB in AMB-1 cells. However, McaB-GFP complements Δ*mcaB* (Supplementary Fig. 7e, f), and is fluorescent in WT and various mutants, indicating the C-terminus of McaB faces the cytoplasm.

**Co-expressing McaA and McaB in AMB-1 cells**

We co-expressed MacB-GFP and McaA-Halo in the same AMB-1 cell under Tac promoter (pAK1255) or under McaAB promoter (pAK1256) (Supplementary Fig. 10a). Overexpression of McaA-Halo and McaB-GFP in the WT background does not affect biomineralization (Supplementary Fig. 10b). Both pAK1255 and pAK1256 can complement ΔiR2 (Supplementary Fig. 10b, c), indicating these two fusion proteins retain their function under both promoters.

**Localization of McaA and McaB in *E.coli***

To further test the localization pattern of McaA and McaB in different bacteria, we checked the localization of McaA-GFP and McaB-GFP in the rod-shaped *Escherichia coli* (*E.coli)* strains. McaA-GFP is located at both poles of the *E.coli* cells (Supplementary Fig. 12a), indicating the localization pattern of McaA-GFP in AMB-1 might be specific to helical-shaped cells. While

McaB-GFP displayed a mesh pattern close to the cytoplasmic membrane (Supplementary Fig. 12b), which is similar to its localization pattern in AMB-1 cells that do not make magnetosomes (Fig. 4d).

***mamJ-like* is not the gene that prevents magnetosome chain collapse in Δ*mamJ*Δ*limJ* strain**

To investigate whether MamJ-like contributes to the different phenotypes of Δ*mamJ* in MSR-1 strain and Δ*mamJ*Δ*limJ* in AMB-1 strain, we created a triple mutant lacking *mamJ* and all *mamJ* homologs. We deleted *mamJ-like* in both WT and Δ*mamJ*Δ*limJ* strains. The Cmag of Δ*mamJ-like* and Δ*mamJ*Δ*limJ*Δ*mamJ-like* are similar to WT (Supplementary Fig. 13a). TEM images show that Δ*mamJ-like* cells have a similar phenotype to WT (Supplementary Fig. 13b). About 73% of Δ*mamJ*Δ*limJ*Δ*mamJ-like* cells look like Δ*mamJ*Δ*limJ*, and about 27% of them contain a small aggregate in the crystal chain that is located from cell pole to pole (Supplementary Fig. 13b, c). These results indicate that MamJ-like is not the protein that prevents magnetosome chain collapse in Δ*mamJ*Δ*limJ* of AMB-1. Surprisingly, when the whole MIS is deleted in the Δ*mamJ*Δ*limJ* strain, the Cmag of the mutant decreases to about 1 (this shows the cells lose nearly all magnetic response), and the magnetosomes collapse to form a large aggregate (Supplementary Fig. 13b, c), indicating MIS genes (not including *mamJ-like*) contribute to chain maintenance in Δ*mamJ*Δ*limJ* strain.

#### Supplementary Figures

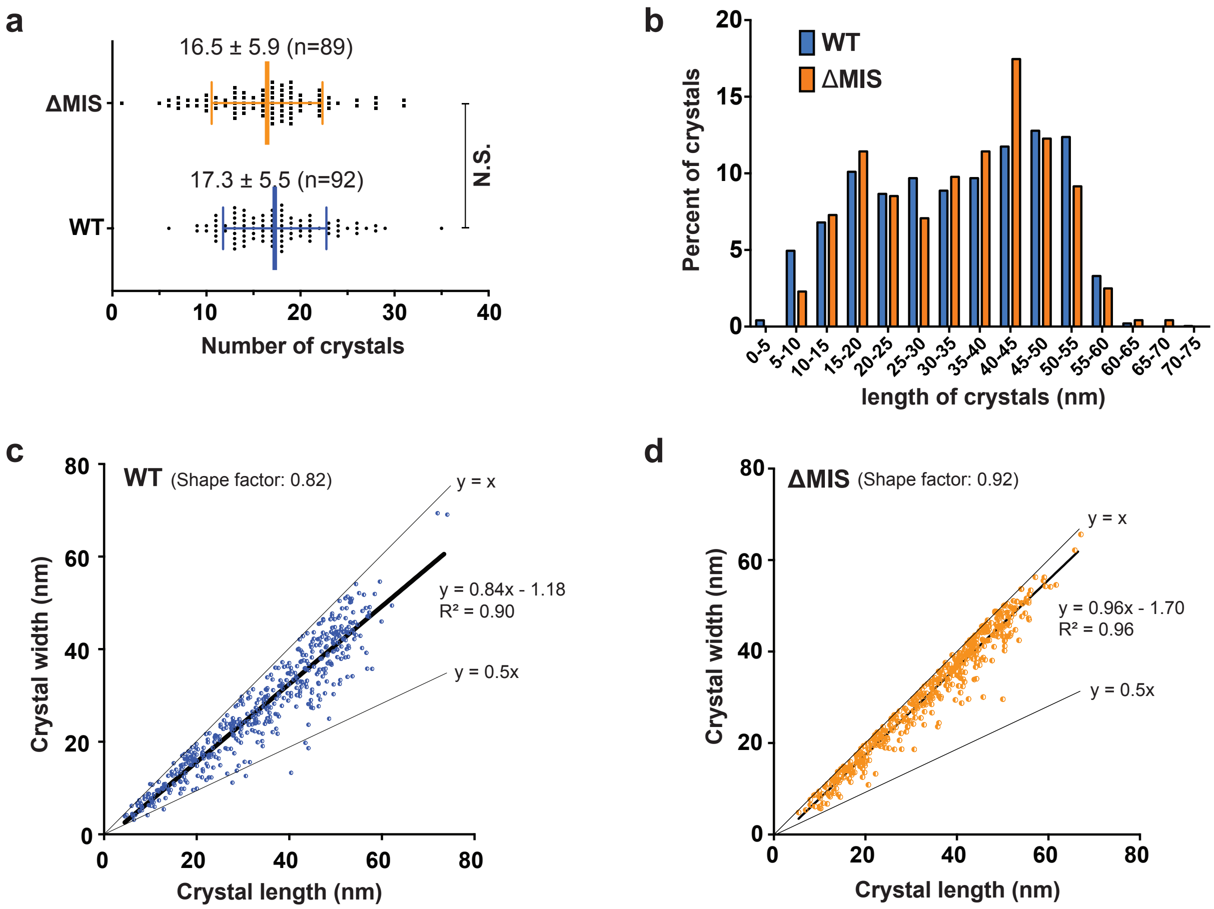


**Supplementary Figure 1: Quantification of the number and size of magnetic crystals in WT and ΔMIS strains that were grown under microaerobic conditions.** Crystal number (a) and length (b) distribution of WT (blue bars) and ΔMIS (orange bars) strains. No statistically significant difference ($P>0.05$, N.S.). Shape factor (width/length ratio) of crystals in WT (c) and ΔMIS (d) strains. The shape factor values in (c) and (d) are the median of the datasets due to their non-normality distribution based on Shapiro-Wilk normality test.

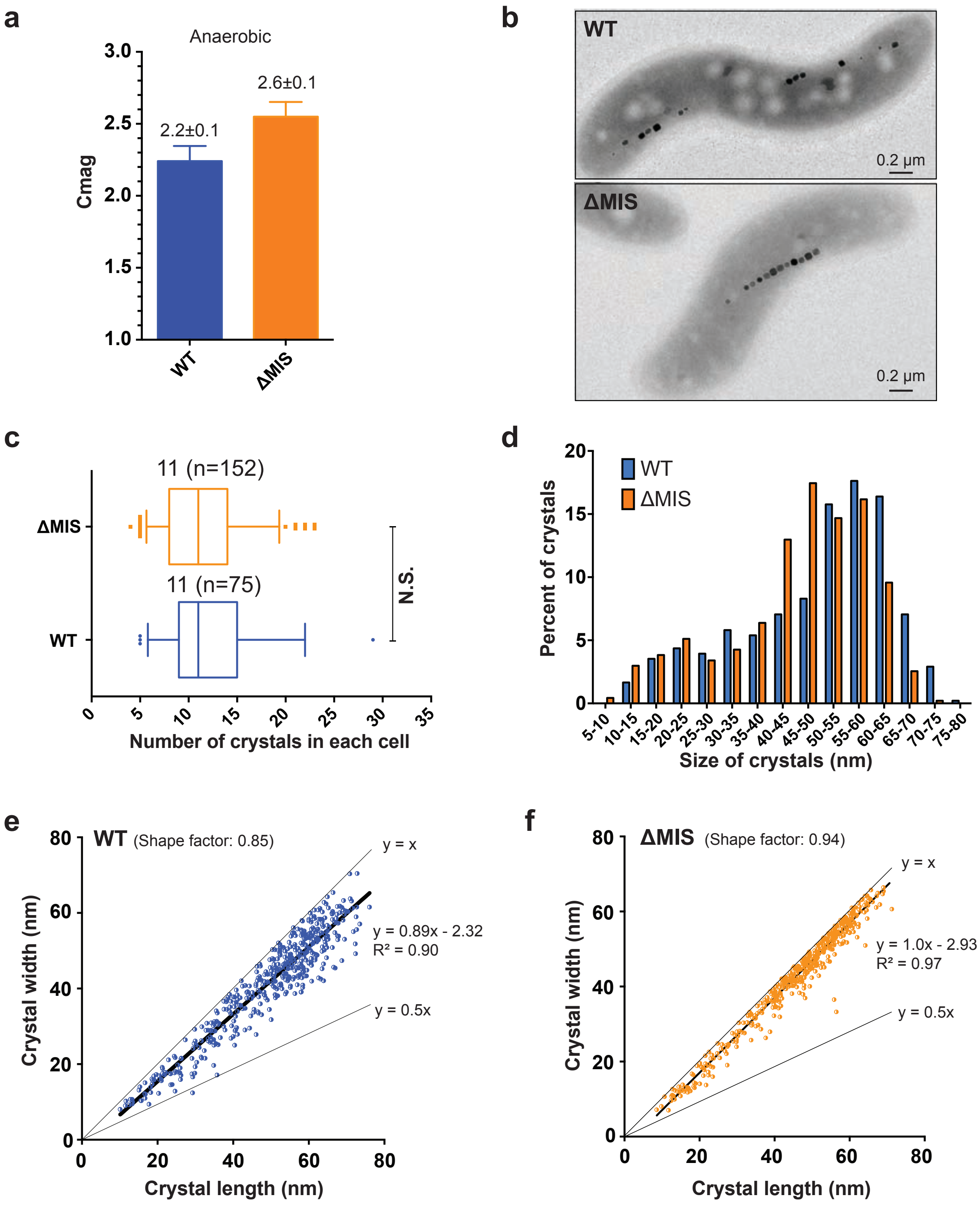


**Supplementary Figure 2: Characterization of the similarity and difference between WT and ΔMIS strains that were grown under anaerobic conditions.** (a) Cmag of WT and ΔMIS cultures. Each measurement represents the average and standard deviation from three independent growth cultures. (b) TEM micrographs of WT and ΔMIS cells. (c) and (d) Crystal number (c) and length (d) distribution of WT (blue bars) and ΔMIS (orange bars) strains. No statistically significant difference ($P>0.05$, N.S.). (e) and (f) Shape factor (width/length ratio) of crystals in WT (e) and ΔMIS (f) strains.

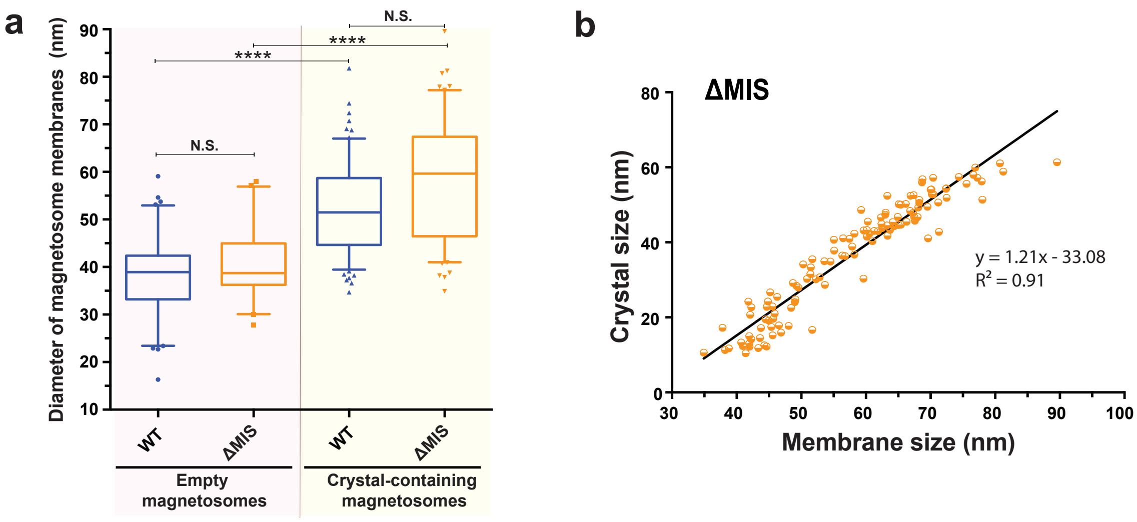


**Supplementary Figure 3: The mechanisms of magnetosome membrane size control are similar in WT and ΔMIS strains.** (a) Magnetosome membrane size distribution in WT and ΔMIS strains. Diameters of empty magnetosome membranes and crystal-containing magnetosome membranes were measured from cryo-electron tomograms of WT and ΔMIS cells. No statistically significant difference (P>0.05, N.S.). Significant difference (****$P < 10^{-4}$). (b) Scatterplot and regression analysis of membrane size versus crystal size for crystal-containing magnetosomes in ΔMIS. The long axis (crystal length) is reported as crystal size.

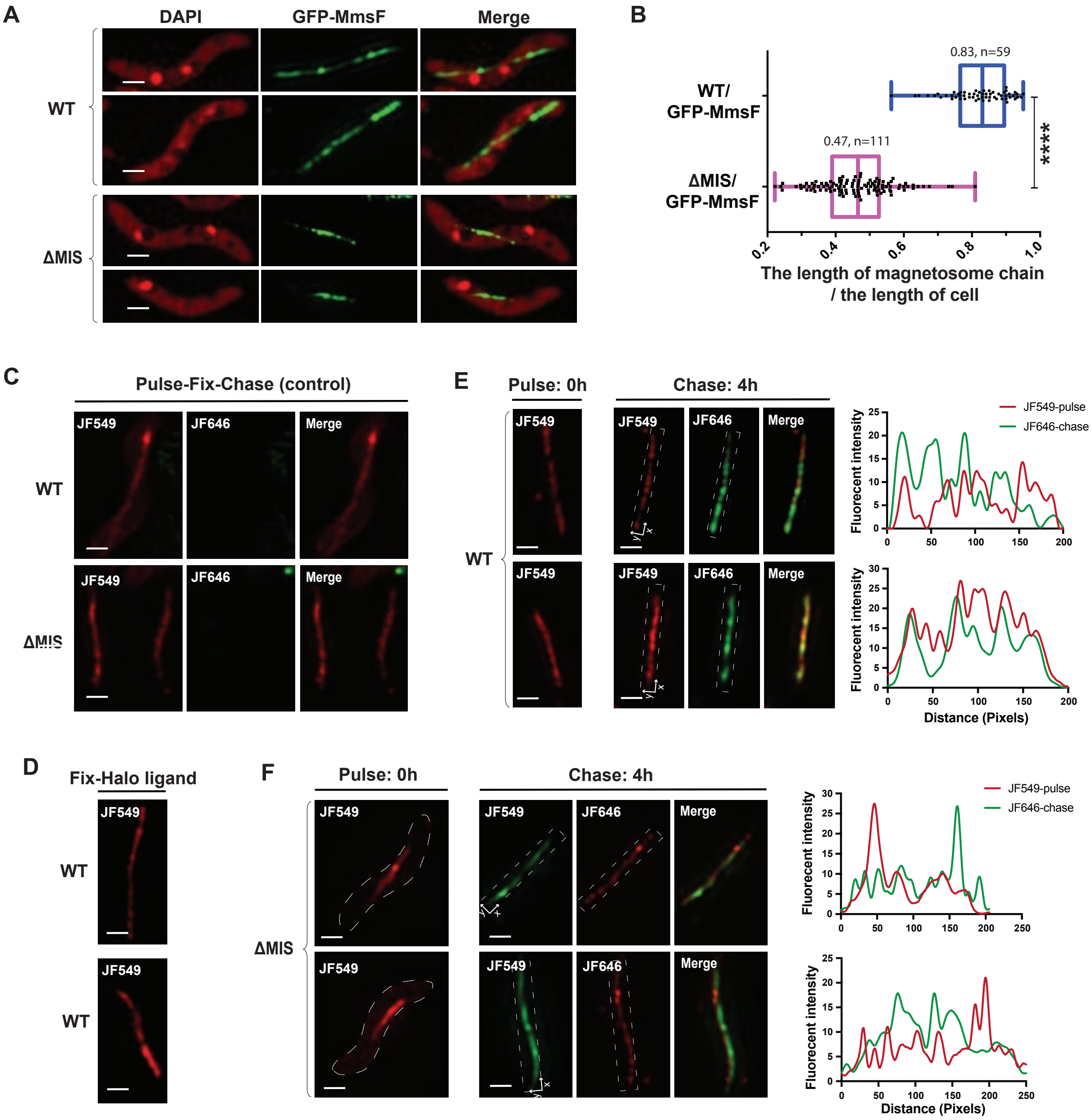


**Supplementary Figure 4: GFP-MmsF is not a good marker for characterizing the addition of newly-formed magnetosomes to the chain.** (a) SIM micrographs of WT and ΔMIS expressing GFP-MmsF under standard growth conditions. The DAPI staining is shown in false-colour red, GFP-MmsF is shown in green. (b) Quantification of the length of magnetosome chain versus the length of cell in WT and ΔMIS strains. (c) Control experiments to make sure that the Halo-tag staining with the pulse ligand JF549 is saturated. (d) Control experiment to test if the Halo ligand could still get into the fixed cells and interact with the Halo proteins. (e) and (f) Left and middle:

SIM micrographs show the pulse-chase experiments with Halo-MmsF fusion protein for analysing the location of newly-formed magnetosomes in WT and ΔMIS stains. The JF549 staining is shown in red, and the JF646 staining is shown in green. Right: fluorescent intensity map of the dashed rectangular area on the SIM micrographs. Scale bars are 0.5 μm in (a) and (c) – (f).

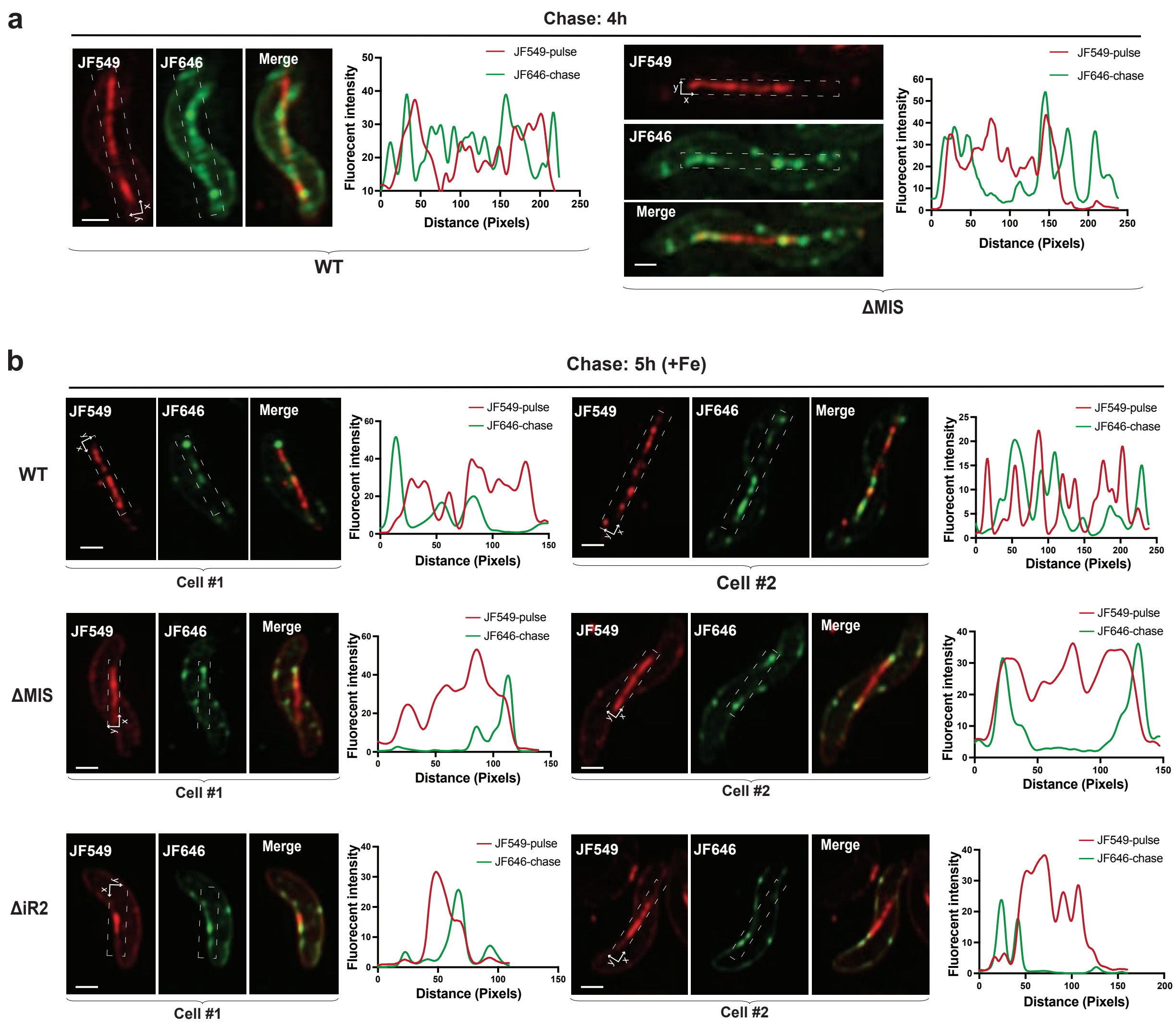


**Supplementary Figure 5: Chase experiments to characterize the addition of newly-formed magnetosomes to the chain.** (a) WT and ΔMIS cells were grown in iron rich conditions. (b) WT, ΔMIS, and ΔiR2 cells were grown from iron starvation to iron rich conditions. For each cell, SIM micrographs (left) and the fluorescent intensity map of the dashed rectangular area (right) show the pulse and chase signals with MamI-Halo fusion proteins. The JF549 staining is shown in red, and the JF646 staining is shown in green. Scale bars: 0.5 µm.

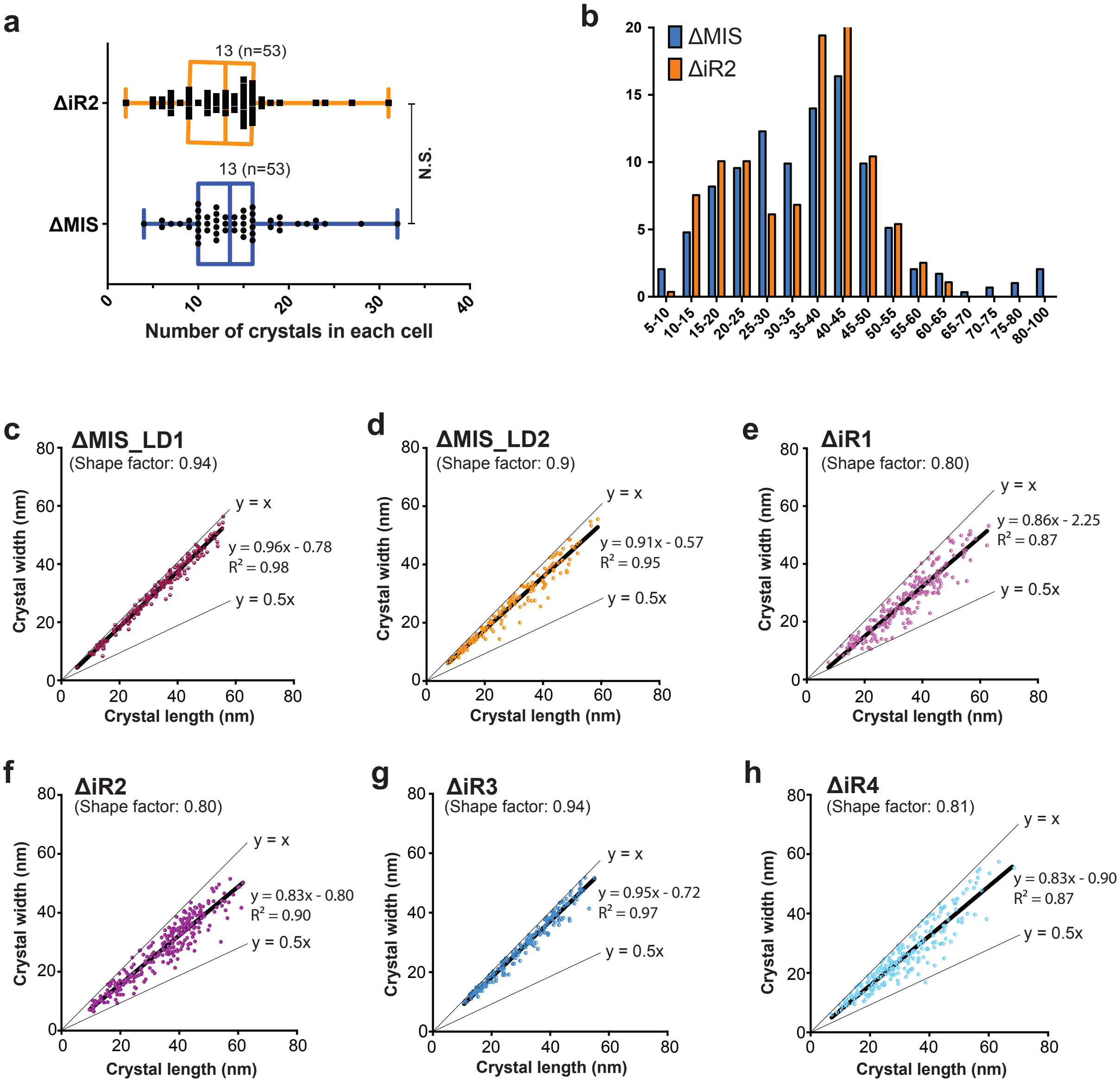


**Supplementary Figure 6: Number and size of magnetic crystals in the islet region deletion mutants.** (a) Number of crystals in ΔiR2 and ΔMIS. No statistically significant difference (P>0.05, N.S.). (b) Crystal length distribution in ΔMIS (blue bars) and ΔiR2 (orange bars) strains. (c) – (h) Shape factor (width/length ratio) of crystals in ΔMIS_LD1, ΔMIS_LD2, and ΔiR1-iR4 strains.

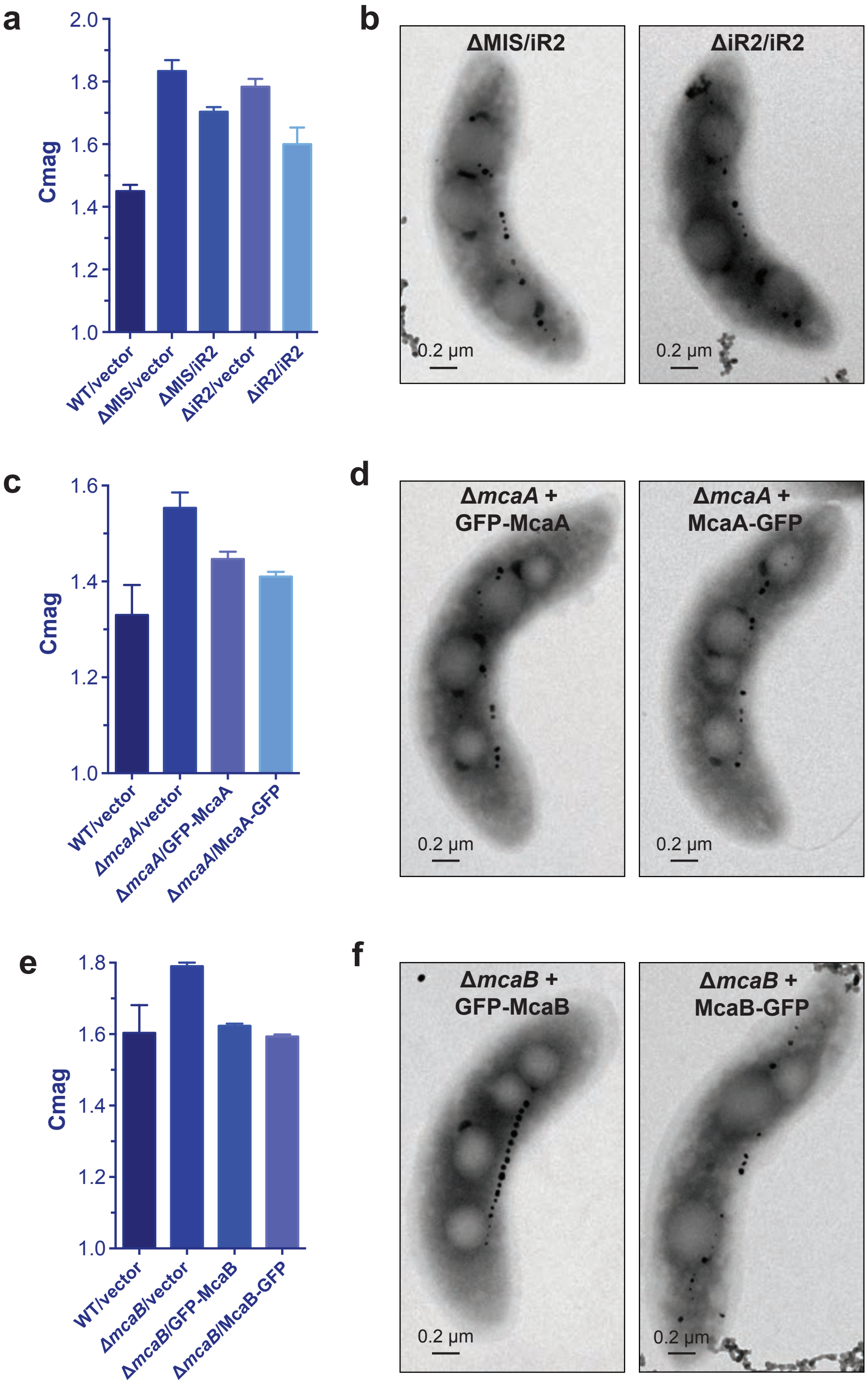


**Supplementary Figure 7: Complementation of ΔMIS or iR2 region deletion mutants.** (a) Magnetic response (Cmag) of ΔMIS and ΔiR2 that complemented with the whole iR2 region. (b) TEM micrographs of ΔMIS/iR2 and ΔiR2/iR2 cells. (c) Cmag of Δ*mcaA* that complemented with GFP-McaA or McaA-GFP. (d) TEM micrographs of Δ*mcaA*/GFP-McaA and Δ*mcaA*/McaA-GFP cells. (e) Cmag of Δ*mcaB* that complemented with GFP-McaB or McaB-GFP. (f) TEM micrographs of Δ*mcaB*/GFP-McaB and Δ*mcaB*/McaB-GFP cells.

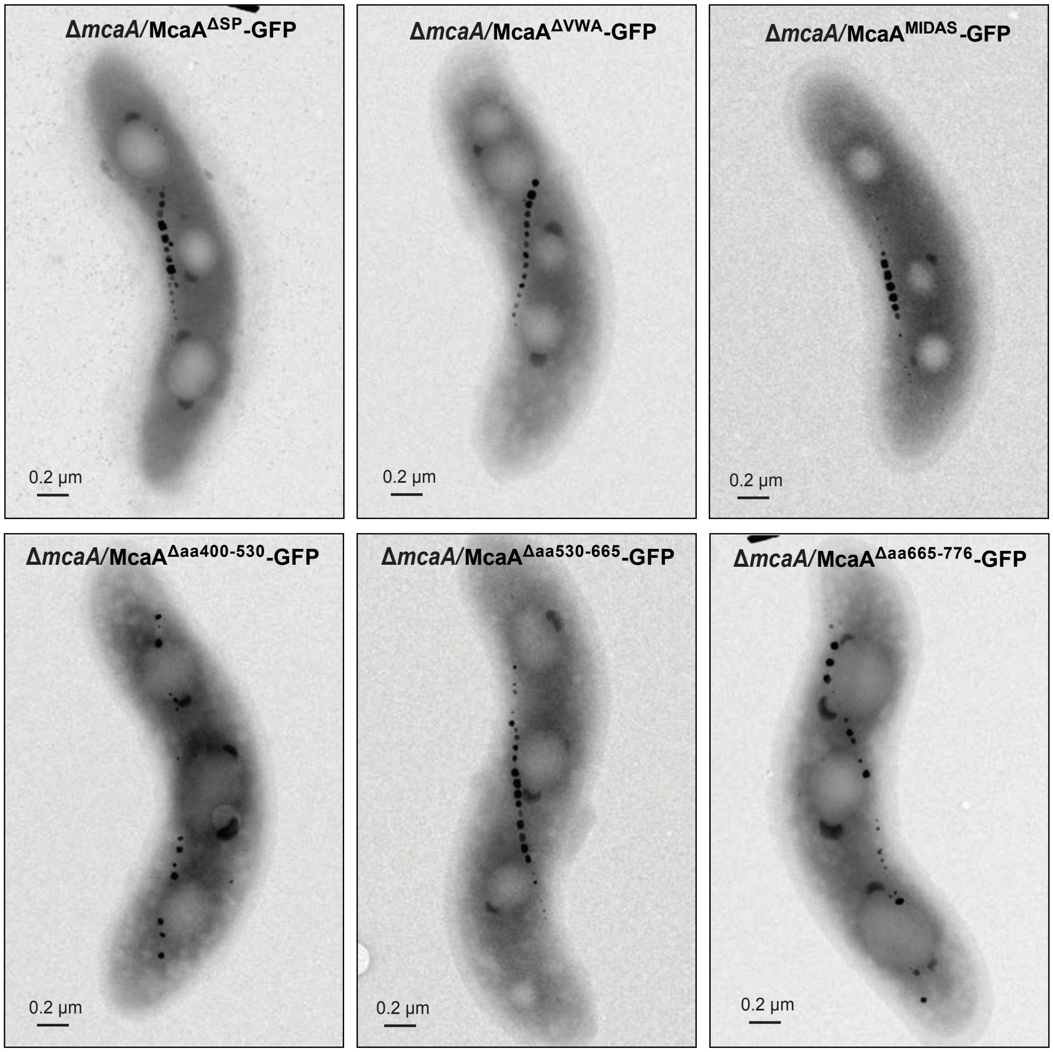


**Supplementary Figure 8: TEM micrographs of different McaA truncation mutants expressed in Δ*mcaA* cells.**

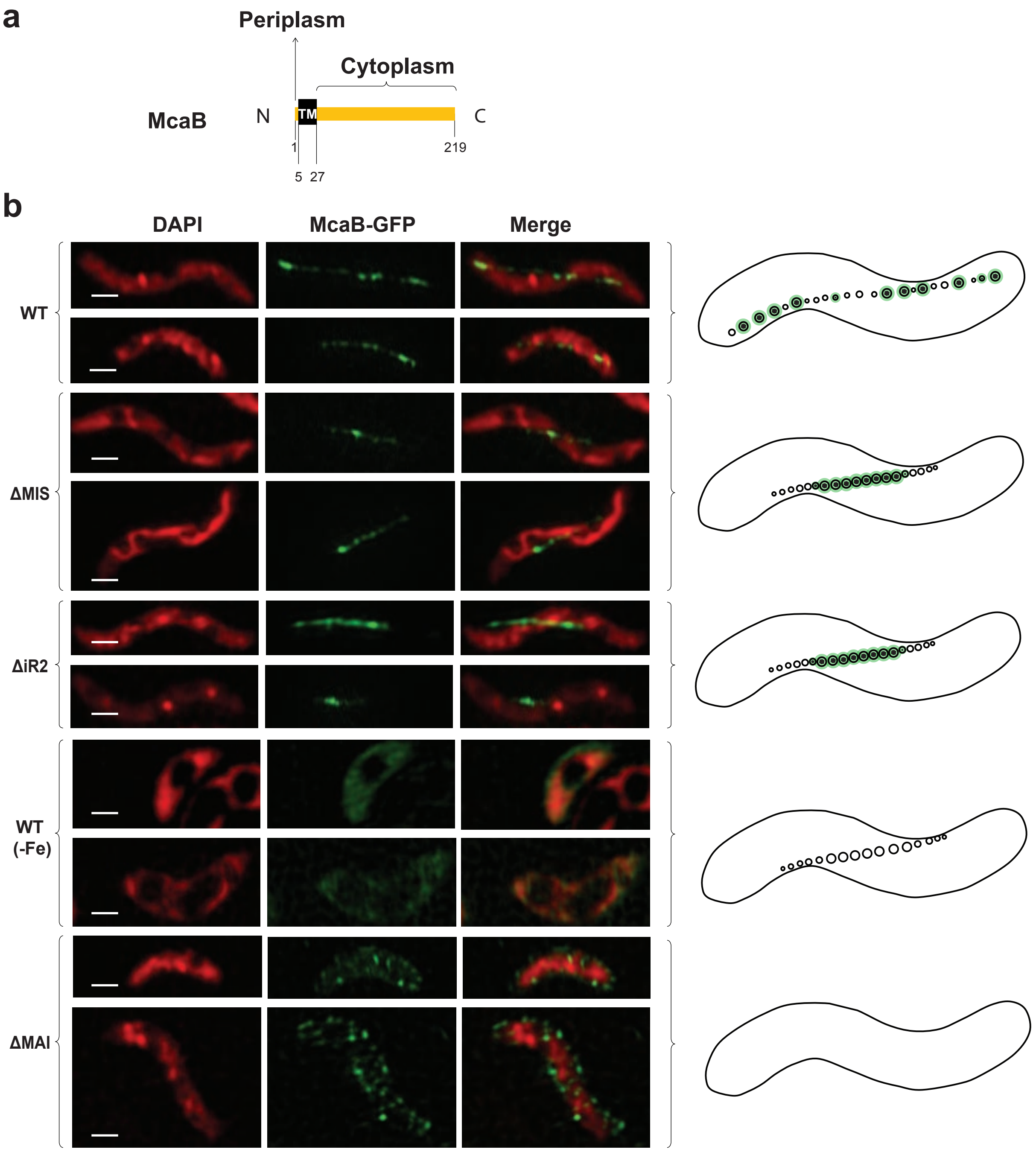


**Supplementary Figure 9: Localization of McaB.** (a)Predicted secondary structure and topology of McaB. TM, transmembrane domain. (b) SIM micrographs show cells expressing McaB-GFP (left) and models of magnetosome production (right) in WT and different genetic backgrounds or growth conditions. The DAPI staining is shown in false-colour red, McaB-GFP is shown in green. Scale bars: 0.5 µm.

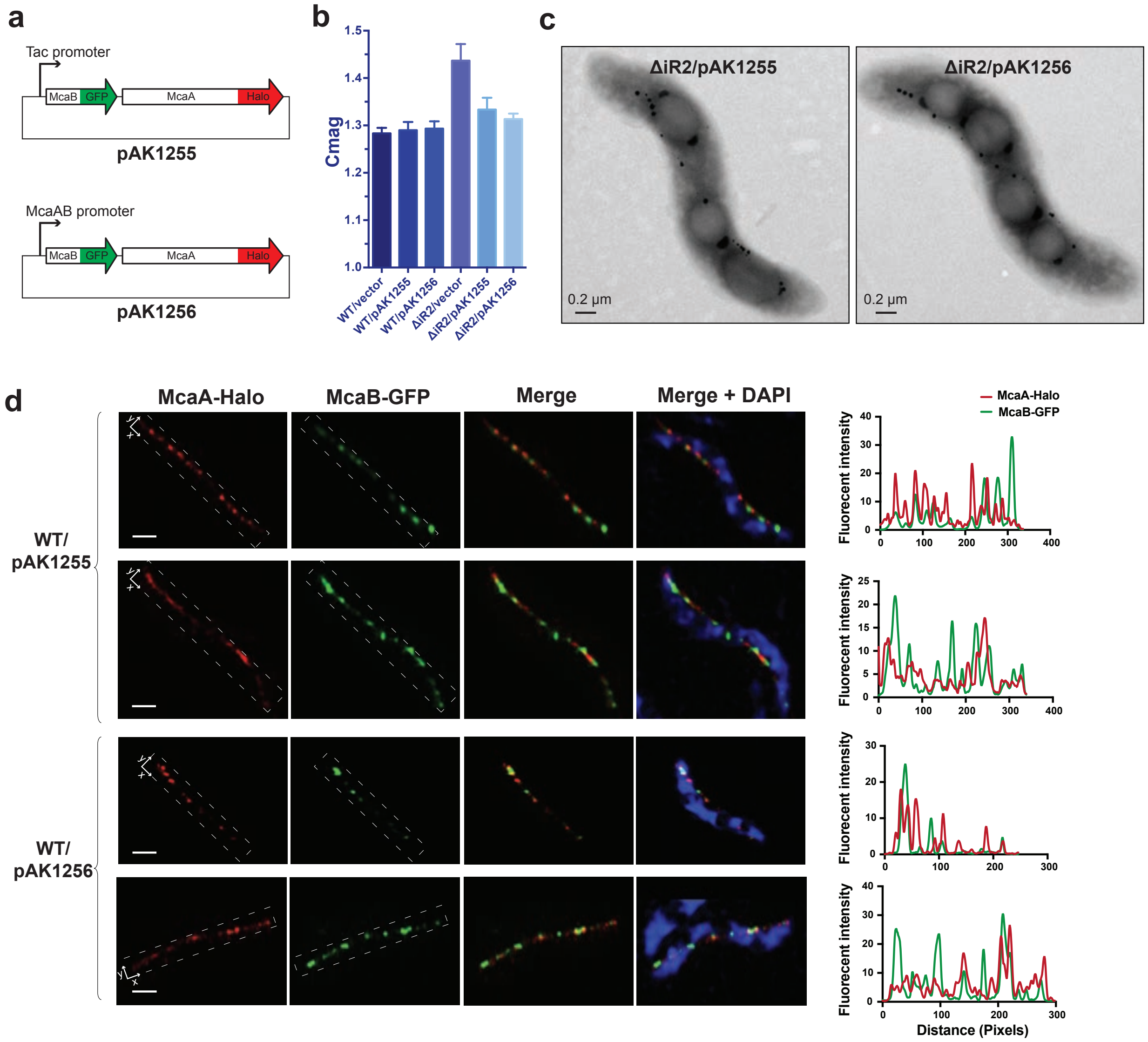


**Supplementary Figure 10: Testing the localization and function of McaA and McaB in the same cell.** (a) Schematic depicting the plasmid constructs of pAK1255 and pAK1256. (b) Cmag of WT and ΔiR2 strains that contains empty vectors, pAK1255 (express McaA and McaB under Tac promoter) or pAK1256 (express McaA and McaB under their native promoter). (c) TEM micrographs of ΔiR2/pAK1255 and ΔiR2/pAK1256. (d) Left: SIM micrographs show the localization of McaA-Halo (stained with Halo ligand JF549) and McaB-GFP in WT AMB-1 cells. The DAPI staining is shown in blue, JF549 staining is shown in red, McaB-GFP is shown in green. Scale bars: 0.5 μm. Right: fluorescent intensity map of the dashed rectangular area on the SIM micrographs.

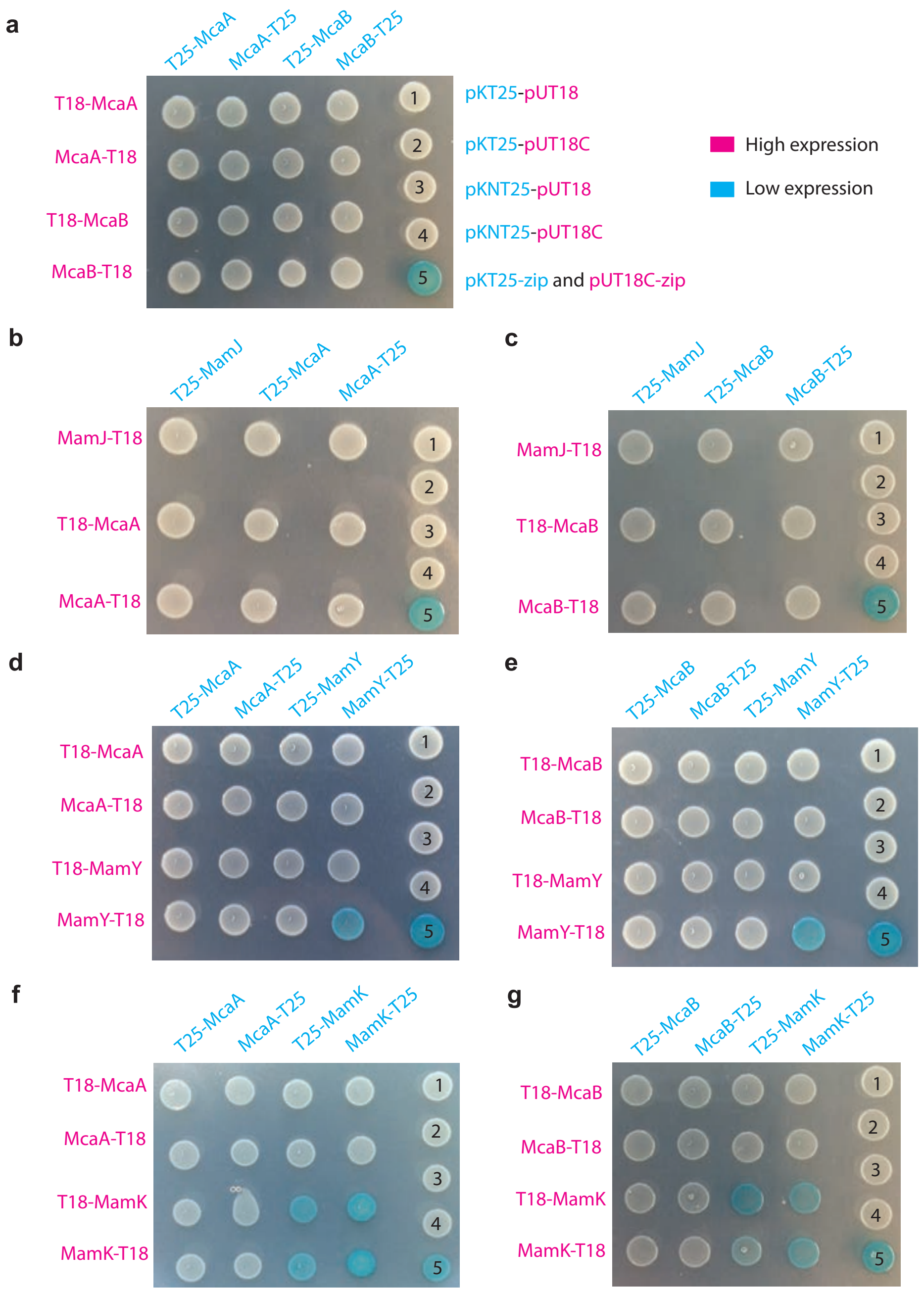


**Supplementary Figure 11: BACTH to identify protein-protein interactions between McaA, -B and MamJ, -Y, -K.** (a) Bacterial two-hybrid between McaA and McaB tagged at their N-termini (X-McaA, X-McaB) or C-termini (McaA-X, McaB-X) on an LB agar plate containing X-gal and IPTG. (b-g) Bacterial two-hybrid between McaA and MamJ (b), McaB and MamJ (c), McaA and MamY (d), McaB and MamY (e), McaA and MamK (f), and McaB and MamK (g). Image representative of 3 independent trials.

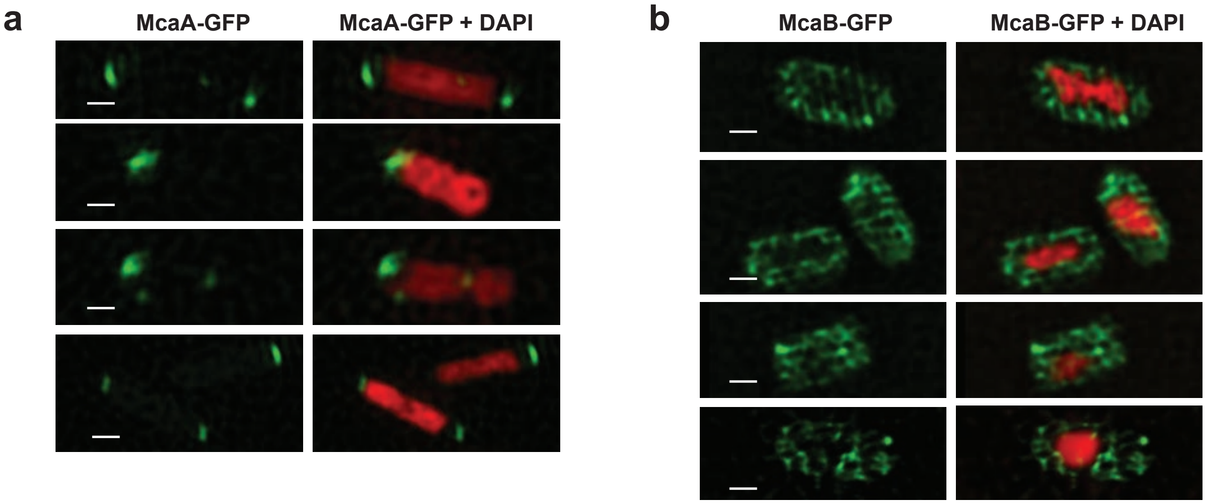


**Supplementary Figure 12: SIM micrographs show *E.coli* cells expressing McaA-GFP (a) and McaB-GFP (b).** The DAPI staining is shown in false-colour red, GFP fusion proteins are shown in green. Scale bars: 0.5 µm.

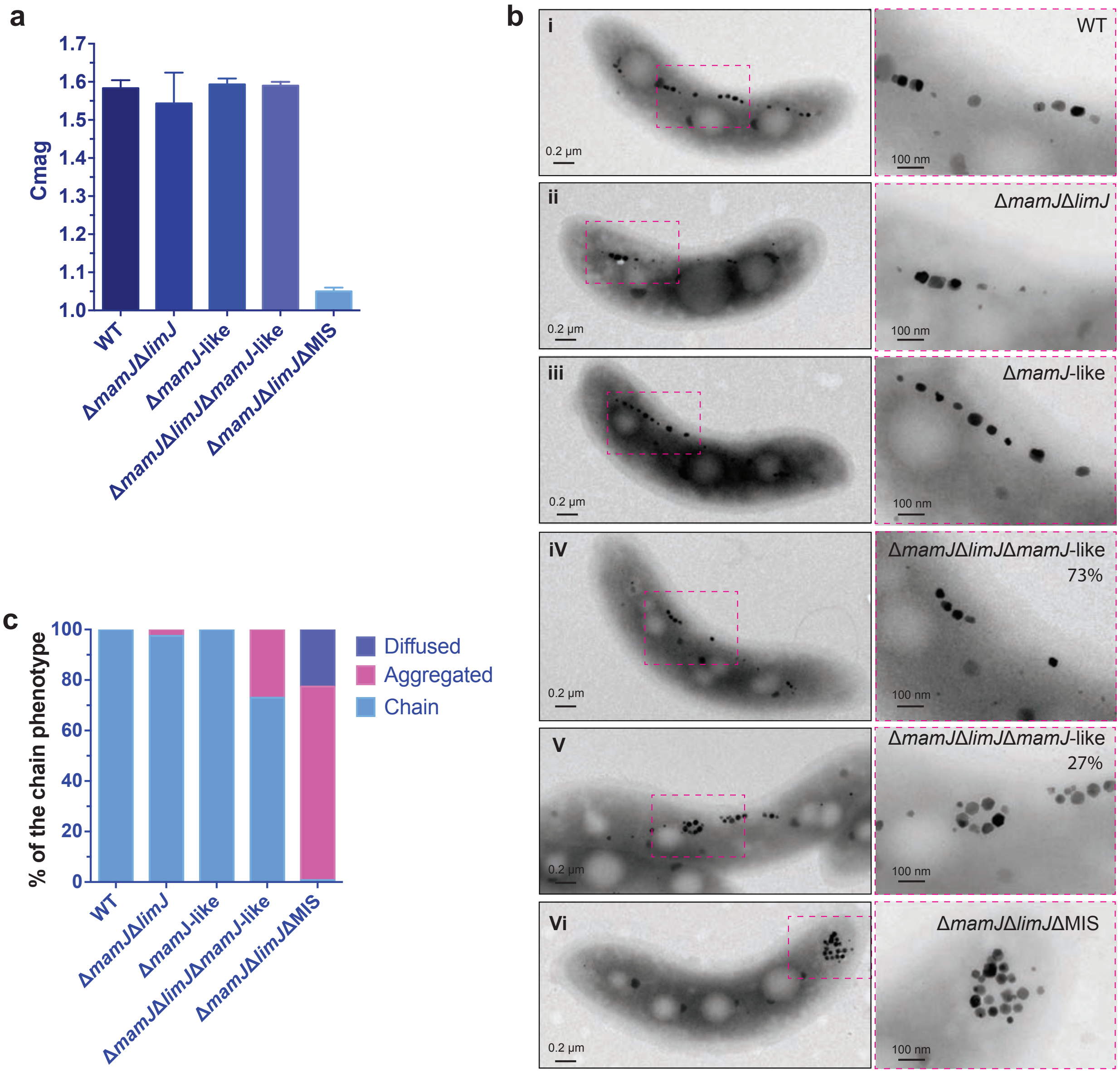


**Supplementary Figure 13: MIS genes (not *mamJ-like*) help to prevent magnetosome aggregation in Δ*mamJ*Δ*limJ* background.** (a) Cmag of WT and *MamJ*-related deletion mutants. (b) TEM micrographs of WT, Δ*mamJ*Δ*limJ*, Δ*mamJ-like*, Δ*mamJ*Δ*limJ*Δ*mamJ-like*, and Δ*mamJ*Δ*limJ*ΔMIS cells. (c) Quantification of the magnetosome chain phenotypes in WT and *MamJ-like* related deletion mutants.

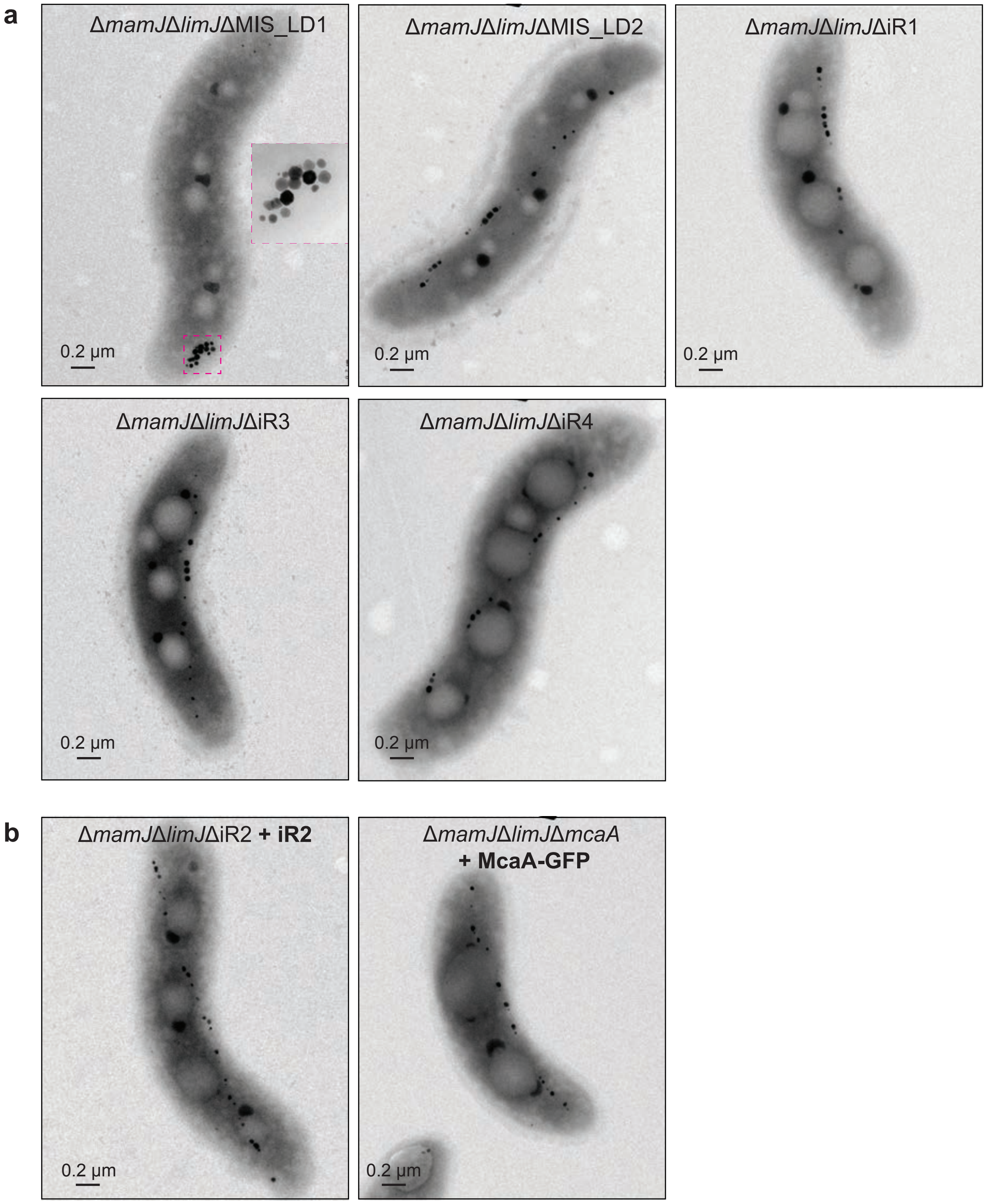


**Supplementary Figure 14: TEM micrographs of MIS large domain and small region deletions in Δ*mamJ*Δ*limJ* backgrounds (a) and the complementation analysis (b).**

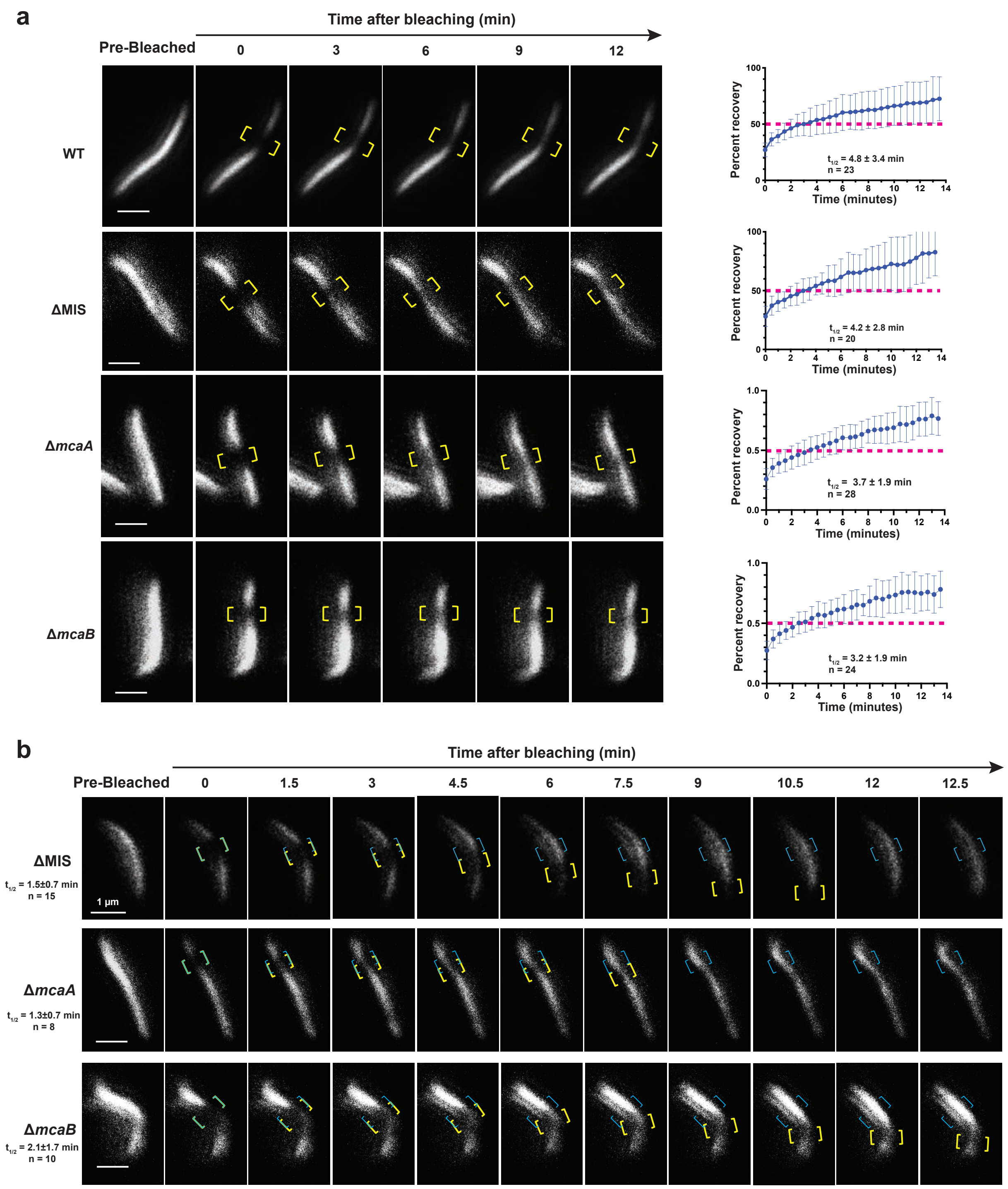


**Supplementary Figure 15: FRAP experiments characterize the dynamics of MamK filaments.** (a) Left: FRAP experiment time courses with WT and different mutants that expressing MamK-GFP. The bleached area is not moving. Yellow brackets indicate the portion of the MamK-GFP filament designated for photobleaching. Right: normalized (average mean and standard deviation

[SD]) percent recovery of each strain's recovering cells with non-moving bleached area. The 50% mark is noted with a dashed magenta line. The MamK-GFP signals are shown in false-colour white. (b) FRAP experiment time courses with ΔMIS, Δ*mcaA*, and Δ*mcaB* cells expressing MamK-GFP where the bleached area moved from its original position toward the cell pole. Yellow and blue brackets indicate the portion of the MamK-GFP filament designated for photobleaching. Blue brackets indicates the original bleaching area, and yellow brackets track the movement of the bleached area. The MamK-GFP signals are shown in false-colour white.

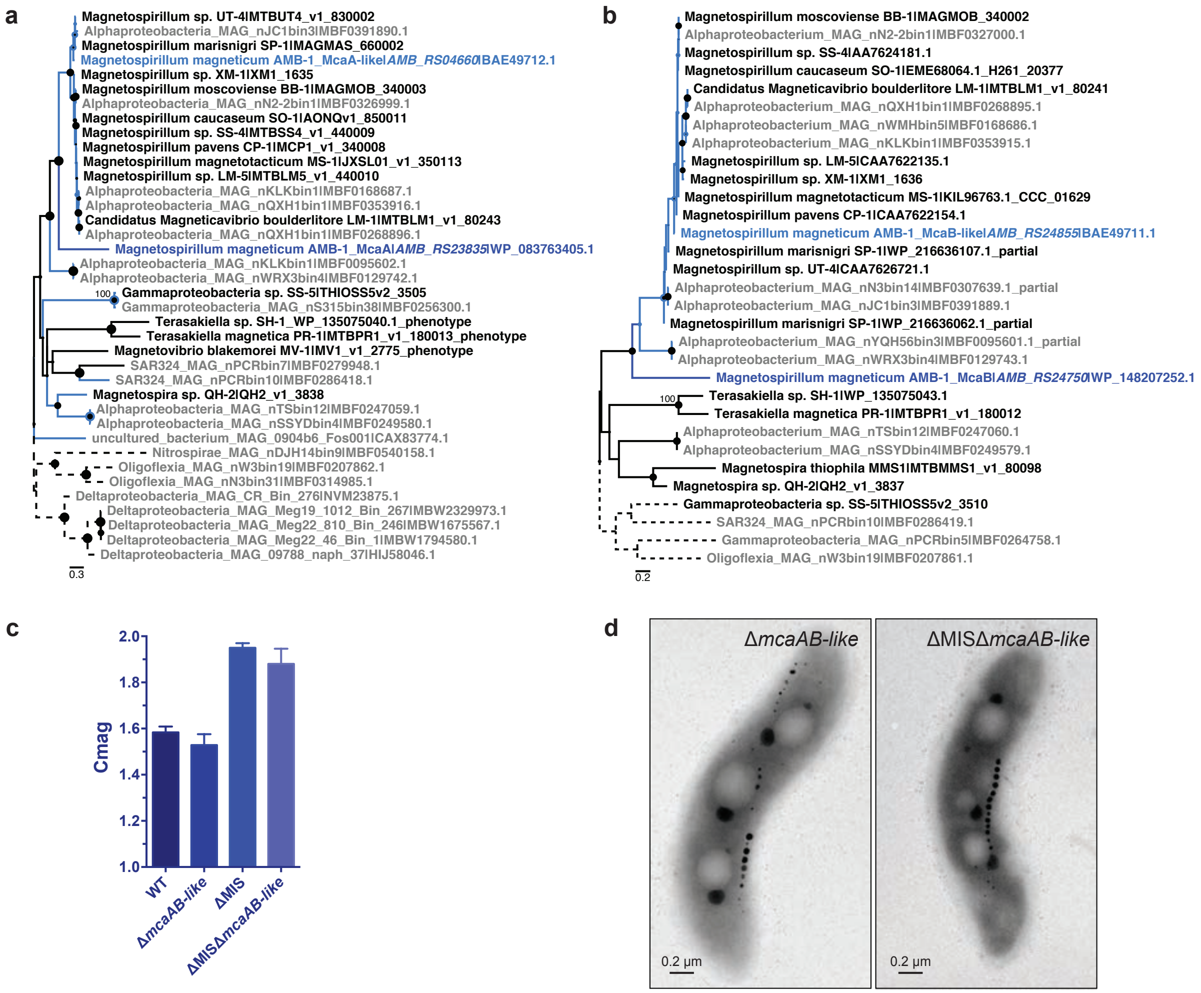


**Supplementary Figure 16: Comparative genomic analysis of McaA and McaB homologs and genetic analysis of *mcaAB*-like genes.** (a) and (b). Maximum likelihood trees showing the diversity of amino acid sequences homologous to McaA (**a**) and McaB (**b**) and their relationships. All sequences detected belong either to genomes of magnetotactic cultivated strains (black, described in Monteil et al. [5], Du et al. [6]), either to metagenome-assembled genomes (MAG) putatively magnetotactic (grey, Lin et al.[7]). Each strain/MAG sequence is associated to an accession number in NCBI or Microscope public databases to which it is separated by a pipe. Dotted branches refer to external groups while coloured clades refer to the two clusters of homologous sequences identified by MMseqs2 to which belong McaA and McaB of *Magnetospirillum magneticum* AMB-1 of the islet (dark blue), and their distant homologs Mca-like detected AMB-1 and other magnetotactic *Rhodospirillaceae* genomes (light blue). For these sequences, locus_names in the latest version of the annotated genome are in italics. Trees were drawn to scale and branch length refers to the numbers of substitution per site. Robustness of the internal branches is symbolized by a circle whose size is proportional to the bootstrap value estimated from 500 non-parametric replicates. Trees were rooted with sequences detected in the deepest branching phyla according to current standard phylogenies [8], i.e. *Desulfobacterota*/ *Bdellovibrionota* and non-*Alphaproteobacteria* respectively. (c) Cmag of WT, Δ*mcaAB*-like, ΔMIS, and ΔMISΔ*mcaAB*-like cultures. Each measurement represents the average and standard deviation from three independent growth cultures. (d) TEM micrographs of Δ*mcaAB*-like and ΔMISΔ*mcaAB*-like cells.

**Supplementary methods**

**Plasmids generation**

Plasmid pAK1036 was constructed to express the fusion protein MamI-Halo under Tac promoter. Firstly, *mmsF* and HL4-Linker were PCR amplified using the primers listed in Supplementary Table 11 and cloned into pAK979 [9] (digested with EcoRI) by Gibson assembly to generate pAK1032. Then, HL4 linker-*halo* was PCR amplified from pAK1032 using the primers listed in Supplementary Table 11 and inserted into pAK976 [9] (digested with EcoRI and SpeI) by Gibson assembly to generate pAK1034. Finally, the *mamI* gene was PCR amplified from the AMB-1 genomic DNA using the primers listed in Supplementary Table 11 and cloned into pAK1034 (digested with EcoRI) by Gibson assembly to generate pAK1036.

Plasmid pAK1037 was constructed to create the in-frame deletion of *mamJ-like* gene by homologous recombination. The *mamJ-like* upstream and downstream regions (about 800-1000 bp) were PCR amplified and inserted into the suicide vector pAK31 (digested with SpeI) using Gibson assembly. All the primers used to generate deletion plasmids in this study are listed in Supplementary Table 8. All the other deletion plasmids were generated similarly to pAK1037. Plasmid pAK1121 was constructed to delete the whole MIS region. Plasmid pAK1151 was constructed to delete the large domain 1 (LD1) of the MIS region (from the beginning of MIS to *mamJ-like*). The upstream region of MIS and the downstream region of *mamJ-like* were amplified and inserted into pAK31 to create pAK1151. Plasmid pAK1152 was constructed to delete the large domain 2 (LD1) of the MIS region (from *mamJ-like* to the end of MIS). The upstream region of *mamJ-like* and the downstream region of MIS were amplified and inserted into pAK31 to create pAK1152. Plasmids pAK1188, pAK1189, pAK1190, and pAK1191 were constructed to delete the iR1, iR2, iR3, and iR4 regions of MIS, respectively. Plasmids pAK1224, pAK1225, and pAK1277 were constructed to delete the genes of *mcaB*, *mcaA*, and *mamY*, respectively.

Plasmid pAK1101 was constructed to express the fusion protein Mms6-Halo under the Tac promoter. The *mms6* gene was PCR amplified from the AMB-1 genomic DNA using the primers listed in Supplementary Table 11 and cloned into pAK1034 (digested with EcoRI) by Gibson assembly.

Plasmid pAK1102 was constructed to express the fusion protein Mms6-GFP under the Tac promoter. The *mms6* gene was PCR amplified from the AMB-1 genomic DNA using the primers listed in Supplementary Table 11 and cloned into pAK22 [10] (double digested with EcoRI and BamHI) by Gibson assembly.

Plasmid pAK1195 was constructed to express the fusion protein Mms6-GFP under Tac promoter with gentamycin resistance. The vector was PCR amplified from pAK1102 (except the kanamycin gene), and the gentamycin gene was PCR amplified from the plasmid pJN105, then the two DNA fragments were assembled by Gibson cloning. The gentamycin gene will use the kanamycin promoter.

Plasmid pAK1199 was constructed to complement the whole iR2 region deletion mutant. The whole iR2 region was PCR amplified from the AMB-1 genomic DNA using the primers listed in Supplementary Table 11, and cloned into pAK22 (double digested with EcoRI and SpeI) by Gibson assembly.

To construct pAK1200 (*gfp-mcaA*) and pAK1237 (*gfp-mcaB*), *ambRS23835* (*mcaA*) and *ambRS24750* (*mcaB*) were PCR amplified from the AMB-1 genomic DNA, and inserted into pAK532 (digested with BamHI and SpeI) using Gibson assembly. To construct pAK1201 (*mcaA-gfp*) and pAK1238 (*mcaB-gfp*), *mcaA* and *mcaB* were PCR amplified from the AMB-1 genomic DNA and inserted into pAK22 (digested with EcoRI and BamHI) using Gibson assembly.

To generate pAK1240-pAK1252, *mcaA*, *mcaB*, and *mamY* were PCR amplified using the primers listed in Supplementary Table 10 and were cloned into pKT25 or pKNT25 (the N or the C-termini of the T25 fragment) and pUT18 or pUT18C (the N or the C-termini of the T18 fragment) vectors in frame with the T25 and T18 fragment open reading frames by Gibson assembly.

Plasmid pAK1255 was constructed to express the fusion proteins of McaB-GFP and McaA-Halo under Tac promoter in the same cell. The *mcaA* gene and its ribosome binding site (*rbs-mcaA*) were PCR amplified from the AMB-1 genomic DNA, *halo* gene was PCR amplified from pAK1036, then both PCR fragments were inserted into pAK1238 (digested with SpeI) using Gibson assembly.

Plasmid pAK1256 was constructed to express the fusion proteins of McaB-GFP and McaA-Halo under the native promoter of the *mcaAB* operon in the same cell. *mcaB* with its native promoter region and *rbs-mcaA* were PCR amplified from the AMB-1 genomic DNA, *gfp* was PCR amplified from pAK22, then all of three PCR fragments were inserted into pAK1036 (digested with BamHI and HindII) using Gibson assembly.

Plasmids pAK1257-pAK1263 were constructed to express the mutated McaA that fused with GFP under Tac promoter. For pAK1257 and pAK1263, the N- and C- terminally truncated *mcaA* were PCR amplified from the AMB-1 genomic DNA, and inserted into pAK22 (digested with BamHI and EcoRI) using Gibson assembly. For pAK1258, pAK1260, pAK1261, and pAK1262, the upstream and downstream deleted regions of *mcaA* were PCR amplified from the AMB-1 genomic DNA, and inserted into pAK22 (digested with BamHI and EcoRI) using Gibson assembly. For pAK1259, the three conserved amino acids of the *mcaA* MIDAS motif (**Asp**-Xaa-**Ser**-Xaa-**Ser** (**D**X**S**X**S**), where X is any amino acid) were mutated into alanines, and the mutations were included in the primers. Then the fragments were PCR amplified from the AMB-1 genomic DNA and inserted into pAK22 (digested with BamHI and EcoRI) using Gibson assembly. All primers used to generate *mcaA* mutants were listed in Supplementary Table 12.

Plasmid pAK1270 was constructed to express the fusion proteins of McaB-GFP and Mms6-Halo under Tac promoter in the same cell. *rbs-mms6-halo* was PCR amplified from pAK1101 and inserted into pAK1255 (digested with SpeI) using Gibson assembly.

**Mutated strain generation**

The whole MIS region was deleted in WT (AK30), ΔMAI, and Δ*mamJ*Δ*limJ* genetic backgrounds by homologous recombination to generate the strains of ΔMIS, ΔMAIΔMIS, and Δ*mamJ*Δ*limJ*ΔMIS, respectively. The *mamJ-like* gene was deleted in WT and Δ*mamJ*Δ*limJ* genetic backgrounds to generate the strains of Δ*mamJ-like* and Δ*mamJ*Δ*limJ*Δ*mamJ-like*, respectively. The LD1 and LD2 regions of MIS were deleted in WT to create strains of ΔMIS_LD1 and ΔMIS_LD2, respectively. The iR1, iR2, iR3, and iR4 regions of MIS were deleted in WT to create strains of ΔiR1, ΔiR2, ΔiR3, and ΔiR4, respectively. The iR1, iR2, iR3, and iR4 regions were deleted in a Δ*mamJ*Δ*limJ* strain to create the triple deletions of Δ*mamJ*Δ*limJ*ΔiR1, Δ*mamJ*Δ*limJ*ΔiR2, Δ*mamJ*Δ*limJ*ΔiR3, Δ*mamJ*Δ*limJ*ΔiR4, respectively. The *mcaA* gene was in-frame deleted in WT and Δ*mamJ*Δ*limJ* genetic backgrounds to generate the strains of Δ*mcaA* and Δ*mamJ*Δ*limJ*Δ*mcaA*, respectively. The *mcaB* gene was in-frame deleted in WT and Δ*mamJ*Δ*limJ* genetic backgrounds to generate the strains of Δ*mcaB* and Δ*mamJ*Δ*limJ*Δ*mcaB*, respectively. The *mamY* gene was in-frame deleted in WT, ΔMIS, ΔiR2, Δ*mcaA*, and Δ*mcaB* genetic backgrounds to create the strains of Δ*mamY*, Δ*mamY*ΔMIS, Δ*mamY*ΔiR2, Δ*mamY*Δ*mcaA,* and Δ*mamY*Δ*mcaB*, respectively.

#### Supplementary Tables

### Supplementary Table 1. Colocalization quantification analysis of fluorescent signals in AMB-1 cells.

| **Fluorescent signals** | | **Strains** | **PCC** | **MCC (M1)** | **MCC (M2)** | **No. of cells** | **Corresponding figures** |
|---|---|---|---|---|---|---|---|
| MamI-Halo | JF549 (pulse, +Fe)/JF646 (chase, +Fe) | WT | 0.17±0.17 | 0.16±0.10 | 0.32±0.19 | 18 | Fig. 2d, S4a |
| | | ΔMIS | 0.18±0.22 | 0.11±0.07 | 0.47±0.30 | 15 | |
| Halo-MmsF | JF549 (pulse, +Fe)/JF646 (chase, +Fe) | WT | 0.56±0.20 | 0.67±0.20 | 0.66±0.16 | 13 | Fig. S5e, f |
| | | ΔMIS | 0.66±0.22 | 0.63±0.19 | 0.74±0.22 | 28 | |
| Mms6-Halo/McaB-GFP | | WT | 0.76±0.10 | 0.74±0.10 | 0.84±0.10 | 24 | Fig. 4e |
| McaA-Halo/McaB-GFP | | WT | 0.36±0.12 | 0.25±0.12 | 0.39±0.17 | 33 | Fig. 4f, S10d |
| MamI-Halo | JF549 (pulse, -Fe)/JF646 (chase, +Fe) | WT | 0.19±0.21 | 0.24±0.15 | 0.31±0.19 | 55 | Fig. 6c, S4b |
| | | ΔMIS | 0.05±0.27 | 0.14±0.11 | 0.39±0.24 | 32 | |
| | | ΔiR2 | 0.13±0.22 | 0.14±0.09 | 0.32±0.22 | 25 | |

PCC, Pearon’s Correlation Coefficient. PCC measures the pixel-by-pixel covariance in the signal levels of two images, and PCC is independent of signal levels and signal offset (background). PCC values range from 1 ( perfect positive correlation) to -1 (perfectly negative correlation). PCC values near 0 indicate no correlation.
MCC, Manders' Colocalization Coefficients. MCC strictly measures co-occurrence independent of signal proportionality, and shows the fraction of each protein that is colocalised with the other. MCC values range from 1 (100% pixel co-occurrence) to 0 (no pixel co-occurrence). M1: the percentage of above-background pixels in the JF549-stained/red channel that overlap above-background pixels in the JF646-stained/green channel. M2: the percentage of above-background pixels in the JF646-stained/green channel that overlap above-background pixels in the JF549-stained/red channel. Values indicate mean ± standard deviation.

#### Supplementary Table 2. Predicted and experimentally determined topologies for McaA and McaB

<table>
<tr><th>Protein</th><th colspan="3">Membrane topology prediction: CCTOP</th><th>Signal peptide prediction: Signal 4.1 and Phobius</th><th>Domain prediction: SMART</th><th>Domain prediction: InterProScan</th><th>N-ter/C-ter (determined experimentally)</th></tr>
<tr><td rowspan="3">McaA (776 aa)</td><td rowspan="3">4 TM, N-ter (C)/ C-ter (C)</td><td>4 TM (aa7-22, aa371-390, aa558-573, aa747-762), N-ter (C)/ C-ter (C)</td><td>HMMTOP, Philius, Scampi, ScampiMsa</td><td rowspan="3">Both Yes, aa1-26, Cleavage site between aa26 and aa27</td><td rowspan="3">1 VWA domain (aa29-258),<br>2 TM (aa7-26 and aa369-391)</td><td rowspan="3">1 VWA domain (aa31-212), signal peptide (aa 1-26)/2 TM (aa7-26 and aa369-391).<br><br>aa27-370 (P) and aa392-776 (C).</td><td rowspan="3">GFP-McaA complements ΔmcaA, but does not fluorescent, McaB-GFP does complement ΔmcaA and fluorescent:<br>N-ter (P)/C-ter (C)</td></tr>
<tr><td>2 TM (aa7-22, aa371-390), N-ter (C)/ C-ter (C)</td><td>Pro, TMHMM</td></tr>
<tr><td>1 TM (aa371-390), N-ter (P)/ C-ter (C)</td><td>Memsat, Phobius (with N-ter SP)</td></tr>
<tr><td rowspan="2">McaB (219 aa)</td><td rowspan="2">1 TM, N-ter (P)/ C-ter (C)</td><td>1 TM (5-27), N-ter (P)/ C-ter (C)</td><td>HMMTOP, Memsat, Octopus, Pro, Prodiv, ScampiMsa</td><td rowspan="2">No</td><td rowspan="2">1 TM (aa 5-27)</td><td rowspan="2">1 TM (aa 5-27), signal peptide (aa 1-20), coil (82-102), aa21-219 (P)</td><td rowspan="2">GFP-McaB does not fluorescent, McaB-GFP does complement ΔmcaB and fluorescent:<br>N-ter (P)/C-ter (C)</td></tr>
<tr><td>1 TM (5-27), N-ter (C)/ C-ter (P)</td><td>Philius, Scampi, TMHMM</td></tr>
</table>

Constrained Consensus Topology prediction server (CCTOP) integrates the prediction from several prediction methods: HMMTOP, Membrain, Memsat-SVM, Octopus, Philius, Phobius, Pro, Prodiv, Scampi, and TMHMM. C, cytoplasmic; P, periplasmic; TM, transmembrane; SP, signal peptide.

#### Supplementary Table 3. Fluorescence recovery after photobleaching

| **Construct** | **Strain** | $t_{1/2}$[a] | **% Recovery[b] of the cells with the bleached region does not move** | **No. of cells with the bleached region move** | **Total no. of cells[c]** |
|---|---|---|---|---|---|
| *mamK-gfp* | WT AMB-1 | 4.9 ± 3.4 | 100 | 0 | 23 |
| *mamK-gfp* | ΔMIS | 4.2 ± 2.8 | 91 | 15 | 37 |
| *mamK-gfp* | Δ*mcaA* | 3.7 ± 1.9 | 100 | 8 | 36 |
| *mamK-gfp* | Δ*mcaB* | 3.2 ± 1.9 | 100 | 10 | 34 |

a. Half-time of recovery was measured in minutes. Shown is the average ± standard deviation.
b. Percentage of cells whose bleached regions regained at least half of the overall filament fluorescence.
c. Total number of cells observed, composed of recovering and non-recovering cells.

#### Supplementary Table 4. Coordinates and magnetic phenotype of the MIS mutants

| **Name of region** | **Genes included in deletion** | **Coordinates** | **Magnetic phenotype** |
|---|---|---|---|
| MAI | (Old tag: *amb0933* to *amb1031*) *amb_RS04800* to *amb_RS25575* | 996,982 - 1,096,635 | NO |
| MIS | *amb_RS02005* to *amb_RS23870* | 421,600 - 450,000 | Much higher than WT |
| LD1 | *amb_RS02005* to *amb_RS24775* | 421,600 - 440,560 | ΔMIS |
| LD2 | *amb_RS24775* to *amb_RS23870* | 439,811 - 450,000 | WT |
| iR1 | *amb_RS02005* to *amb_RS23830* | 421,600 - 426,000 | WT |
| iR2 | *amb_RS23835* to *amb_RS26010* | 426,000 - 431,000 | ΔMIS |
| iR3 | *amb_RS23840* to *amb_RS23850* | 429,699 - 437,000 | Slightly higher than WT |
| iR4 | *amb_RS23850* to *amb_RS02100* | 437,000 - 439,750 | WT |
| *MamJ-like* | *amb_RS24775* | 440,560 - 439,811 | WT |
| *mcaA* | *amb_RS23835* | 426,172 - 428,502 | ΔMIS |
| *mcaB* | *amb_RS24750* | 428,523 - 429,182 | Much higher than WT but lower than ΔMIS |
| Potential transposase in iR2 region | *amb_RS23840* | 429,860 - 430,701 | - |

Magnetic phenotypes were determined by Cmag measurement. NO, cells are nonmagnetic; WT, similar to wild type.

#### Supplementary Table 5. Strains used in this study

| Strain | Organism | Description | Reference |
|---|---|---|---|
| AK30 | *M. magneticum* AMB-1 | Wild-type AMB-1 | [11] |
| AK31 | *M. magneticum* AMB-1 | ΔMAI | [11] |
| AK108 | *M. magneticum* AMB-1 | Δ*mamJ*Δ*limJ* | [12] |
| AK271 | *M. magneticum* AMB-1 | ΔMIS | This work |
| AK272 | *M. magneticum* AMB-1 | ΔMAIΔMIS | This work |
| AK273 | *M. magneticum* AMB-1 | Δ*mamJ*Δ*limJ*ΔMIS | This work |
| AK274 | *M. magneticum* AMB-1 | Δ*mamJ-like* | This work |
| AK275 | *M. magneticum* AMB-1 | Δ*mamJ*Δ*limJ*Δ*mamJ-like* | This work |
| AK299 | *M. magneticum* AMB-1 | ΔMIS_LD2 | This work |
| AK316 | *M. magneticum* AMB-1 | ΔMIS_LD1 | This work |
| AK321 | *M. magneticum* AMB-1 | Δ*mamJ*Δ*limJ*ΔiR1 | This work |
| AK322 | *M. magneticum* AMB-1 | Δ*mamJ*Δ*limJ*ΔiR2 | This work |
| AK323 | *M. magneticum* AMB-1 | Δ*mamJ*Δ*limJ*ΔiR3 | This work |
| AK324 | *M. magneticum* AMB-1 | Δ*mamJ*Δ*limJ*ΔiR4 | This work |
| AK325 | *M. magneticum* AMB-1 | ΔiR1 | This work |
| AK326 | *M. magneticum* AMB-1 | ΔiR2 | This work |
| AK327 | *M. magneticum* AMB-1 | ΔiR3 | This work |
| AK328 | *M. magneticum* AMB-1 | ΔiR4 | This work |
| AK331 | *M. magneticum* AMB-1 | Δ*mcaB* | This work |
| AK332 | *M. magneticum* AMB-1 | Δ*mamJ*Δ*limJ*Δ*mcaB* | This work |
| AK333 | *M. magneticum* AMB-1 | Δ*mcaA* | This work |
| AK334 | *M. magneticum* AMB-1 | Δ*mamJ*Δ*limJ*Δ*mcaA* | This work |
| AK339 | *M. magneticum* AMB-1 | Δ*mamY* | This work |
| AK340 | *M. magneticum* AMB-1 | Δ*mamY*ΔMIS | This work |
| AK348 | *M. magneticum* AMB-1 | Δ*mamY*ΔiR2 | This work |
| AK349 | *M. magneticum* AMB-1 | Δ*mamY*Δ*mcaA* | This work |
| AK351 | *M. magneticum* AMB-1 | Δ*mamY*Δ*mcaB* | This work |
| DH5α (λpir) | *E. coli* | Standard cloning strain | [11] |
| WM3064 | *E. coli* | Conjugation strain | [11] |
| DHM1 | *E. coli* | Reporter strain for BACTH assay | Euromedex |
| XL1-Blue | *E. coli* | standard *E. coli* K12 recA strain | Agilent |

#### Supplementary Table 6. Plasmids used or generated in this study (except for the BACTH)

| Name | Purpose | Origin | Reference |
|---|---|---|---|
| pAK272 | C-terminal GFP fusion to MamI | pAK22 | This work |
| pAK532 | N-terminal GFP fusion to MmsF | pAK22 | [1] |
| pAK983 | N-terminal Halo fusion to MmsF | pAK978 | [9] |
| pAK1032 | C-terminal Halo fusion to MmsF | pAK979 | This work |
| pAK1034 | HL4-Linker halo | pAK976 | This work |
| pAK1036 | C-terminal Halo fusion to MamI | pAK22 | This work |
| pAK1037 | *mamJ-like* deletion | pAK31 | This work |
| pAK1101 | C-terminal Halo fusion to Mms6 | pAK22 | This work |
| pAK1102 | C-terminal GFP fusion to Mms6 (Kanamycin) | pAK22 | This work |
| pAK1121 | MIS deletion | pAK31 | This work |
| pAK1151 | LD1 deletion | pAK31 | This work |
| pAK1152 | LD2 deletion | pAK31 | This work |
| pAK1188 | iR1 deletion | pAK31 | This work |
| pAK1189 | iR2 deletion | pAK31 | This work |
| pAK1190 | iR3 deletion | pAK31 | This work |
| pAK1191 | iR4 deletion | pAK31 | This work |
| pAK1195 | C-terminal GFP fusion to Mms6 (Gentamicin) | pJN105 | This work |
| pAK1199 | iR2 complementation | pAK22 | This work |
| pAK1200 | N-terminal GFP fusion to McaA | pAK532 | This work |
| pAK1201 | C-terminal GFP fusion to McaA | pAK22 | This work |
| pAK1224 | *mcaB* deletion | pAK31 | This work |
| pAK1225 | *mcaA* deletion | pAK31 | This work |
| pAK1226 | pBBR1MCS-2 (empty vector) | pAK31 | This work |
| pAK1237 | N-terminal GFP fusion to McaB | pAK1200 | This work |
| pAK1238 | C-terminal GFP fusion to McaB | pAK1201 | This work |
| pAK1255 | Tac_McaB-GFP_rbs-McaA-Halo | pAK1038 | This work |
| pAK1256 | Own promoter McaB-GFP_rbs-McaA-Halo | pAK1036 | This work |
| pAK1257 | McaA-mutant-1(aa1-28 deletion) | pAK22 | This work |
| pAK1258 | McaA-mutant-2(aa29-258 deletion) | pAK22 | This work |
| pAK1259 | McaA-mutant-3(MIADS motif mutation) | pAK22 | This work |
| pAK1260 | McaA-mutant-4(aa259-400 deletion) | pAK22 | This work |
| pAK1261 | McaA-mutant-5(aa400-530 deletion) | pAK22 | This work |
| pAK1262 | McaA-mutant-6(aa530-665 deletion) | pAK22 | This work |
| pAK1263 | McaA-mutant-7(aa665-776 deletion) | pAK22 | This work |
| pAK1270 | Tac_McaB-GFP_rbs-Mms6-Halo | pAK1255 | This work |
| pAK1277 | *mamY* deletion | pAK31 | This work |

#### Supplementary Table 7. Plasmids for BACTH

| Name | Purpose | Origin | Reference |
|---|---|---|---|
| pAK318 | pKT25 | pSU40 | Euromedex |
| pAK319 | pKNT25 | pSU40 | Euromedex |
| pAK320 | pUT18 | pUC19 | Euromedex |
| pAK321 | pUT18C | pUC19 | Euromedex |
| pAK322 | pKT25-zip | pKT25 | Euromedex |
| pAK323 | pUT18C-zip | pUT18C | Euromedex |
| pAK324 | pKT25-*mamK* | pKT25 | [13] |
| pAK325 | pKNT25-*mamK* | pKNT25 | [13] |
| pAK326 | pUT18-*mamK* | pUT18 | [13] |
| pAK327 | pUT18C-*mamK* | pUT18C | [13] |
| pAK816 | pUT18-*mamJ* | pUT18 | [13] |
| pAK818 | pKT25-*mamJ* | pKT25 | [13] |
| pAK1240 | pKT25-*mcaA* | pKT25 | This work |
| pAK1241 | pKNT25-*mcaA* | pKNT25 | This work |
| pAK1242 | pUT18-*mcaA* | pUT18 | This work |
| pAK1243 | pUT18C-*mcaA* | pUT18C | This work |
| pAK1245 | pKT25-*mamY* | pKT25 | This work |
| pAK1246 | pKNT25-*mamY* | pKNT25 | This work |
| pAK1247 | pUT18-*mamY* | pUT18 | This work |
| pAK1248 | pUT18C-*mamY* | pUT18C | This work |
| pAK1249 | pKT25-*mcaB* | pKT25 | This work |
| pAK1250 | pKNT25-*mcaB* | pKNT25 | This work |
| pAK1251 | pUT18-*mcaB* | pUT18 | This work |
| pAK1252 | pUT18C-*mcaB* | pUT18C | This work |

### Supplementary Table 8. List of primers for generating deletion plasmids

| Name | Sequence | Target | In plasmid |
|---|---|---|---|
| Δ*mamJ-like*-A | Fwd: 5'-gaattcctgcagcccgggggatcc***ACTAGT***gatggacatgatggtccaggg-3' | *mamJ-like* upstream region | pAK1037 |
| Δ*mamJ-like*-B | Rev: 5'-CCCATCCACTAAATTTAAATAtgatgatttgcatgctcatcg-3' | | |
| Δ*mamJ-like*-C | Fwd: 5'-TATTTAAATTTAGTGGATGGGcacctcactggtaacagcagc-3' | *mamJ-like* downstream region | |
| Δ*mamJ-like*-D | Rev: 5'-caccgcggtggcggccgctctaga***ACTAGT***gacttggcatactccgagaag-3' | | |
| ΔMIS-A | Fwd: 5'-gaattcctgcagcccgggggatcc***ACTAGT***gttcccctccatcacctatac-3' | MIS upstream region | pAK1121 |
| ΔMIS -B | Rev: 5'-CCCATCCACTAAATTTAAATAagcggcggtatagcccatg-3' | | |
| ΔMIS -C | Fwd: 5'-TATTTAAATTTAGTGGATGGGctattcgaaccgcctgctc-3' | MIS downstream region | |
| ΔMIS -D | Rev: 5'-caccgcggtggcggccgctctaga***ACTAGT***gatagcgagaaccgtcatac-3' | | |
| ΔiR1-A | Fwd: 5'-gaattcctgcagcccgggggatcc***ACTAGT***gttcccctccatcacctatac-3' | iR1 upstream region | pAK1188 |
| ΔiR1-B | Rev: 5'-CCCATCCACTAAATTTAAATAagcggcggtatagcccat-3' | | |
| ΔiR1-C | Fwd: 5'-TATTTAAATTTAGTGGATGGGgttgggtcgtgtcccgcc-3' | iR1 downstream region | |
| ΔiR1-D | Rev: 5'-caccgcggtggcggccgctctaga***ACTAGT***gttgttcgttgctgctcc-3' | | |
| ΔiR2-A | Fwd: 5'-gaattcctgcagcccgggggatcc***ACTAGT***atatcgctccagaatggaattc-3' | iR2 upstream region | pAK1189 |
| ΔiR2-B | Rev: 5'-CCCATCCACTAAATTTAAATAcaccatggaggccgattt-3' | | |
| ΔiR2-C | Fwd: 5'-TATTTAAATTTAGTGGATGGGcaatctgcgctgcaggct-3' | iR2 downstream region | |
| ΔiR2-D | Rev: 5'-caccgcggtggcggccgctctaga***ACTAGT***agcatgcttcggccccct-3' | | |
| ΔiR3-A | Fwd: 5'-cctgcagcccgggggatcc***ACTAGT***atataagactgatttttcgat-3' | iR3 upstream region | pAK1190 |
| ΔiR3-B | Rev: 5'-CCCATCCACTAAATTTAAATAtccccaatagcggccatg-3' | | |
| ΔiR3-C | Fwd: 5'-TATTTAAATTTAGTGGATGGGcgggccagctcgatccgc-3' | iR3 downstream region | |
| ΔiR3-D | Rev: 5'-caccgcggtggcggccgctctaga***ACTAGT***ccatgggatcgcgccaca-3' | | |
| ΔiR4-A | Fwd: 5'-gaattcctgcagcccgggggatcc***ACTAGT***ccgccatgtacaccctgatc-3' | iR4 upstream region | pAK1191 |
| ΔiR4-B | Rev: 5'-CCCATCCACTAAATTTAAATAgtttgcgccggatgcagt-3' | | |
| ΔiR4-C | Fwd: 5'-TATTTAAATTTAGTGGATGGGgcacctcaacggtcgtga-3' | iR4 downstream region | |
| ΔiR4-D | Rev: 5'-caccgcggtggcggccgctctaga***ACTAGT***gcccccccctgaaatactag-3' | | |
| Δ*mcaB*-A | Fwd: 5'-gaattcctgcagcccgggggatcc***ACTAGT***gatatttcgctcctgaag-3' | *mcaB* upstream region | pAK1224 |
| Δ*mcaB*-B | Rev: 5'-CCCATCCACTAAATTTAAATAgtcgatcacggaggaacg-3' | | |
| Δ*mcaB*-C | Fwd: 5'-TATTTAAATTTAGTGGATGGGcatcccctccactgccatag-3' | *mcaB* downstream region | |
| Δ*mcaB*-D | Rev: 5'-caccgcggtggcggccgctctaga***ACTAGT***ccaaagcgccaagacccaaac-3' | | |
| Δ*mcaA*-A | Fwd: 5'-gaattcctgcagcccgggggatcc***ACTAGT***ccagcttgagtaagagctttttg-3' | *mcaA* upstream region | pAK1225 |
| Δ*mcaA*-B | Rev: 5'-CCCATCCACTAAATTTAAATAggagagcagttaaggagg-3' | | |
| Δ*mcaA*-C | Fwd: 5'-TATTTAAATTTAGTGGATGGGtacgttcctccgtgatcg-3' | *mcaA* downstream region | |
| Δ*mcaA*-D | Rev: 5'-caccgcggtggcggccgctctaga***ACTAGT***ccttgatgtaagtgctac-3' | | |
| Δ*mamY*_A | Fwd: 5'-gaattcctgcagcccgggggatcc***ACTAGT***cagggcgcccacggtcat-3' | *mamY* upstream region | pAK1277 |
| Δ*mamY*_B | Rev: 5'-CCCATCCACTAAATTTAAATAgattccggcatggactgatgg-3' | | |
| Δ*mamY*_C | Fwd: 5'-TATTTAAATTTAGTGGATGGGgatggccgcaatcgccatccc-3' | *mamY* downstream region | |
| Δ*mamY*_D | Rev: 5'-caccgcggtggcggccgctctaga***ACTAGT***gacatctcgaacagcagcttggc-3' | | |

#### Supplementary Table 9. List of primers for verification of deletion mutant strains

| Name | Sequence | Target |
|---|---|---|
| ΔMAI-Confirm-Fwd | 5'-cattgcagcacccatcaccac-3' | MAI |
| ΔMAI-Confirm-Rev | 5'-gaattcctcatggcgccgaag-3' | |
| ΔMIS-Confirm-Fwd | 5'-cattgcagcacccatcaccac-3' | MIS |
| ΔMIS-Confirm-Rev | 5'-gaattcctcatggcgccgaag-3' | |
| Δ*mamJ-like*-Confirm-Fwd | 5'-caattactagaaatccaactaggg-3' | *mamJ-like* |
| Δ*mamJ-like*-Confirm-Rev | 5'- ccatcaaattattggagctgtatc-3' | |
| ΔiR1-Confirm-Fwd | 5'-gacacgatgagagtcacg-3' | iR1 |
| ΔiR1-Confirm-Rev | 5'-catctggcgtgatcgatg-3' | |
| ΔiR2-Confirm-Fwd | 5'-cagatcgaaactcgcagg-3' | iR2 |
| ΔiR2-Confirm-Rev | 5'-ccatttccggctgcagtc-3' | |
| ΔiR3-Confirm-Fwd | 5'-gcttctgtgcatcactgg-3' | iR3 |
| ΔiR3-Confirm-Rev | 5'-cgagttgaggccaagtgc-3' | |
| ΔiR4-Confirm-Fwd | 5'-gattatctgctgaagcgc-3' | iR4 |
| ΔiR4-Confirm-Rev | 5'-cctggttaccgcttatcc-3' | |
| Δ*mcaA*-Confirm-Fwd | 5'-gaattcggcctttcatgg-3' | *mcaA* |
| Δ*mcaA*-Confirm-Rev | 5'-gtgcgcatcactgctgtc-3' | |
| Δ*mcaB*-Confirm-Fwd | 5'-gacataaaatccgacacc-3' | *mcaB* |
| Δ*mcaB*-Confirm-Rev | 5'-ggctaccttggagaaaag-3' | |
| Δ*mamY*-Confirm-Fwd | 5'-gtcagatagaacagcacc-3' | *mamY* |
| Δ*mamY*-Confirm-Rev | 5'-gaatcctcgatcagggatg-3' | |

#### Supplementary Table 10. List of primers for making BACTH plasmids

| Name | Sequence | Target | In plasmid |
|---|---|---|---|
| JW-1240-Fwd | 5'-gcgggctgcagggtcgactctaga***GGATCC***cgtaattcgaggggggggg-3' | pKT25-*mcaA* | pAK1240 |
| JW-1240-Rev | 5'-tcacgacgttgtaaaacgacggcc***GAATTC***tcactttttgaaaatggc-3' | | |
| JW-1241-Fwd | 5'-aacagctatgaccatgattacgcc***AAGCTT***ggtggtaattcgagggggg-3' | pKNT25-*mcaA* | pAK1241 |
| JW-1241-Rev | 5'-attgaattcgagctcggtacccgg***GGATCC***tcctttttgaaaatggcattc-3' | | |
| JW-1242-Fwd | 5'-aacagctatgaccatgattacgcc***AAGCTT***ggtggtaattcgagggggg-3' | pUT18-*mcaA* | pAK1242 |
| JW-1242-Rev | 5'-gctgaattcgagctcggtacccgg***GGATCC***tcctttttgaaaatggcattc-3' | | |
| JW-1243-Fwd | 5'-acgccactgcaggtcgactctaga***GGATCC***cgtaattcgaggggggggg-3' | pUT18C-*mcaA* | pAK1243 |
| JW-1243-Rev | 5'-accatattacttagttatatcgat***GAATTC***tcactttttgaaaatggc-3' | | |
| JW-1245-Fwd | 5'-gcgggctgcagggtcgactctaga***GGATCC***cgcgattgcggccatcatg-3' | pKT25-*mamY* | pAK1245 |
| JW-1245-Rev | 5'-tcacgacgttgtaaaacgacggcc***GAATTC***tcagtccatgccggaatc-3' | | |
| JW-1246-Fwd | 5'-aacagctatgaccatgattacgcc***AAGCTT***gatggcgattgcggccatc-3' | pKNT25-*mamY* | pAK1246 |
| JW-1246-Rev | 5'-attgaattcgagctcggtacccgg***GGATCC***tcgtccatgccggaatcggg-3' | | |
| JW-1247-Fwd | 5'-aacagctatgaccatgattacgcc***AAGCTT***gatggcgattgcggccatc-3' | pUT18-*mamY* | pAK1247 |
| JW-1247-Rev | 5'-gctgaattcgagctcggtacccgg***GGATCC***tcgtccatgccggaatcggg-3' | | |
| JW-1248-Fwd | 5'-acgccactgcaggtcgactctaga***GGATCC***cgcgattgcggccatcatg-3' | pUT18C-*mamY* | pAK1248 |
| JW-1248-Rev | 5'-accatattacttagttatatcgat***GAATTC***tcagtccatgccggaatc-3' | | |
| JW-1249-Fwd | 5'-gcgggctgcagggtcgactctaga***GGATCC***cattgaactggtcgtactc-3' | pKT25-*mcaB* | pAK1249 |
| JW-1249-Rev | 5'-tcacgacgttgtaaaacgacggcc***GAATTC***tcactcgagcttagaaag-3' | | |
| JW-1250-Fwd | 5'-aacagctatgaccatgattacgcc***AAGCTT***gatgattgaactggtcgtac-3' | pKNT25-*mcaB* | pAK1250 |
| JW-1250-Rev | 5'-attgaattcgagctcggtacccgg***GGATCC***tcctcgagcttagaaaggattattg-3' | | |
| JW-1251-Fwd | 5'-aacagctatgaccatgattacgcc***AAGCTT***gatgattgaactggtcgtac-3' | pUT18-*mcaB* | pAK1251 |
| JW-1251-Rev | 5'-gctgaattcgagctcggtacccgg***GGATCC***tcctcgagcttagaaaggattattg-3' | | |
| JW-1252-Fwd | 5'-acgccactgcaggtcgactctaga***GGATCC***cattgaactggtcgtactc-3' | pUT18C-*mcaB* | pAK1252 |
| JW-1252-Rev | 5'-accatattacttagttatatcgat***GAATTC***tcactcgagcttagaaag-3' | | |

#### Supplementary Table 11. List of primers for generating GFP/Halo fusion or complementation plasmids

| Name | Sequence | Target | In plasmid |
|---|---|---|---|
| JW-1032-a-Fwd | 5'-gacccccgggttggaggaataac***GAATTC***atgactgaagctatccttcgc-3' | *mmsF* | pAK1032 |
| JW-1032-a-Rev | 5'-ttagccgccgcggcctcggccag***GGATCC***gatccggtgggcgaccca-3' | | |
| JW-1032-b-Fwd | 5'-gctgggtcgcccaccggatc***GGATCC***ctggccgaggccgcggcg-3' | HL4 linker | |
| JW-1032-b-Rev | 5'-aagccagtaccgatttctgc***GGATCC***cgctgctgctttggccgccg-3' | | |
| JW-1034-Fwd | 5'-gataacaatttcacacaggaaaca***GAATTC***ctggccgaggccgcggcg-3' | linker-*halo* | pAK1034 |
| JW-1034-Rev | 5'-caccgcggtggcggccgctctaga***ACTAGT***ctagccggaaatctcgagcgt-3' | | |
| JW-1036-Fwd | 5'-gataacaatttcacacaggaaaca***GAATTC***atgccaagcgtgattttc-3' | *mamI* | pAK1036 |
| JW-1036-Rev | 5'-tttagccgccgcggcctcggccag***GAATTC***accatcgatgtcagggtc-3' | | |
| JW-1101-Fwd | 5'-gataacaatttcacacaggaaaca***GAATTC***atgccagctcagatcgccaac-3' | *mms6* | pAK1101 |
| JW-1101-Rev | 5'-tttagccgccgcggcctcggccag***GAATTC***ggccagcgcgtcgcgcag-3' | | |
| JW-1102-Fwd | 5'-gataacaatttcacacaggaaaca***GAATTC***gtgccagctcagatcgccaacgg-3' | *mms6* | pAK1102 |
| JW-1102-Rev | 5'-gaaaagttcttctcctttactcat***GGATCC***ggccagcgcgtcgcgcag-3' | | |
| JW-1195-a-Fwd | 5'-gcgggactctggggttcgaaatg-3' | pAK1102 except for the kanamycin gene | pAK1195 |
| JW-1195-a-Rev | 5'-gcgaaacgatcctcatcctgtc-3' | | |
| JW-1195-b-Fwd | 5'-gagacaggatgaggatcgtttcgcATGTTACGCAGCAGCAAC-3' | gentamycin gene | |
| JW-1195-b-Rev | 5'-tcatttcgaaccccagagtcccgcTTAGGTGGCGGTACTTGG-3' | | |
| JW-1199-Fwd | 5'-ggataacaatttcacacaggaaaca***GAATTC***gttgggtcgtgtcccgcc-3' | Whole iR2 region | pAK1199 |
| JW-1199-Rev | 5'-ccaccgcggtggcggccgctctaga***ACTAGT***gtgcccgcgactaatgcc-3' | | |
| JW-1200-Fwd | 5'-gaagcggcggccaaagcagcagcg***GGATCC***gtaattcgagggggggggg-3' | *mcaA* | pAK1200 |
| JW-1200-Rev | 5'-caccgcggtggcggccgctctaga***ACTAGT***tcactttttgaaaatggc-3' | | |
| JW-1201-Fwd | 5'-gataacaatttcacacaggaaaca***GAATTC***gtggtaattcgaggggggg-3' | *mcaA* | pAK1201 |
| JW-1201-Rev | 5'-gaaaagttcttctcctttactcat***GGATCC***ctttttgaaaatggcattc-3' | | |
| JW-1237-Fwd | 5'-gaagcggcggccaaagcagcagcg***GGATCC***attgaactggtcgtactc-3' | *mcaB* | pAK1237 |
| JW-1237-Rev | 5'-caccgcggtggcggccgctctaga***ACTAGT***ctactcgagcttagaaag-3' | | |
| JW-1238-Fwd | 5'-gataacaatttcacacaggaaaca***GAATTC***atgattgaactggtcgtac-3' | *mcaB* | pAK1238 |
| JW-1238-Rev | 5'-gaaaagttcttctcctttactcat***GGATCC***ctcgagcttagaaaggattattg-3' | | |
| JW-1255-a-Fwd | 5'-ggcatggatgaactatacaaatag***ACTAGT***gtcgatcacggaggaacg-3' | rbs-*McaA* | pAK1255 |
| JW-1255-a-Rev | 5'-gtaccgatttctgc***GGATCC***ctttttgaaaatggcattcc-3' | | |
| JW-1255-b-Fwd | 5'-cattttcaaaaag***GGATCC***gcagaaatcggtactggctttc-3' | *halo* | |
| JW-1255-b-Rev | 5'-caccgcggtggcggccgctctaga***ACTAGT***ctagccggaaatctcgag-3' | | |
| JW-1256-a-Fwd | 5'-gggccccccctcgaggtcgacggtatcgat***AAGCTT***gtgcccgcgactaatgcc-3' | Native promoter plus *mcaB* | pAK1256 |
| JW-1256-a-Rev | 5'-ctcctttactcat***GGATCC***ctcgagcttagaaaggattattg-3' | | |
| JW-1256-b-Fwd | 5'-ctttctaagctcgag***GGATCC***atgagtaaaggagaagaac-3' | *gfp* | |
| JW-1256-b-Rev | 5'-cctccgtgatcgac***ACTAGT***ctatttgtatagttcatccatg-3' | | |
| JW-1256-c-Fwd | 5'-ctatacaaatag***ACTAGT***gtcgatcacggaggaacgtag-3' | rbs-*mcaA* | |
| JW-1256-c-Rev | 5'-tggaaagccagtaccgatttctgc***GGATCC***ctttttgaaaatggcattcctgcttttc-3' | | |
| JW-1270-Fwd | 5'-ggcatggatgaactatacaaatag***ACTAGT***tcacacaggaaacagaattc-3' | rbs-*mms6*-*halo* | pAK1270 |
| JW-1270-Rev | 5'-caccgcggtggcggccgctctaga***ACTAGT***ctagccggaaatctcgag-3' | | |

#### Supplementary Table 12. List of primers for making mcaA mutant plasmids

| Name | Sequence | Target | In plasmid |
|---|---|---|---|
| JW-1-Fwd | 5'-gataacaatttcacacaggaaaca***GAATTC***gtggatgccgacattattgttc-3' | $McaA^{\Delta SP}$ | pAK1257 |
| JW-1-Rev | 5'-gaaaagttcttctcctttactcat***GGATCC***ctttttgaaaatggcattc-3' | | |
| JW-2a-Fwd | 5'-gataacaatttcacacaggaaaca***GAATTC***gtggtaattcgaggggggg-3' | Upstream-$McaA^{\Delta VWA}$ | pAK1258 |
| JW-2a-Rev | 5'-ccatggccgtcaattggcgtgcattcgcacttc-3' | | |
| JW-2b-Fwd | 5'-gaatgcacgccaattgacggccatgggggtacttc-3' | Downstream-$McaA^{\Delta VWA}$ | |
| JW-2b-Rev | 5'-gaaaagttcttctcctttactcat***GGATCC***ctttttgaaaatggcattc-3' | | |
| JW-3a-Fwd | 5'-gataacaatttcacacaggaaaca***GAATTC***gtggtaattcgaggggggg-3' | Upstream-$McaA^{MIDAS}$ | pAK1259 |
| JW-3a-Rev | 5'-cagctcctgccttagcaaaaagaacaataatgtc-3' | | |
| JW-3b-Fwd | 5'-gctaaggcaggagctgtaaataggcacgacccaaag-3' | Downstream-$McaA^{MIDAS}$ | |
| JW-3b-Rev | 5'-gaaaagttcttctcctttactcat***GGATCC***ctttttgaaaatggcattc-3' | | |
| JW-4a-Fwd | 5'-gataacaatttcacacaggaaaca***GAATTC***gtggtaattcgaggggggg-3' | Upstream-$McaA^{\Delta aa259\text{-}400}$ | pAK1260 |
| JW-4a-Rev | 5'-cctgagccgccttcagctggaccgaaggattctc-3' | | |
| JW-4b-Fwd | 5'-cttcggtccagctgaaggcggctcaggcggcacg-3' | Downstream-$McaA^{\Delta aa259\text{-}400}$ | |
| JW-4b-Rev | 5'-gaaaagttcttctcctttactcat***GGATCC***ctttttgaaaatggcattc-3' | | |
| JW-5a-Fwd | 5'-gataacaatttcacacaggaaaca***GAATTC***gtggtaattcgaggggggg-3' | Upstream-$McaA^{\Delta aa400\text{-}530}$ | pAK1261 |
| JW-5a-Rev | 5'-caatgaattcagcaacttgtccagccagcttacgacg-3' | | |
| JW-5b-Fwd | 5'-gctggctggacaagttgctgaattcattgatggg-3' | Downstream-$McaA^{\Delta aa400\text{-}530}$ | |
| JW-5b-Rev | 5'-gaaaagttcttctcctttactcat***GGATCC***ctttttgaaaatggcattc-3' | | |
| JW-6a-Fwd | 5'-gataacaatttcacacaggaaaca***GAATTC***gtggtaattcgaggggggg-3' | Upstream-$McaA^{\Delta aa530\text{-}665}$ | pAK1262 |
| JW-6a-Rev | 5'-cagttcctttcgccatgtggtcggagagtgagac-3' | | |
| JW-6b-Fwd | 5'-ctccgaccacatggcgaaaggaactgagcacgatg-3' | Downstream-$McaA^{\Delta aa530\text{-}665}$ | |
| JW-6b-Rev | 5'-gaaaagttcttctcctttactcat***GGATCC***ctttttgaaaatggcattc-3' | | |
| JW-7-Fwd | 5'-gataacaatttcacacaggaaaca***GAATTC***gtggtaattcgaggggggg-3' | 7-$McaA^{\Delta aa665\text{-}776}$ | pAK1263 |
| JW-7-Rev | 5'-gaaaagttcttctcctttactcat***GGATCC***aattctgctaacaacctcgatc-3' | | |
